# Bacterial motility governs the evolution of antibiotic resistance in spatially heterogeneous environments

**DOI:** 10.1101/2022.10.21.513270

**Authors:** Vit Piskovsky, Nuno M. Oliveira

## Abstract

Bacteria evolving in natural and clinical settings experience spatial fluctuations of multiple factors and this heterogeneity is expected to affect bacterial adaptation. Notably, spatial heterogeneity in antibiotic concentrations is believed to accelerate the evolution of antibiotic resistance. However, current literature overlooks the role of cell motility, which is key for bacterial survival and reproduction. Here, we consider a quantitative model for bacterial evolution in antibiotic gradients, where bacteria evolve under the stochastic processes of proliferation, death, mutation and migration. Numerical and analytical results show that cell motility has major effects on bacterial adaptation. If migration is relatively rare, it accelerates adaptation because resistant mutants can colonize neighbouring patches of increasing antibiotic concentration avoiding competition with wild-type cells; but if migration is common throughout the lifespan of bacteria, it decelerates adaptation by promoting genotypic mixing and ecological competition. If migration is sufficiently high, it can limit bacterial survival, and we derive conditions for such a regime. Similar patterns are observed in more complex scenarios, namely where bacteria can bias their motion or switch between motility phenotypes either stochastically or in a density-dependent manner. Overall, our work reveals limits to bacterial adaptation in antibiotic landscapes that are set by cell motility.

---

Active motility is a defining feature of many cell types, governing their ecology and physiological functions. Immune cells such as neutrophils patrol multicellular organisms in their relentless search for invading microbes. Sperm cells actively swim and search ovules to fuse with. Growth cones of neurons seek their synaptic targets. Active motility is also fundamental for many unicellular organisms ranging from bacteria to amoeba and algae. It allows them to find nutrients, light or a host, and avoid toxic compounds, predators or parasites. In particular, bacterial motility has been thoroughly studied once it became understood as key for the reproductive success of bacteria and, more specifically, for their ability to cause disease [1–7]. Yet, despite these important realizations, there is surprisingly little evidence about how motility contributes to bacterial adaptation, namely to the evolution of antibiotic resistance, which is ranked among other major threats such as climate change and terrorism [8].

Laboratory experiments suggest that bacterial motility is important for bacterial adaptation in antibiotic landscapes where cells can move through different concentrations of antibiotics [9–11]. In particular, it was found that in spatially heterogeneous environments, bacteria evolve antibiotic resistance faster than in homogeneous conditions [9]. However, cell motility was not controlled in these experiments, and its precise contribution to the evolutionary dynamics found is not known. Notably, other authors did not find an accelerated adaptation in their antibiotic landscapes and argued that the discrepancy likely derived from the ability of bacteria to experience the diverse antibiotic concentrations, which was different in their experiments [10].

Arguably, the best understood link between cell motility and the ability of bacteria to cope with antibiotics comes from bacterial swarming, a form of group motility on surfaces where cells are particularly resilient to antibiotic stress [12, 13]. More precisely, in swarming conditions, cell motility is thought to reduce exposure of individual cells to antibiotics, which leads to antibiotic tolerance [14]. While there are many studies relating bacterial swarming and phenotypic resistance [12, 14–21], the effect of swarming motility on the evolution of genetic resistance has not been explored. In addition to swarming, other links between bacterial motility and antibiotic landscapes have been reported. In particular, it has been shown that antibiotics trigger motile responses [22–24]. However, the impact of such bacterial behaviour on the evolution of antibiotic resistance was not addressed. In short, one can find works that study bacterial evolution in antibiotic landscapes, but these do not explore the role of cell motility explicitly; and one can find works that study bacterial motility upon antibiotic exposure, but these do not explore evolutionary timescales.

In addition to these experimental works, several mathematical models have been developed to understand how concentration gradients of antibiotics and other drugs affect resistance evolution [25–31], and a key conclusion is that antibiotic gradients accelerate the evolution of antibiotic resistance [25, 26]. These quantitative models consider bacterial movement between environments with different antibiotic concentrations, such as different organs or parts of the human body, but the migration considered is essentially a passive and rare event. This assumption is in sharp contrast with the migration rates of bacteria evolving in natural and experimental settings, where cells can move at speeds up to ~ 50μm/s, and the fact that in these environments bacteria can experience steep and stable gradients of antibiotics and other drugs [9–11, 22]. Here, we build upon these quantitative models and study how bacterial motility affects the evolution of antibiotic resistance in spatially heterogeneous environments with different concentrations of antibiotics.

## I. RESULTS

### A. Model overview and basic dynamics

We are interested to understand how bacterial motility shapes the evolution of resistance in heterogeneous environments where cells experience spatial gradients of antibiotics. We build upon the so-called staircase model [25], which was developed to understand bacterial evolution in drug gradients. The staircase model is a lattice model, which considers cells of specified genotype *g* ∈ {1,…, *L*} (of increasing resistance) and spatial compartments *x* ∈ {1,…, *L*} (of increasing antibiotic concentration), Fig. 1a. An initial population of susceptible cells *g* = 1 at position *x* = 1 evolves via stochastic processes of death (rate *δ*), movement to a neighbouring spatial compartment (rate v), mutation (rates *μ_f_* /*μ_b_* for forward/backward), or division (maximal rate *r*) [25]. In this model, antibiotics inhibit the growth of susceptible genotypes *g* < *x*. Therefore, the diagonal of position-genotype lattice in Fig. 1a defines a staircase which separates areas where cells can divide (above the staircase, white) from areas where they cannot (below the staircase, grey). Moreover, the division rate of the *N_x_* cells at position *x* is logistic with a carrying capacity *K*:

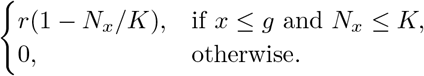

**FIG. 1.**
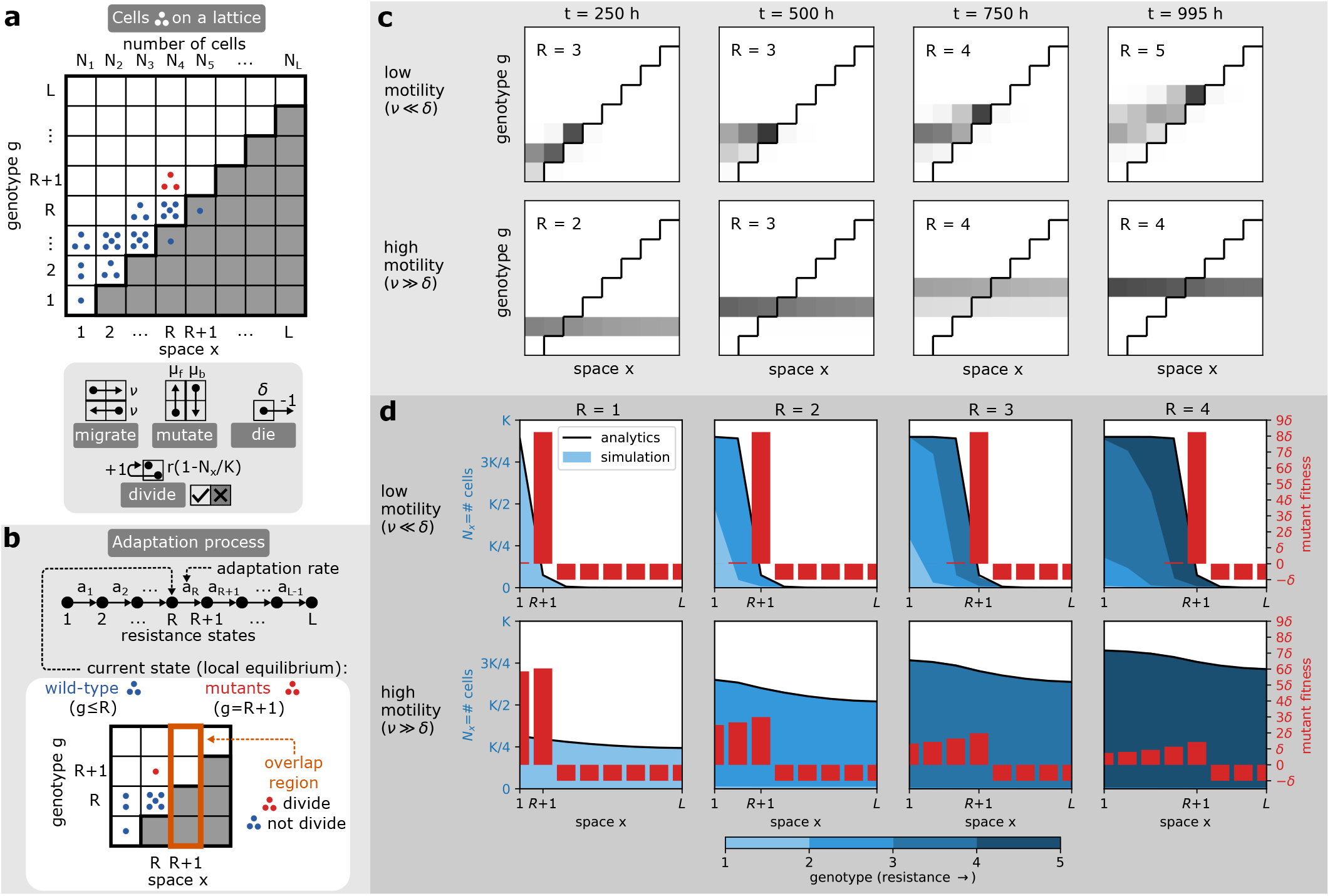
Model overview and basic evolutionary dynamics. **a)** Model description. In the staircase model, bacterial cells (black dots) have a specified position *x* (of increasing antibiotic concentration) and a genotype *g* (of increasing resistance). The population adapts to a stable spatial antibiotic gradient as cells migrate, mutate, die and divide at specified stochastic rates. Antibiotics prevent cell division of susceptible genotypes *g* < *x* (shaded region under the staircase). **b)** Adaptation process. Bacterial adaptation in the staircase model can be conceptualised as a series of random jumps between locally stable resistance states *R* that happen at an adaptation rate *a_R_*. At a given time, *R* denotes the genotype of highest resistance in the population, as well as the highest spatial compartment where this genotype can divide. Genotypes *g* ≤ *R* are defined as wild-type and genotype *g* = *R* +1 as mutant. Importantly, in the overlap region *x* = *R* +1, mutant cells can divide but wild-type cells cannot. **c)** Snapshots of the population evolving antibiotic resistance for different motility rates, *ν*. Shades of grey represent density of cells, where black represents highest density. The adaptive dynamics of bacteria with low (top) and high (bottom) motility is qualitatively different. **d)** Population profiles and fitness along the antibiotic gradient. Different motility regimes affect the distribution of wild-type bacteria along the gradient, which shapes the fitness of resistant mutants.

The processes of birth, death, mutation and migration drive bacterial adaptation up the antibiotic gradient. Assuming low mutation rates [25, 26], this adaptation corresponds to random jumps *R* → *R* +1 between relatively stable population states *R* ∈ {1,…,*L*}, which we define as the resistance states (Fig. 1b). We note, however, that *R* can be understood as both the genotype *g* of highest resistance in the population and the compartment of highest antibiotic concentration *x* where this genotype can divide because the genotypic and spatial dimensions are interchangeable in our model. Genotypes *g* ≤ *R* are defined as wild-type and genotype *g* = *R* + 1 as mutant.

Resistance *R* increases to *R* +1 when mutants outcompete the wild-type in the overlap region at a position *x* = *R* + 1. The overlap region is the region where the population first adapts because mutant cells can divide there but wild-type cells cannot (Fig. 1b).

We start by studying how different motility rates *ν* affect the evolutionary dynamics of bacteria in this model as previous work [25, 26] only considers situations where bacteria are unlikely to move between different antibiotic concentrations during their lifetime (*ν* < *δ*). In particular, we start by describing and comparing bacterial adaptation when cells rarely move during their lifespan (*ν* < *δ*) and situations where motility is common (*ν* > *δ*). Unexpectedly, our simulations show that bacterial adaptation is qualitatively different for low and high motility (Fig. 1c, SI Movie S1, S2). To understand this difference, we first study the qualitative properties of the stable population at a fixed resistance state *R* and in the next section we focus on the adaptation rate *a_R_* at which the jump in resistance *R* → *R* +1 occurs (Fig. 1b).

Notably, we find that the low and high motility regimes differ in the shape of the wild-type population *N_x_* (SI Theorem 1), which impacts the mutant fitness landscape (Fig. 1d) that is defined by:

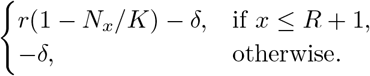

Moreover, we find that at low motility (*ν* < *δ*, SI Movie S1), bacterial adaptation is limited by the movement of the first mutant into the overlap region, where mutant cells have low competition from wild-type cells and high fitness to set up a mutant population (Fig. 1d). In this case, bacterial adaptation is located at the population front (SI Theorem 3), while the rest of the population forms a typical inclined comet tail (Fig. 1c) corresponding to a diversity of strains with different antibiotic susceptibility (Fig. 1d). At high motility (*ν* > *δ*, SI Movie S2), the wild-type is spread across space (SI Theorem 3), competing with mutants for space even in regions where the wild-type cannot grow. The fitness of mutants is decreased in the overlap region and leveled across the region where mutants can divide (Fig. 1d). Therefore, mutant cells grow slowly and outcompete the wild-type everywhere. In these conditions, the population is made of a single strain that can grow in *x* ≤ *R* (Fig. 1d) and the comet tail is now horizontal (Fig. 1c).

### B. Cell motility accelerates and decelerates bacterial adaptive evolution

We have shown that bacterial adaptation in antibiotic gradients is qualitatively different at low and high motility regimes. To gain quantitative understanding about this difference, we now study how motility affects the adaptation rate *a_R_* (Fig. 1b, 2a), which is defined by two waiting times,

1. The evolutionary time *T_evo_*, defined as the earliest time when the wild-type produces a first mutant in the overlap region (Fig. 2a).
2. The ecological time *T_eco_*, defined as the earliest time when the first mutant establishes a mutant population that becomes larger than the wild-type in the overlap region (Fig. 2a).

**FIG. 2.**
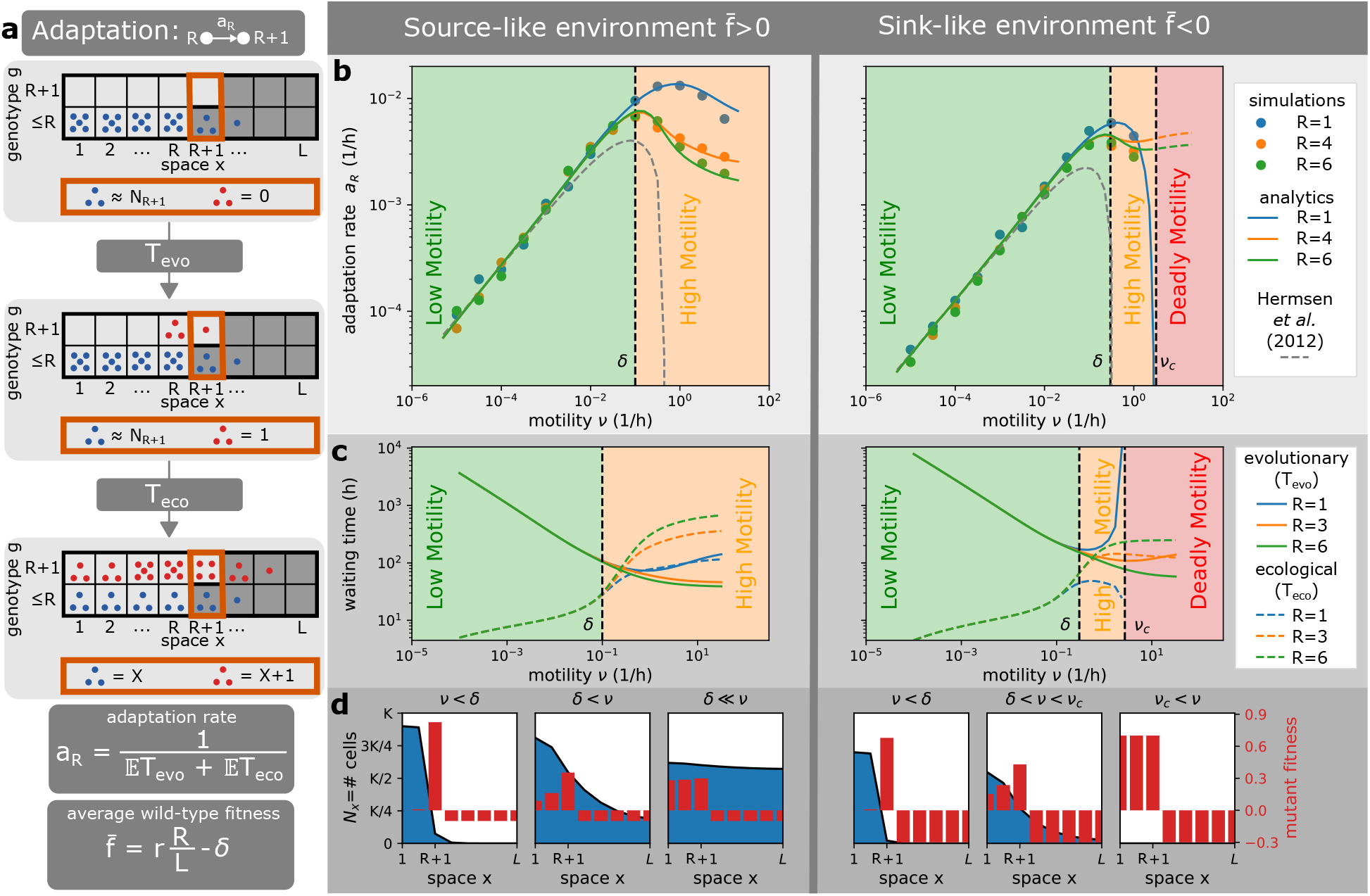
Motility can accelerate and decelerate the rate of bacterial adaptation to antibiotics. **a)** Definitions. The adaptation process in the staircase model corresponds to jumps between resistance states *R* → *R* + 1 (Fig. 1b) that happen at the rate *a_R_*. This adaptation rate is defined by two waiting times, the evolutionary time *T_evo_* (the time it takes for a first mutant to appear in the overlap region) and the ecological time *T_eco_* (the time it takes for mutants to have larger population than the wild-type in the overlap region). The average wild-type fitness is defined as 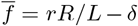 with *r* being the birth rate, *δ* death rate, and *R/L* the proportion of space where wild-type cells can divide. **b)** Adaptation rate as a function of motility rate. Motility accelerates adaptation in the low motility regime, but decelerates adaptation in the high motility regime. If the wild-type has a negative average fitness and motility above a critical motility *ν_c_*, it cannot survive and adapt (deadly motility regime). The current resistance state *R* does not affect the adaptation rate when motility is low, but decreases the adaptation rate when motility is high. We plot the predicted dynamics from the analytical theory of [26] (dashed line), which matches our results for lower motility only, but note that this theory was developed under the assumption of low motility rates and the interpolation for higher motility shown for completeness is not valid. **c)** Relative importance of evolutionary and ecological times. At low motility, bacterial adaptation is largely dependent on the evolutionary time *T_evo_*. At high motility, however, the ecological time *T_eco_* becomes equally important due to increased competition between strains. Near critical motility *ν_c_*, bacterial adaptation is again mainly limited by the evolutionary time *T_evo_*. At high motility, the number of compartments where the wild-type can divide *R* increases the ecological time *T_eco_* and decreases the evolutionary time *T_evo_*. **d)** The importance of wild-type profiles. At low motility regimes, motility promotes the movement of mutants from the population front into the overlap region (*T_evo_* reduced), where mutants have larger fitness than the wild-type and grow quickly (*T_evo_* ≫ *T_eco_*). At high motility regimes, motility promotes the spatial spread of the wild-type population with genotypes *g* ≤ *R*, so that the first mutant is produced quickly but mutant fitness is decreased by increased competition with the wild-type (*T_evo_* < *T_eco_*).

These waiting times are random variables and their expected value contribute to the adaptation rate as follows:

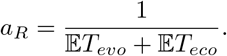

To determine waiting times and adaptation rates, we used computer simulations (Fig. 2b), and a combination of analytical and numerical techniques that confirm and extend our simulations (Fig. 2b, Fig. 2c and Methods).

How does the adaptation rate *a_R_* vary in the two motility regimes we identified earlier? Our simulations show that low motility (*ν* < *δ*) accelerates the adaptation rate of bacteria (Fig. 2b), in accordance with previous theory [25, 26]. Our analysis shows that in these conditions the evolutionary time is much bigger than the ecological time (Fig. 2c), because mutants have high fitness in the overlap region and can grow quickly (Fig. 2d). For this reason, the ecological time has been neglected in previous works [26] and the adaptation rate has been defined by the evolutionary time alone as 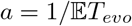, irrespective of the resistance state *R*. Indeed, [26] that considered low motility only noticed that the rate at which the population front advances becomes constant (i.e., it is independent of *R*) and our work supports this claim as the adaptation rate *a_R_* is identical for all *R* when motility is low (Fig. 2b) and the shape of the population front is independent of *R* (Fig. 1d, SI Theorem 3). Under these conditions, motility helps the first mutant to move from the population front into the overlap region and decreases the dominant waiting time, the evolutionary time *T_evo_*. Therefore, in this low motility regime, cell motility accelerates bacterial adaptation. In contrast, high motility (*ν* > *δ*) decelerates adaptation rate (Fig. 2b). In this regime, the ecological time becomes bigger than the evolutionary time (Fig. 2c) because mutant fitness is significantly decreased, which results from the increased competition for space with a large wild-type population in the overlap region. This competition is amplified with increasing motility *ν* and resistance state R, which promote the spread and size of the population (Fig. 2c).

The reduction in adaptation rate with higher motility becomes even more important if the wild-type has negative spatially-averaged fitness, which is defined as 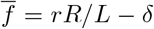 with *r* being the birth rate, *δ* death rate, and *R/L* the proportion of space where wild-type cells can divide. By definition, environments with this property, 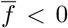, cannot sustain the wild-type if the system is spatially homogeneous, and we call them sink-like (as opposed to source-like with 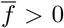). This terminology is inspired by the source-sink theory, which defines a source (resp. sink) as an environment with positive (resp. negative) local fitness [32], as opposed to the average fitness in our model that considers fitness across all compartments. Notably, we find that cell motility creates sufficient mixing in sink-like environments above a critical motility threshold (*ν* > *ν_c_*), and that in these conditions the wild-type cannot survive and adapt (Fig. 2b, SI Theorem 1, SI Theorem 4, SI Corollary 1). Therefore, we call it a deadly motility regime. We note that the deadly motility regime would not occur if resistant cells were present in the population from the beginning, which would increase the average wild-type fitness from 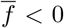 to 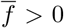 due to higher *R*. In this case, the same environment would become source-like, with survival and adaptation possible for any motility (dashed lines in Fig. 2b). We further note that the existence of the deadly motility regime requires the system to be closed, with no external import of wild-type cells.

In summary, our analysis identifies three distinct regimes of bacterial adaptation in antibiotic gradients that depend on the motility rate of cells, which we call low motility, high motility and deadly motility regimes (Fig. 2a). Notably, we find the same regimes when antibiotic resistance carries fitness costs (Fig. S1, SI Text S4), and when instead of bacteriostatic antibiotics we consider bactericidal antibiotics (Fig. S2, SI Text S4). Unexpectedly, we find the same regimes even when cell motility is biased (Fig. S3, SI Text S4, SI Movies S3 and S4). It has recently been shown that antibiotics can trigger both positive [22] and negative [24] chemotaxis in bacterial cells, but our results show that such biased motility does affect our key findings (Fig. S3).

### C. Evolutionary dynamics of phenotypically heterogeneous populations

So far, we have considered evolving populations that are phenotypically homogenous and bacterial cells always move at the same rate. The same simplifying assumption has been made in previous models of bacterial evolution in antibiotic gradients [25–31]. However, in reality, bacteria can change their motility rate, and their communities are phenotypically heterogeneous in terms of cell motility. For example, one of the most important transitions in a bacterial life-cycle from free-living (plankton) to surface-attached communities (biofilms) is associated with major changes in cell motility. Planktonic bacteria can move at 45 *μm/s* [2], while biofilm bacteria can move at 1 *μm/min* or less [33]. The plankton-biofilm switch can occur both stochastically [34] or as a response to multiple factors, such as cell density [35] or the presence of other strains [36]. Another well-known transition in bacterial populations that is associated with changes in cell movement is the swarming phenotype, where bacteria move collectively on semi-solid surfaces when a threshold of cell density is reached [37]. We found it important to to understand if this phenotypic heterogeneity could affect the evolutionary dynamics we presented earlier.

To address this issue, we now consider populations harbouring two motility phenotypes where cells can switch between phenotypes either stochastically (Fig. 3) or as a response to local cell density (Fig. 4). In practice, for stochastic switching, we add a new dimension of motility phenotypes to our position-genotype lattice and allow cells to switch between phenotypes at a stochastic rate *s*, where each phenotype is characterized by different motility rates, *ν_1_* and *ν_2_* (Fig. 3a). Density-dependent motility *ν*(*N_x_*) is modeled differently, and the switch between motility phenotypes is modeled by a step-function that is controlled by the local number of cells *Nx* and a switching density threshold *S* (Fig. 4a). If cell density *N_x_* at position *x* is below (resp. above) the switching threshold *S*, all cells at position *x* move at low-density (resp. high-density) motility *ν_L_* (resp. *ν_H_*).

**FIG. 3.**
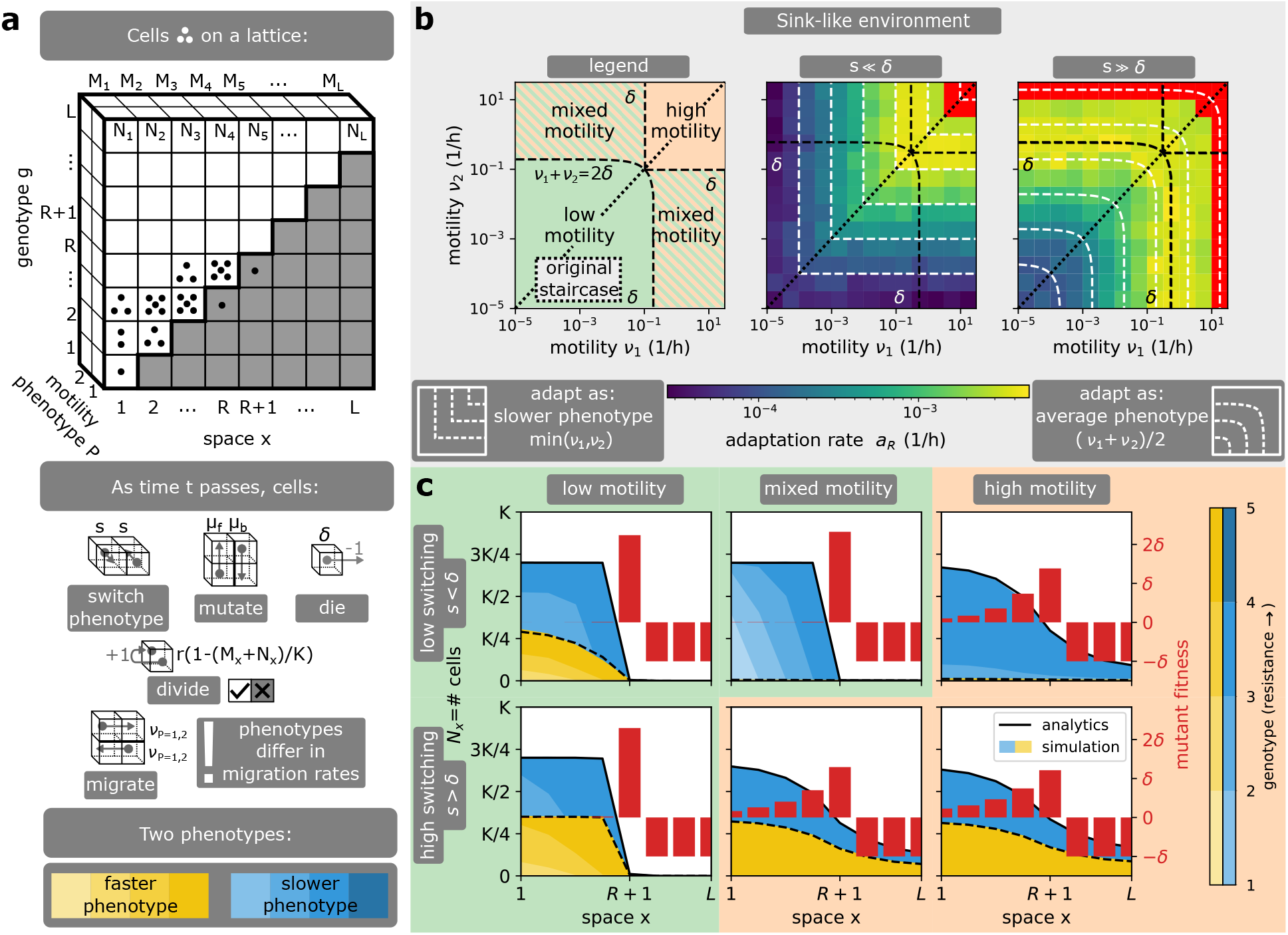
Effect of switching between motility phenotypes stochastically. **a)** Model description. Phenotypes of motility *ν_1,2_* are added as an extra dimension to the staircase model, with individual cells allowed to switch between these phenotypes at rate *s*. **b)** Adaptation rate heatmap on the (*ν_1_, ν_2_*) plane for different switching rates *s*. The (*ν_1_, ν_2_*) plane can be partitioned into different combinations of adaptation regimes and its diagonal corresponds to a population of a single motility. In this model, bacteria adapt as if all cells had the same effective motility, which corresponds to the intersection of the level sets (white dashed lines) with the diagonal. At low switching rate s, the effective motility matches the slower motility present in the population min(*ν_1_,ν_2_*). At high switching rate s, the effective motility matches the average motility (*ν_1_ + ν_2_*)/2. When the environment is sink-like, a deadly motility regime (red) exists in the high motility combination, and also appears in the mixed motility combination when the switching rate is high. **c)** Wild-type profiles for different switching rates *s*. At low switching rate *s*, the wild-type profile is dominated by the slower phenotype. At high switching rate s, the wild-type profile coincides with the profile of a single average-motility phenotype. Therefore, the adaptation in low (resp. high) motility combinations follows the low (resp. high) motility regime irrespective of switching rate *s*, but a change of *s* in the mixed motility combination can change the adaptation regime. In short, stochastic motility switching shapes the evolution of antibiotic resistance by determining the effective motility of bacterial populations.

**FIG. 4.**
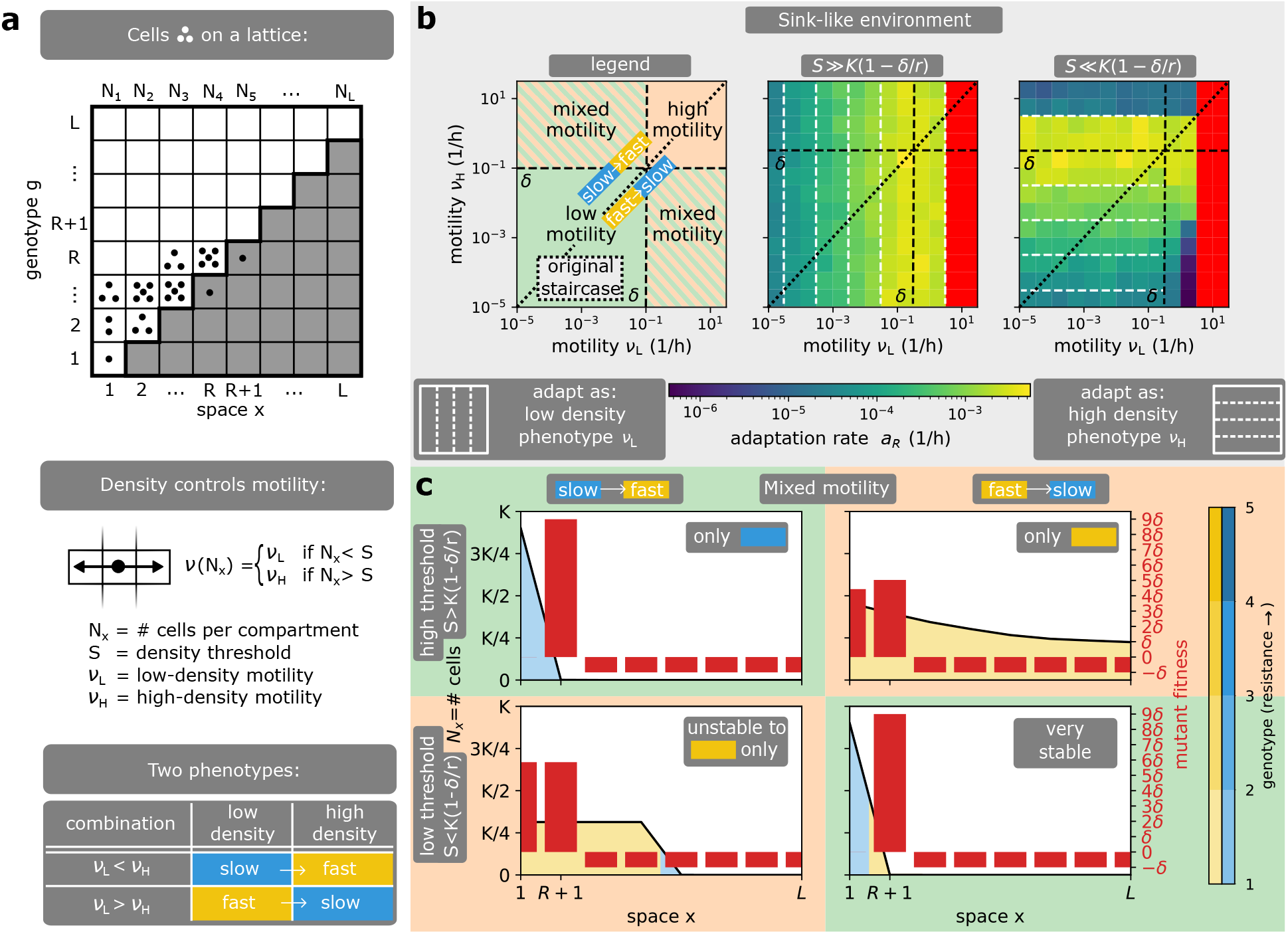
Effect of density-dependent motility. **a)** Model description. In this model, bacterial motility *ν*(*N_x_*) depends on the local number of cells *N_x_*. If the cell density *N_x_* at position *x* is below (resp. above) the switching threshold *S*, all cells at position *x* move at low-density (resp. high-density) motility (*ν_H_*). **b)** Adaptation rate heatmap on the (*ν_L_,v_H_*) plane for different switching thresholds *S*. The (*ν_L_, v_H_*) plane can be partitioned into different combinations of adaptation regimes and its diagonal corresponds to a population of a single motility. The diagonal separates slow-to-fast and fast-to-slow switching combinations. Bacteria generically adapt as if all cells had the same effective motility, which corresponds to the intersection of the level sets (white dashed lines) with the diagonal. This effective motility generically matches the low-density (resp. high-density) motility at high (resp. low) threshold *S*. At low threshold *S*, this generic rule has three exceptions. First, the adaptation rate is reduced in the mixed motility combination of fast-to-slow switching. Second, the adaptation rate is reduced for slow-to-fast switching when the environment is sink-like and the high-density motility phenotype moves above the critical motility. Third, a deadly motility regime can only occur if the initial population is of low density and the low-density motility phenotype moves above the critical motility. **c)** Wild-type profiles in the mixed motility combination. Wild-type profiles match the low-density motility phenotype at high threshold *S*. At low threshold *S*, the low-density motility phenotype appears only at the front while the bulk of the population is in the high-density motility phenotype. This composition of phenotypes can reduce the antibiotic exposure of the bacterial population: in slow-to-fast switching, the spatial expansion of the fast (yellow) high-density motility phenotype is reduced in sink-like environments; while in fast-to-slow switching, a very slow (blue) and stable population occupies low antibiotic concentrations. In short, density-dependent motility shapes bacterial adaptation by determining the effective motility of bacterial populations and affecting their antibiotic exposure.

Interestingly, we find that in both models, phenotypically heterogeneous populations of bacteria generically behave as if all cells had a single effective motility phenotype whose motility is controlled by the switching rate *s* in stochastic switching, or the switching threshold *S* if switching is density-dependent. Therefore, the previously described adaptation regimes exist in these models and are governed by an effective motility that we characterize next. For stochastic switching, the effective motility matches the motility of the slower phenotype *ν*_ = min(*ν_1_,v_2_*) when the switching rate is low (*s* ≪ *δ*, SI Movie S5) and it matches the average motility *ν*_+_ = (*ν_1_* + *ν_2_*)/2 when the switching rate is high (*s* ≫ *δ*, SI Movie S6), see Fig. 3b and SI Theorems 6, 7, 8 and SI Corollary 2, 3. This result stems from the fact that the slower phenotype is naturally selected for in the absence of phenotypic switching, given that the slower phenotype is exposed to antibiotics less often and maximises the local fitness of the wild-type, and stochastic switching levels out the numbers of both phenotypes (Fig. 3c). For density-dependent motility, the effective motility matches the low-density motility phenotype *ν*_ = *ν_L_* at high switching threshold (*S* > *K*(1 – *δ*/*r*)) and the high-density motility phenotype *ν*_+_ = *ν_H_* at low switching thresholds (*S* < *K*(1 – *δ*/*r*)), see Fig. 4b. This result stems from the fact that the population cannot reach a sufficient density for the high-density motility phenotype when the threshold is high, while a lower threshold switches the dense bulk of the population into the high-density motility phenotype.

Accordingly, if the population harbours phenotypic heterogeneity of the same motility regime, the adaptation regime corresponding to the effective motility is not changed by the stochastic rate *s* and the density threshold *S*. However, for populations that harbour motility phenotypes from different motility regimes (mixed motility in Fig. 3b, Fig. 4b), the level of switching shapes the adaptation regime of bacteria (Fig. 3c, Fig. 4c). For stochastic switching, such heterogeneous populations have an effective motility in the low motility regime at low switching (*ν*_ < *δ*) and in the high (resp. deadly) motility regime at high switching (*ν*_+_ > *δ*, resp. *ν*_+_ > *ν_c_*). Put differently, a gradual increase in switching rate changes the adaptation regime between the low, high and deadly motility regimes (Fig. 3c, S4a, SI Theorems 6, 8). Similarly to phenotypically homogeneous populations, the deadly motility regime can only occur if the average wild-type fitness is negative (sink-like environment, Fig. 3b, S4c, SI Theorem 8, SI Corollary 3).

For highly heterogenous populations with densitydependent motility, the gradual change in switching threshold *S* also dictates the effective motility and the corresponding adaptation regime, but there are three exceptions when the density threshold *S* is low (Fig. 4b). First, when cells switch from fast to slow above the threshold *S*, the population becomes very stable as the fast cells at the front quickly return to the slow population bulk (Fig. 4c). As a result, the population has decreased antibiotic exposure and adaptation rate (Fig. 4b). Second, a deadly motility regime occurs if the low-density motility (not the effective motility) is above the critical motility (Fig. 4b). Third, if the high-density motility is above the critical motility, the population does not go extinct but instead decreases its range and antibiotic exposure (Fig. 4c, SI Movie S7), leading to a decreased adaptation rate (Fig. 4b). Since the last two exceptions are related to critical motility, they are only important in sink-like environments where the wild-type has negative average fitness (Fig. S5).

With the exception of the highly heterogeneous populations with density-dependent motility, we showed that bacterial populations with different motility phenotypes evolve antibiotic resistance as if all cells had the same effective motility. Therefore, in these conditions, bacterial adaptation is characterised by the same adaptation regimes of phenotypically homogeneous populations. Switching between motility phenotypes can shape the adaptation regime under which bacteria evolve only if the motility of the different phenotypes is very different. But even in these cases, we find that our general conclusions hold, low effective motility accelerates bacterial adaptation while high effective motility decelerates it. For very high effective motility, bacterial populations can perish.

### D. Cell motility as a factor that limits bacterial survival in sink-like environments

We have seen how cell motility shapes the evolution of antibiotic resistance in a variety of models. We have shown that the wild-type profiles form the basis of bacterial adaptation by creating mutants and dictating their fitness in antibiotic landscapes (Fig. 1d). Moreover, all our models admit a deadly motility regime, defined by a critical motility (Fig. 2b, Fig. 3b, Fig. 4b). Above the critical motility, the wild-type goes extinct and adaptation is not possible. The goal of this last section is to understand the emergence of such critical motility regime.

We start by studying analytically how the wild-type profile forms at a fixed resistance state *R* (Fig. 1b, 1d). A wild-type profile (*N*_1_,…, *N_L_*) can be conceptualised as a point in the profile space of cell numbers (Fig. 5a) and its time-evolution can be described by the mean-field equations that are derived from the stochastic dynamics. We illustrate the general approach (SI Theorems 2, 3) using the staircase model with only two compartments (*L* = 2, *R* =1), which is known as source-sink model [25]. In this model, the mean-field equations for the wild-type profile formation are the following

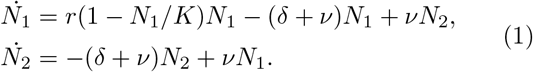

**FIG. 5.**
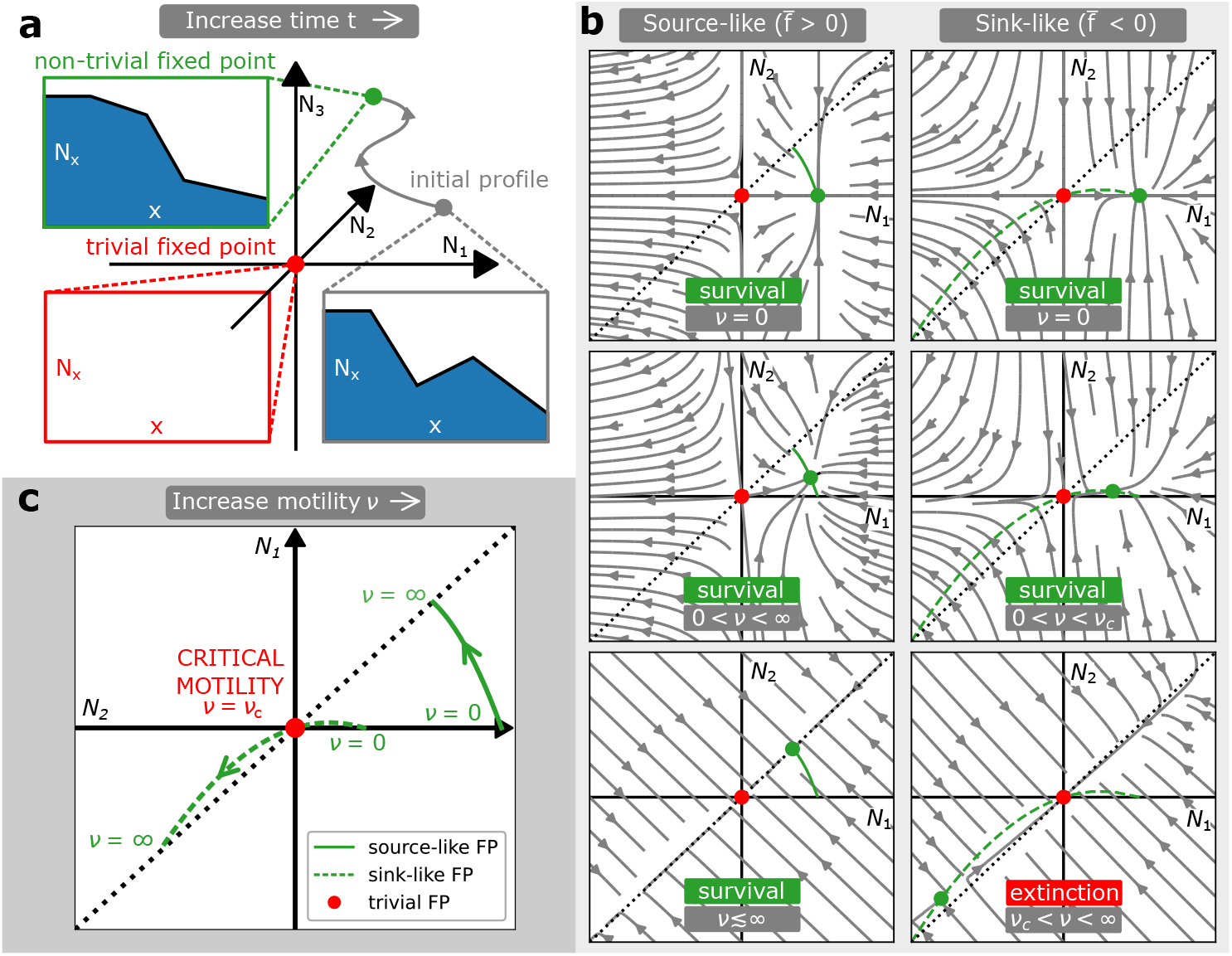
Cell motility as a factor that limits bacterial survival in sink-like environments. **a)** The formation of the wild-type profile. The formation of a wild-type profile corresponding to a fixed resistance state *R* (Fig. 1b) is described by mean-field equations, which explain the flow of the initial profile in the profile space of cell numbers (*N*_1_, …, *N_L_*). This flow admits two fixed points: a trivial fixed point *N_x_* = 0 (red, population extinction if stable) and a non-trivial fixed-point (green, population survival if stable). **b)** Phase portraits of the wild-type profile formation in the staircase model with *L* = 2, *R* =1 (the so-called source-sink model [25]). Phase portraits are shown for different motility rates *ν* and different environment types (source-like/sink-like), depending on the average wild-type fitness 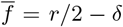. Only two topologically distinct types of flow are possible: the non-trivial fixed point globally attracts all possible wild-type profiles with *N_x_* > 0 (population survival), or the trivial fixed point globally attracts all possible wild-type profiles with *N_x_* > 0 (population extinction). Extinction occurs in sink-like environments 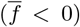 with motility above the critical motility *ν* > *ν_c_*. **c)** Critical motility *ν_c_* corresponds to a bifurcation of this dynamical system. When motility *ν* is varied, the non-trivial fixed point moves through the space of possible profiles. At low motility, the non-trivial fixed point is stable and corresponds to a wild-type profile that predominantly occupies spatial positions *x* ≤ *R* where it can divide. As motility increases, the wild-type profile gets increasingly levelled across all spatial compartments, and its dynamics becomes governed by the average fitness 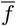. Precisely when the environment is sink-like 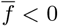, the stable non-trivial fixed point (surviving wild-type) collides with the unstable trivial fixed point (extinct wild-type), and they exchange their stability at the critical motility *ν_c_*. In short, highly motile populations experience an average environment, which can drive their extinction if the environment is sink-like.

This system can be analysed with dynamical systems theory, which studies qualitative features of systems of ordinary differential equations (ODE). More precisely, the time-evolution of this system on ℝ^2^ can be characterised by phase portraits (Fig. 5b), which in this case have two fixed-points: a trivial fixed point *N*_1_ = *N*_2_ = 0 where the population goes extinct, and a non-trivial fixed-point where the population survives, which is defined as

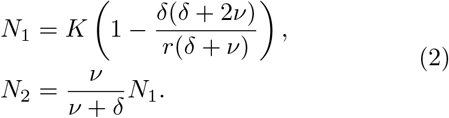

These phase portraits admit only two distinct types of trajectory flows (Fig. 5b). Either the non-trivial fixed point has positive physical densities *N_x_* > 0 and globally attracts all possible wild-type profiles with *N_x_* > 0 (i.e., the bacterial population survives); or the non-trivial fixed point is non-physical *N_x_* < 0 and the trivial fixed point attracts all possible wild-type profiles with *N_x_* > 0 (i.e., the bacterial population does not survive). These two types of phase portraits appear at different motility rates *ν*. Furthermore, we can understand this dynamics with bifurcation theory, and study how the non-trivial fixed point moves through the profile space as motility *ν* increases from zero (Fig. 5c). At low motility, the non-trivial fixed point is stable and located in the nonnegative quadrant *N*_1,2_ ≥ 0. As motility increases, the dynamics is different in source-like 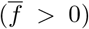 and sinklike 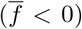 environments, where the average wild-type fitness of the source-sink model is 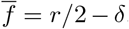. If the environment is source-like, the non-trivial fixed point never leaves the quadrant *N*_1,2_ > 0 and remains stable. However, in sink-like environment, the non-trivial fixed point collides with the trivial fixed point through a transcritical bifurcation, which exchanges their stability. In these cases, there is a critical motility,

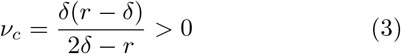

which corresponds to this bifurcation. The emergence of a bifurcation is necessary, since a) the trivial fixed point must be stable as *ν* → ∞ when cells experience an average sink-like environment, and b) the non-trivial fixed point must be stable as *ν* → 0 when cells grow locally in the source at *x* =1 (SI Theorem 3). Therefore, these fixed points must collide and exchange their stability through a transcritical bifurcation. This bifurcation theory argument is general and provides a robust mechanism for the emergence of the critical motility regime in all our models (SI Text S2).

In short, and put differently in a more biological realm, a bacterial population can be driven to extinction by increasing the rate at which cells move in sink-like environments (i.e., in spatially heterogeneous environments where the average wild-type fitness is negative) because highly motile cells experience an average environment.

## II. DISCUSSION

In this work, we have used mathematical analysis to study how bacterial motility affects the evolution of antibiotic resistance in spatially heterogeneous environments that have different antibiotic concentrations. By doing so, our study contributes to the large body of literature that has recognized spatial heterogeneity as an important factor for bacterial adaptive evolution but where cell motility has largely been overlooked [9–11, 25–31].

We have identified three regimes of bacterial adaptation, which are defined by the degree of cell motility in evolving populations. Accordingly, we called them low motility, high motility and deadly motility regimes (Fig. 2b). The theory of the low motility regime has already been discussed [25, 26], and has been compared with experimental works that studied how antibiotic gradients affect bacterial adaptation rate [9, 10]. Interestingly, while [9] reported an increased adaptation rate in their antibiotic gradient compared to well-mixed conditions, [10] did not find such an acceleration in theirs. We note that while the same bacterial species was used, the antibiotic landscape that cells experience is very different in these two experimental systems, which would be sufficient to affect the adaptation rates according to our work (Fig. 2b). Such difference can emerge when cells move at different rates (Fig. 2b), or for different gradient steepness [28, 31]. Indeed, these effects are interchangeable because increasing gradient steepness brings spatial points of fixed antibiotic concentrations closer, which resembles the effect of increasing effective motility.

As a corollary of our analysis, we find that there is an optimal level of cell motility for bacterial adaptation in spatially heterogeneous environments (Fig. 2b). Similar optimum has been identified in models of HIV [38] and cancer [39] resistance evolution. These models are analogous to our setting in that they consider the movement of viruses or cancer cells between compartments of different drug concentrations. However, the motion of viruses considered in the HIV model is passive as opposed to the active movement of bacteria we consider, and the deadly motility regime was not identified in either of the models. It is not clear if these models do not allow for a deadly motility regime or if instead the authors simply did not explore their model fully. Since our models rest on general features of living cells, they may be useful to understand the evolutionary dynamics of cell types other than bacteria, namely cancer cells during chemotherapy, where the role of cell motility and drug gradients remains poorly explored [40, 41].

Some of our findings are closely related to those from ecology works that study biological adaptation at range edges where species expand their range in an environmental gradient by dispersal and mutation [42–45]. In this literature, local adaptation is known to be prevented by high motility, the so-called motility load [43], where less fit wild-type genotypes migrate enough to increase competition with mutant genotypes, which decreases selection for the latter. As a result, genetic diversity is lower in their high motility regime when compared to the low motility regime, as we find in our work (Fig. 1d). This loss of genetic diversity in the ecological models is often associated with critical motility [43, 46, 47], which can be compared with our critical motility. Notably, the emergence of a critical motility in these works can be explained by the same bifurcation mechanism that we identified (SI Text S2), which highlights the robustness of our mechanism across modelling frameworks. Moreover, due to its simplicity, the staircase model can be used to provide analytical insights into biological adaptation at range edges as in [42, 43, 48], while considering the important stochasticity of natural processes as in [45].

Our models make multiple simplifying assumptions regarding the biology of bacteria. While we extended our initial staircase model to account for some of this biology, such as the effect of resistance costs (Fig. S1), bactericidal antibiotics (Fig. S2) and biased motility (Fig. S3), these processes could be modelled differently. For example, we modelled resistance costs as a reduction in division rate (Fig. S1), while one could study resistance costs that decrease motility rate. Moreover, we assumed implicitly that bacterial adaptation results from the vertical transmission of resistance genes only, and horizontal gene transfer (HGT) is known to affect the evolution of antibiotic resistance [49]. However, the staircase model is a closed system and it is known that for these systems, HGT does not have an important impact [50–54]. While considering the effect of immigration of cells was beyond the scope of this paper, we confirmed the predictions for closed systems by implementing HGT in the original staircase model. As predicted, we find that HGT does not affect our conclusions (Fig. S6, SI Text S4).

In addition to considering the effect of HGT more thoroughly, one may extend our models to account for other relevant bacterial biology such as the existence of persister phenotypes, which can affect the stability of microbial communities [55], or the effect of phage infection, which can affect bacterial motility [24]. Regardless, our current results lead us to conclude that cell motility limits bacterial adaptive evolution in spatially heterogeneous environments. This realization then suggests that manipulating motility may prove a useful approach for controlling bacterial systems and their impacts on us.

## III. METHODS

### A. Simulations

A major component of our methods is computer simulations of the staircase model and its extensions. Hereby, we describe how to simulate the basic staircase model, and note that the same algorithm is used to simulate all the remaining models. The staircase model treats bacterial evolution in drug gradients as a continuous-time Markov process, which is a stochastic process whose future state depends only on the present state and not on the past states. A state of the staircase model is specified by the number of bacterial cells *N_x,g_* at each lattice point (*x, g*) ∈ {1, …, *L*}^2^ of the position-genotype lattice. The initial state of the staircase model at time *t* = 0 is chosen as *N*_1,1_ = 10^-3^*K* and *N_x,g_* = 0 for other lattice points (*x,g*) = (1,1). The time-evolution of the initial state of the staircase model *N_x,g_* is determined by the following processes *i* ∈ *I*:

- death of a cell of genotype g at position *x* (*N_x,g_* → *N_x,g_* – 1): rate *b_i_* = *δN_x,g_*,
- division of a cell of genotype *g* at position *x* ≤ *g* when ∑_*g*_ *N_x,g_* ≤ *K* (*N_x,g_* → *N_x,g_* + 1): rate *b_i_* = *r*(1 – Σ_*g*_ *N_x,g_*),
- movement of a cell of genotype *g* from position *x* to position *x* + 1 (*N_x,g_* → *N_x,g_* – 1, *N*_*x*+1,*g*_ → *N*_*x*+1,*g*_ + 1): rate *b_i_* = *νN_x,g_*,
- movement of a cell of genotype *g* from position *x* to position *x* – 1 (*N_x,g_* → *N_x,g_* – 1, *N*_*x*–1,*g*_ → *N*_*x*–1,*g*_ + 1): rate *b_i_* = *νN_x,g_*,
- mutation of a cell at position *x* from genotype *g* to genotype *g* +1 (*N_x,g_* → *N_x,g_* – 1, *N*_*x,g*+1_ → *N*_*x,g*+1_ + 1): rate *b_i_ = μ_f_N_x,g_*,
- mutation of a cell at position *x* from genotype *g* to genotype *g* – 1 (*N_x,g_* → *N_x,g_* – 1, *N*_*x,g*–1_ → *N*_*x,g*–1_ + 1): rate *b_i_* = *μ_g_N_x,g_*.

To simulate this time-evolution, we use the exact sampling of stochastic trajectories of the staircase model via a computationally efficient Next Reaction Method [56], which samples the next process that takes place and the time when it happens. Effectively, this method combines computationally efficient data structures and the Gillespie’s algorithm [57]:

1. sample the next process i, which happens with probability 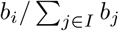,
2. sample the waiting time Δ*t*, which has distribution 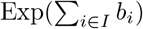
3. update the time *t* → *t* + *Δt* when the next process *i* happens and execute the process *i*.

Using this sampling technique, the evolutionary (resp. ecological) times *T_evo_* (resp. *T_eco_*) at a resistance state *R* can be sampled by measuring when mutants appear (resp. outcompete the wild-type) in the overlap region *x* = *R*+1. By averaging over many independent samples, the expected evolutionary and ecological times can be estimated and the adaptation rate 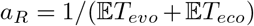 computed. As in [26], we have used 50 independent samplings. The same algorithm is used to simulate the extensions of the staircase model and the precise parameters used in simulations are detailed in the SI Text S8.

### B. Analytical and numerical techniques

In addition to computer simulations, we developed a combination of analytical and numerical techniques, which are used to compute the evolutionary time, ecological time and the adaptation rate. In contrast to simulations, these techniques are based on closed-form solutions of Master equations, numerical methods for solving algebraic equations (such as Newton-Raphson method) and solving ODEs (such as Runge-Kutta method). To illustrate these techniques, we show how to compute the expected waiting times and the adaptation rate *a_R_* in the original staircase model:

1. **Find the wild-type profile** *N_x_* **at a resistance state** *R*. Use Newton-Raphson method to solve the algebraic equations for the non-trivial fixed point of the mean-field theory

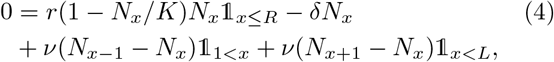

where 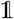 is the indicator function.
2. **Compute the evolutionary time** 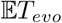. A closed-form solution exists and follows from a modification of the theory in [25, 26]. We consider three independent adaptation paths along which wild-type can produce a first mutant in the overlap region *x* = *R* + 1. On the position-genotype lattice (*x, g*), they are the division path D: (*R, R*) → (*R, R* +1) → (*R* + 1, *R* +1) (mutation at *x* = *R* followed by migration to the right); the overlap path O: (*R* + 1,*R*) → (*R* + 1, *R* + 1) (mutation at *x* = *R* + 1); and the no-division path N: (*R* + 2, *R*) → (*R* + 2, *R* + 1) → (*R* + 1, *R* + 1) (mutation at *x* = *R* + 2 followed by migration to the left). As the original theory [26] assumes low motility, it neglects the no-division path since there are no wild-type cells to mutate at position *x* = *R*+2 in this regime (SI Theorem 3). Moreover, we assume that the flux of mutants between neighbouring compartments in the division region *x* ≤ *R* and the no-divison region *x* ≥ *R* + 2 is equilibrated, so that we can ignore the net probability flux of mutants that enter or exit the compartments *x* = *R* – 1, *R, R* + 1 and also ignore longer paths such as (*R* – 1, *R*) → (*R* – 1,*R* +1) → (*R,R* +1) → (*R* +1,*R* +1) (mutation at *x* = *R* – 1 followed by double migration to the right). This is a reasonable assumption as the flux of mutant-producing wild-type is equilibrated far from the population front. The computation of 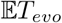 has the following structure.

a. The waiting times till a mutant is produced in the overlap region along path *i* ∈ {*D, O, N*} are denoted by *T_i_* (Fig. 2a).
b. The probability that the waiting will be longer than *t* is 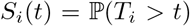 and the probability density function of *T_i_* is given by *F_i_*(*t*) = –d*S_i_*/d*t*.
c. Notice that *T_evo_* = min_*i*_ *T_i_* and that *T_i_* are independent. Thus, *S*(*t*) = *P*(*T_evo_* > *t*) = *S_D_* (*t*)*S_O_* (*t*)*S_N_* (*t*).
d. The probability density function of *T_evo_* is

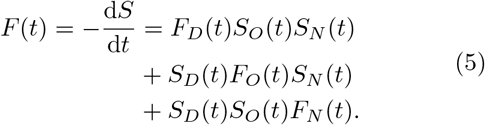
e. The expected time 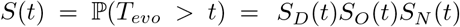 is found by numerical integration. Therefore, the problem is solved by finding the functions *S_i_*(*t*) and differentiating them to produce *F_i_*(*t*). According to [25, 26], these functions are:

a. **For path D:**

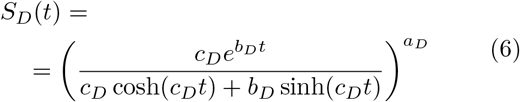

where

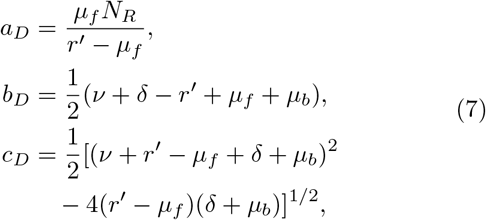

and

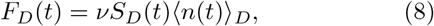

where 〈*n*(*t*)〉_*D*_ is the average mutant population in compartment *x* = *R* at time *t*

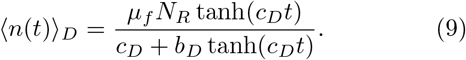
b. **For path O:**

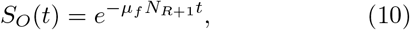

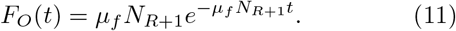
c. **For path N:**

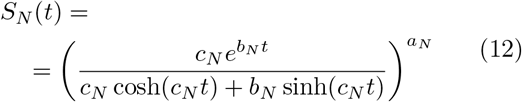

where

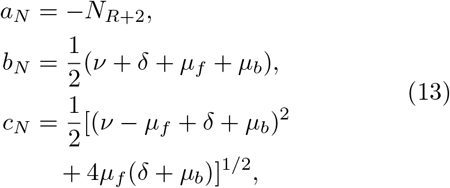

and

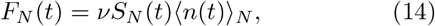

where 〈*n*(*t*)〉_*N*_ is the average mutant population in compartment *x* = *R* + 2 at time *t*

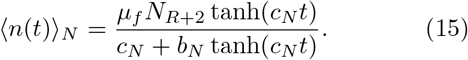
3. **Compute the ecological time** 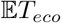. The competition between wild-type and mutants can be described well by the mean-field equations:

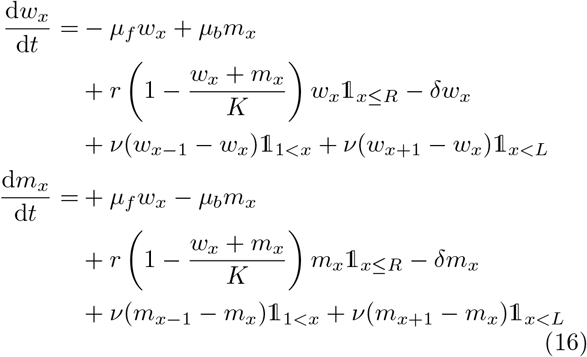

with initial conditions at time *t* = 0:

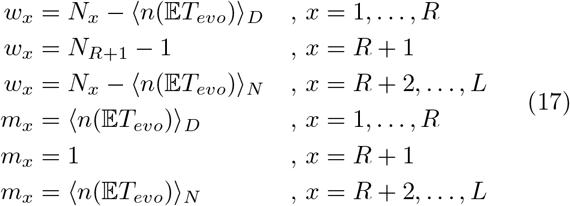 In these equations *w_x_* and *m_x_* stand for the number of wild-type and mutant cells in spatial compartment *x*, respectively. The system is simulated from the end state of the previous stochastic process when a first mutant appears in the overlap region. This system can be simulated efficiently by a Runge-Kutta (RK4) method. The time 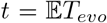 is reached when *w*_*R*+1_ < *m*_*R*+1_ for the first time, i.e., the mutants outcompete the wild-type in the overlap region.
4. **Find the adaptation rate.** The adaptation rate is found from

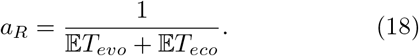

For convenience, we refer to this set of techniques as analytical techniques, even though they implement many numerical procedures. The benefit of the analytical techniques, as opposed to simulations, comes from their computational efficacy. In addition to these form of analytical techniques, we developed a purely analytical approach, which describes the formation of the wild-type profile and this approach is detailed in the Supplementary Text.

## IV. ACKNOWLEDGEMENTS

We thank Kacper Kornet and Deryck Thake from the IT team of the Department of Applied Mathematics and Theoretical Physics of the University of Cambridge for their help with the managing of the supercomputing cluster. We thank Julian Parkhill for discussions about the evolution of antibiotic resistance and for suggesting the study of HGT. We thank Kevin Foster, Oscar Despard, George Fortune, Pierre Haas, Julian Parkhill, Robert Austin, Mike Cates and Philip Maini for their comments on earlier versions of this manuscript. V.P. was supported by a CMS Summer Research Grant, and had additional support from Gonville & Caius College. N.M.O. was supported by a BBSRC Discovery Fellowship No. BB/T009098/1.

## SUPPLEMENTARY TEXT

The main purpose of the supplementary text (SI text) is to present the analytical study of the formation of wildtype profiles in our models. In the main text, we claim that the wild-type profile is key for bacterial adaptation in an antibiotic gradient as it shapes the fitness landscape of mutants (see Results). The quantitative analysis of the wild-type profile provides the first step to the computation of the evolutionary time, the ecological time and the adaptation rate (see Methods). Therefore, understanding the formation of the wild-type profile is key for both intuitive and quantitative analysis.

This SI text is organised as separate sections for convenience and clarity. We start by restating the analytical results for the wild-type profile of the simple source-sink model presented in the final subsection of Results. Starting from the end of the main text is deliberate because the source-sink model is a toy model, which provides an intuition for the formation of the wild-type profile in other more complex models considered in this work. However, the sections of the SI text are relatively self-contained and can be read in any order. We provide next a short guide to explain the content of each section and its links to other parts of our work:

- Section S1: summary of results about the wild-type profile in the source-sink model presented in Fig. 5 (fourth subsection of Results).
- Section S2: utilization of Section S1 to explain why the bifurcation argument for the emergence of critical motilities applies to a large variety of models (fourth subsection of Results).
- Section S3: wild-type profile in the staircase model presented in Fig. 1 and Fig. 2 (first and second subsections of Results).
- Section S4: modified staircase models and their wild-type profile, including the effect of resistance costs in Fig. S1, bactericidal antibiotics in Fig. S2, bacterial chemotaxis in Fig. S3 (second subsection of Results) and horizontal gene transfer in S6 (Discussion).
- Section S5: wild-type profile in the model of stochastic phenotypic switching presented in Fig. 3 and Fig. S4 (third subsection of Results).
- Section S6: wild-type profile in the model of density-dependent motility presented in Fig. 4 and Fig. S5 (third subsection of Results).
- Section S7: mathematical proofs of all theorems presented in all the SI sections mentioned above.
- Section S8: supplementary methods detailing the exact parameter choices for each figure of our work, including information about main and supplementary figures.

### S1. Source-sink model

In the main text, we studied the wild-type profile of the source-sink model (staircase with *L* = 2, *R* = 1), described by the dynamical system (1). The phase portraits of the system are shown in Fig. 5b and the results are summarised in Theorem 1.

#### Theorem 1.

*System* (1) *admits exactly two fixed points*

- *trivial fixed point: N*_2_ = *N*_1_ = 0,
- *non-trivial fixed point*:

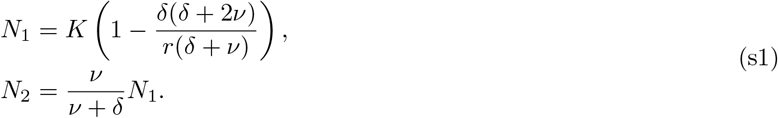

*Exactly one of these fixed points is stable and corresponds to a stable wild-type profile, with non-negative cell numbers N*_1,2_ ≥ 0. *Let* 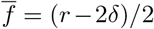 *be the average wild-type fitness. The stable wild-type profile corresponds to the non-trivial fixed point iff*

- *the environment is source-like* 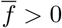 *and cell motility ν is arbitrary, or*
- *the environment is sink-like* 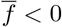 *and cell motility is constrained to v* ∈ [0, *ν_c_*], *where the critical motility is*

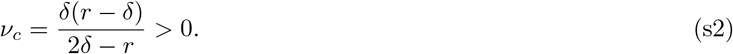

There are two implications of Theorem 1. First, this theorem provides an explanation for the existence of a critical motility as explained in the fourth subsection of Results and further discussed in the next SI section S2. Second, it explains the shape of the wild-type profile *N_x_* in Fig. 1d. The population in compartment *x* = 2 is significantly suppressed at low motility (*ν* ≪ *δ*) and the population is homogeneously distributed across space at high motility (*ν* ≫ *δ*). We elaborate on this phenomenon in the SI section S3.

### S2. Robust mechanism for the emergence of the critical motility

In the main text, we claimed that the critical motility can be understood as a bifurcation point of a dynamical system for the wild-type profile formation and that this mechanism is applicable to a large variety of models. Below we first provide a heuristic argument for why critical motilities shall arise for all models in our paper, and then we make a comparison with critical motilities in ecological models [S1-S3].

Theorems with analogous structure to Theorem 1 shall exist for more complicated models of our paper and shall predict the existence of a critical motility. Heuristically, we can argue as follows. Vary motility *ν* ≥ 0 as the bifurcation parameter. When there is no motility *ν* = 0, the dynamics between spatial compartments decouples and the nontrivial fixed point is stable while the trivial fixed point is not. As cell motility increases, the non-negative quadrant *N_x_* ≥ 0 becomes forward invariant and its boundary contains a single fixed point, the trivial fixed point. Assuming that the underlying Markov process is ergodic, a unique stable fixed point must exist within the non-negative quadrant *N_x_* ≥ 0. This implies that the non-trivial fixed point either stays in the positive quadrant and remains stable, or it leaves the positive quadrant through a transcritical bifurcation with the trivial fixed point, see Fig. 5b. Populations of high motility are generically governed by the average fitness [S4], explaining the stability of the trivial fixed point at high motility in sink-like environments. Therefore, there must exist a critical motility associated with the transcritical bifurcation if the environment is sink-like.

In the main text, we claimed that this bifurcation mechanism also applies to the critical motility in ecological models that study range expansion [S1-S3]. We elaborate on this claim with a migration-selection model from [S3] and relate this model to our source-sink model. Consider two alleles a and *A* and two habitats, such that allele *A* is selected for in habitat 1 with strength s and allele a is selected for in habitat 2 with strength *αs* (*α, s* > 0). Moreover, these alleles can move between habitats at rate *ν*. The relative frequencies of allele a in habitat *i* ∈ {1, 2} are denoted by *a_i_* ∈ [0,1], and their dynamics can be described by a replicator equation with constant selection:

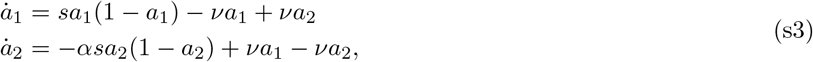

This is a dynamical system on a square (*a*_1_, *a*_2_) ∈ [0,1]^2^. When *ν* = 0, there is a globally attracting non-trivial fixed point (*a*_1_, *a*_2_) = (1,0). When *ν* > 0, the square [0,1]^2^ becomes forward invariant and the boundary supports two trivial fixed points: (*a*_1_, *a*_2_) = (0, 0) (allele A wins) and (*a*_1_, *a*_2_) = (1, 1) (allele a wins). Moreover, as *ν* increases from 0, the globally attracting fixed point is pushed from the boundary of the square into the interior. Similarly to the source-sink model, the non-trivial fixed point can leave the square only through a transcritical bifurcation with one of the trivial fixed points, giving rise to a critical motility *ν_c_*. A straightforward calculation for the non-trivial fixed point shows that it collides with the trivial fixed point (*a*_1_, *a*_2_) = (0,0) when *α* > 1 and with the trivial fixed point (*a*_1_, *a*_2_) = (1, 1) when 0 < *α* < 1 at

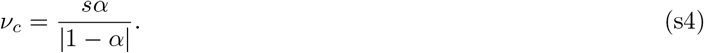

Therefore, in accordance with [S3], both alleles coexist if *ν* < *ν_c_* and the overall more fit allele takes over if *ν* > *ν_c_*.

The similarity between this model and the source-sink model is not a coincidence. The source-sink system (1) can be manipulated and rearranged into

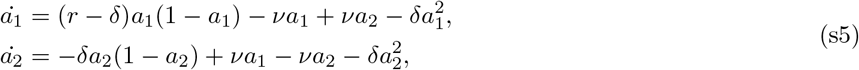

by relabelling *N_i_* ≡ *a_i_* ∈ [0, ∞). This is similar to (s3) with *s* = *r* – *δ*, *α* = *δ*/(*r* – *δ*) near the trivial fixed point (*a*_1_,*a*_2_) = (0,0). However, the point (*a*_1_,*a*_2_) = (1,1) is no longer a fixed point due to the corrections of 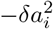. Therefore, when *α* > 1, the environment is sink-like and a critical motility exists, with (s4) matching (s2) upon substitution of *s* and *α*. When 0 < *α* < 1, the environment is source-like and no critical motility exists, since the corresponding trivial fixed point is removed from the dynamics by the correction terms 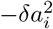.

### S3. Staircase model

In the main text, we studied the shape of the wild-type profile in the staircase model and its impact on mutant fitness (Fig. 1d). Hereby, we study this process analytically by using the same framework as for the source-sink model (SI section S1, fourth subsection of Results). The wild-type profile *N_x_* at state *R* of the adaptation process (Fig. 1b) is governed by a dynamical system (s6) on ℝ^*L*^, where *N_x_* is the total number of cells in spatial compartment *x*:

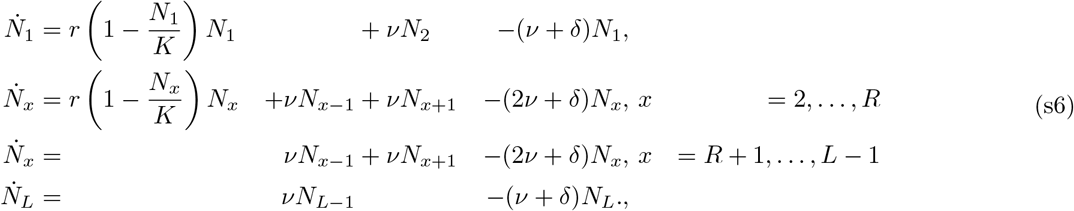

Similarly to the source-sink model, the non-negative quadrant *N_x_* ≥ 0 can only contain the trivial and the nontrivial fixed point^1^. Let us concentrate on the non-trivial fixed point *N_x_*. While a closed form expression for the fixed point does not exist for *R* > 1, important properties of the non-trivial fixed point *N_x_* can be derived analytically. To study the curvature of the wild-type profile, we define the convexity of the profile ^2^.

#### Definition 1.

*N_x_ is convex, resp. concave on* [*a, b*] *if for any x* ∈ [*a, b*] ⋂ {1, …, *L*}:

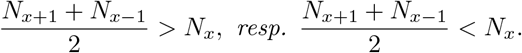

Using this definition, important qualitative properties of the wild-type profile can be derived.

#### Theorem 2.

*Let N_x_ be any fixed point of* (s6) *that lies in the strictly positive quadrant N_x_* > 0. *Note that N_x_ coincides with the non-trivial fixed point N_x_ if it is stable. This wild-type profile N_x_ of resistance R satisfies*:

1. *N_x_ is a decreasing function of x*.
2. *In any compartment above the staircase, the effective wild-type birth rate r*(1 – *N_x_*/*K*) *is bigger than the death rate δ*.
3. *N_x_ is concave above the staircase, i.e., on* [1,*R* + 1].
4. *N_x_ is convex below the staircase, i.e., on* [*R, L*].
5. *The curvature of the wild-type population profile at position x is proportional to wild-type fitness* 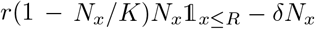 *at this position and inversely proportional to motility.*
6. *The mutant fitness* 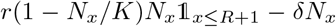 *is a positive, convex and increasing function on* [1,*R* + 1], *and a constant* –*δ function on* [*R* +2, *L*]. *In particular, mutant fitness is maximised in the overlap region and matches the wild-type fitness characterised above in compartments x ≤ R*.
7. *In any compartment N_x_* < *K*(1 – *δ/r*).
8. *r > δ is a necessary condition for the existence of this non-trivial fixed point in the strictly positive quadrant.*

Theorem 2 explains the generic shape of a wild-type profile for any motility *ν* (Fig. 1d) and the shape of the mutant fitness landscape. However, it does not explain the difference between the low and high motility regimes discussed in the main text. In order to understand the behaviour of the wild-type profile at low and high motility, asymptotic expansion for the non-trivial fixed point are needed.

#### Theorem 3.

*In the low motility regime v* ≪ *δ, the non-trivial fixed point is given by*:

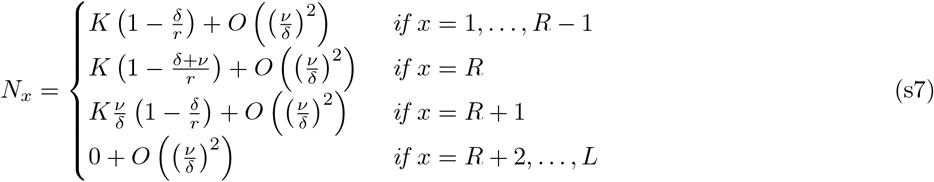

*In the high motility regime v* ≫ *δ, the non-trivial fixed point is given by:*

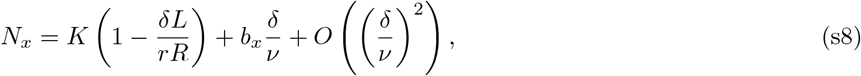

*where the coefficients b_x_ satisfy*

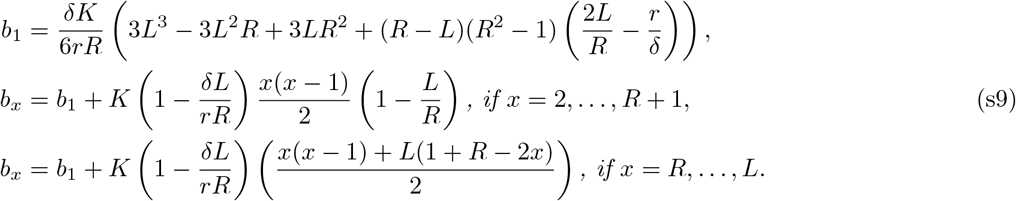

Using Theorem 3, we can see that the shape of the population front is always the same in the low motility regime and overlap region *x* = *R* +1 provides a refuge for mutants (Fig. 1d). The high motility regime leads to a homogeneous wild-type population *N_x_*, which increases with resistance state *R* and competes with mutants for space (Fig. 1d). Moreover, this fixed point can be in the negative quadrant to first order if the environment is sink-like. In this case, the non-trivial fixed point leaves the positive quadrant *N_x_* > 0 at a critical motility *ν_c_* and exchanges its stability with the trivial fixed point via a transcritical bifurcation.

#### Corollary 1.

*Let* 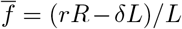 *be the average wild-type fitness. Then, the stable wild-type profile is non-vanishing iff*

- *the environment is source-like 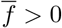 and motility ν is arbitrary, or*
- *the environment is sink-like 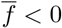 and motility is constrained to ν ∈* [0, *ν_c_*], *where to first order*

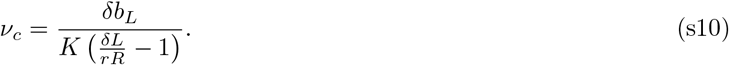

In the main text, we find a deadly motility regime characterised by extinction and no adaptation when *ν > v_c_*. We can prove that the adaptation rate is vanishing by deriving its upper bound.

#### Theorem 4.

*Fix a resistance state R and define* 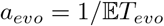. *Then*,

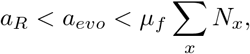

*In particular, when critical motility v_c_ exists, the adaptation rate α_R_ decreases to 0 faster than μ_f_*Σ_*x*_ *N_x_ as* 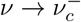. *Moreover, no adaptation can occur if the wild-type population is extinct*.

### S4. Modified staircase model

In the main text, we noted that the staircase model can be modified to consider the effect of fitness costs of resistance (Fig. S1), bactericidal antibiotics (Fig. S2), bacterial chemotaxis (Fig. S3) and horizontal gene transfer (Fig. S6). In this section, we explain each of these modifications of the original staircase model in turn and present the results for the modified models, with the key conclusion that motility governs the bacterial adaptation by different mechanisms in the low motility, high motility and deadly motility regimes as identified in the original staircase model (Fig. 2a). We finish this section with a mathematical analysis of the wild-type profile.

#### a. Resistance costs

The original staircase model assumes that there are no fitness costs associated to resistance. Yet, adaptations that confer antibiotic resistance often have a cost that is expressed in the absence of antibiotic pressure, and in these cases resistant genotypes often grow slower than susceptible ones [S5, S6]. We model such fitness costs of resistance by decreasing the division rate of cells above the staircase *g* ≥ *x* by a factor of (1 – *c*)^*g−x*^, where *c* ∈ [0,1) is the resistance cost [S7]. We find that sufficiently high resistance costs can decrease the adaptation rate *a_R_* when cell motility is sufficiently low (Fig. S1). This agrees with [S8] that shows that the resistance cost *c* affects the adaptation rate precisely when cell motility is very low (*ν* < *c*^2^*δ*). The rationale for this result is simple. When cell motility is low, resistant genotypes that harbour a resistance cost are less able to avoid competition with the wild-type and cannot easily move to where they have higher growth advantage. Importantly, we find that resistance costs do not affect our key conclusion: low motility accelerates adaptation, high motility decelerates adaptation and very high motility leads to extinction if the average wild-type fitness 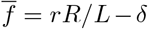 is negative. As a result, the deadly motility regime is not influenced by resistance costs (Fig. S1).

#### b. Bactericidal antibiotics

The original staircase model studies the effect of antibiotics that inhibit bacterial growth (i.e., we considered bacteriostatic antibiotics). However, many natural and clinical antibiotics kill bacteria (bactericidal antibiotics). We model bactericidal antibiotics by increasing the death rate under the staircase *σ*-times as in [S7]. Importantly, our results show that bacterial adaptation to bactericidal antibiotics again has the same regimes dictated by cell motility (Fig. S2). Interestingly, while bacterial adaptation is identical to the one found for bacteriostatic antibiotics in the low motility regime, a deadly motility regime can be induced at high motility if the bactericidal effect is sufficiently strong *σ* ≫ 1. In this case, bacteria experience deadly antibiotic concentrations more often and the average wild-type fitness 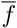 is decreased by *σ*, as derived later.

#### c. Bacterial chemotaxis

The original staircase model assumes that bacteria move randomly. Yet, it is well-known that bacteria can bias their motion in chemical gradients (chemotaxis). In particular, it has been shown that antibiotics can trigger both positive [S9] and negative [S10] chemotaxis. We model the effect of biased motility by changing the probability *p* of bacterial movement up or down the antibiotic gradient. Notably, the usual adaptation regimes can be identified in this model (Fig. S3). Our results show that chemotaxis has little impact on the adaptation rate when cell motility is low (Fig. S3), mimicking what we found for bactericidal antibiotics described above. However, when motility is high, positive chemotaxis *p* > 0.5 increases antibiotic exposure and promotes the deadly motility regime (SI Movie S3), while negative chemotaxis *p* < 0.5 decreases antibiotic exposure and prevents the deadly motility regime (SI Movie S4). Put differently, chemotaxis changes the average wild-type fitness 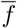, decreasing it for positive chemotaxis and increasing it for negative chemotaxis, as derived later. For negative chemotaxis, the wild-type has higher average fitness but resides far from higher antibiotic concentrations. Therefore, mutants are unlikely to reach the overlap region and the adaptation rate is decreased (Fig. S3).

#### d. Horizontal gene transfer

Bacteria can exchange resistance genes through horizontal gene transfer (HGT), which is known to accelerate bacterial adaptation [S11]. In particular, it has been shown that the immigration of susceptible cells from a detached habitat can promote adaptation in a focal habitat via HGT [S12]. However, for closed systems without immigration of susceptible genotypes, it has been argued that HGT does not affect bacterial adaptation substantially [S13–S17]. In an attempt to recapitulate these results, we model HGT in our system by allowing less resistant genotypes g to acquire resistance from *N_x,g’_* more resistant cells of genotype *g*’ > *g* located at the same position *x* at a rate *h* × *N_x,g_*/. For realistic values of HGT rate *h* (< 1 transfer per generation [S12]), we find that the adaptation rate is slightly increased, but our key conclusion still holds: low motility accelerates bacterial adaptation, while high motility decelerates it. Moreover, we find that bacteria with very high motility experience deadly motility regimes if they have negative average fitness 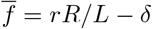, irrespective of HGT rate (Fig. S6).

#### e. Wild-type profile

So far, we considered various modifications of the staircase model. We now aim to understand how the conditions for the deadly motility regime change in the modified models. We see that there is no change to the deadly motility regime in the case of resistance costs and HGT (Fig. S1, S6). Therefore, we focus on the bactericidal antibiotics and bacterial chemotaxis that impact the deadly motility regime by modulating the average wild-type fitness 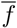 (Fig. S1, S6). We find the formulae for the average wild-type fitness 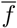 in these two modified models by studying the wild-type profiles in general modified staircase models with spatially variable division rate *r_x_*, death rate *δ_x_* and motility rate *ν_x_*. In this setting, chemotaxis corresponds to the choice *ν_R,x_* = 2*pv* and *ν_L,x_* = 2(1 – *p*)*ν*, where *p* is the probability to move up the gradient; the effect of bactericidal antibiotics corresponds to stress-induced death rate *δ_x_* = *σδ* below the staircase (*x* > *R*) and the usual death rat*e δ_x_* = *δ* above the staircase (*x* ≤ *R*). The wild-type profile formation at state *R* of the adaptation process (Fig. 1b) is governed by the mean-field equations:

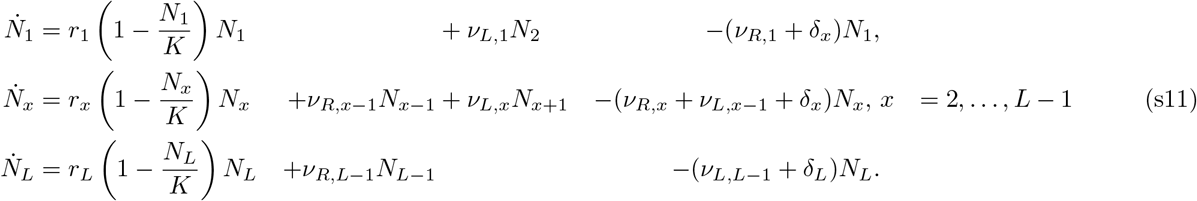

This dynamical system has two fixed points of interest, the trivial fixed point *N_x_* = 0 and the non-trivial fixed point, which correspond to possible wild-type populations. Next, we compute the non-trivial fixed point at low and high motility and derive results for the existence of a critical motility. To do this, we assume that *ν_L,x_,v_R,x_* ~ *O*(*ν*), *δ_x_* ~ *O*(*δ*), which is true for the modifications mentioned above.

##### Theorem 5.

*In the low motility regime ν ≪ δ, the wild-type profile is given by:*

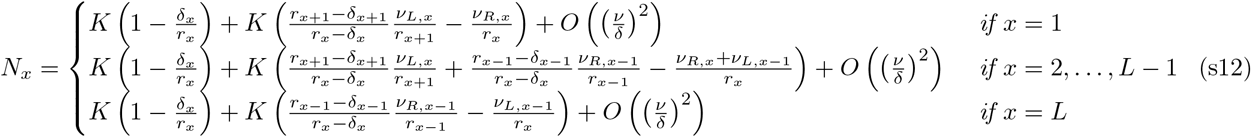

*In the high motility regime v* ≫ *δ, the wild-type profile is given by*:

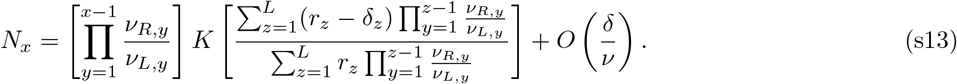

*The average wild-type fitness therefore is* 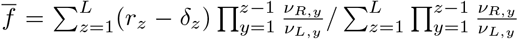. *The stable non-vanishing wild-type profile exists iff*

1. *the environment is source-like* 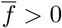 *and motility ν is arbitrary, or*
2. *the environment is sink-like* 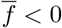 *and motility is constrained to v* ∈ [0, *ν_c_*] *for a critical motility v_c_*.

We note that this result coincides with the one from the basic staircase model (SI Theorem 3, SI Corollary 1) if *or_x_* = *r* above the staircase and 0 otherwise, *δ_x_* = *δ* and *ν_R,x_* = *ν_L,x_* = *ν*. Furthermore, this result explains the forms of the average wild-type fitness 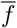 that were presented above for bactericidal antibiotics and chemotaxis.

For bactericidal antibiotics, the antibiotics-induced death increases *σ*-times, which decreases the average wild-type fitness 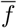 below zero if *σ* is sufficiently high, as

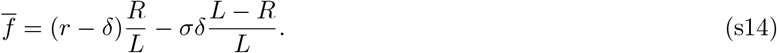

Therefore, Theorem 5 explains why the bactericidal effect σ converts the high motility regime into the deadly motility regime, while leaving adaptation in the low motility regime almost unchanged (Fig. S2). Intuitively, at low motility regime the population is not present in stressful regions below the staircase, making the adaptation invariant of antibiotics-induced death. In the high motility regime, the population experiences the average environment with increased overall death rate, which decreases the average wild-type fitness.

For chemotaxis, probability p that a cell moves up the antibiotic gradient decreases (resp. increases) the average wild-type fitness 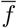 below (resp. above) zero when *p* is sufficiently high (resp. low), as

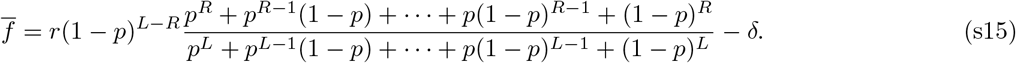

This result means that positive chemotaxis can always create a deadly motility regime (irrespective of *R*), and negative chemotaxis can always prevent a deadly motility regime (irrespective of *R*), as we confirm with simulations (Fig. S3). Theorem 5 also shows that, at high motility, the wild-type profile of a population with negative chemotaxis becomes concentrated away from the overlap region. This result explains the reduction in the adaptation rate in Fig. S3 as a consequence of reduced antibiotic exposure. Moreover, Theorem 5 also predicts that chemotaxis does not affect the wild-type at low motility, explaining why the adaptation rate is not affected at low motility in Fig. S3. We note that it is easier to introduce the deadly motility regime by controlling chemotaxis (power law dependence on *p*) rather than controlling bactericidal effects of antibiotics (linear dependence on *σ*).

### S5. Stochastic switching of motility phenotypes

In the main text, we studied the wild-type profiles of phenotypically heterogeneous populations with bacteria switching between different motility rates stochastically. Hereby, we study this process analytically. For convenience, we modify the notation slightly. The phenotypes have cell numbers *N_x_* and *M_x_*; motilities *ν_N_* and *ν_M_*; and phenotypes switch stochastically at rate *s*. Without loss of generality, we assume that *ν_M_* ≥ *ν_N_*. At ecological timescales, the wild-type profile is governed by a dynamical system (s16):

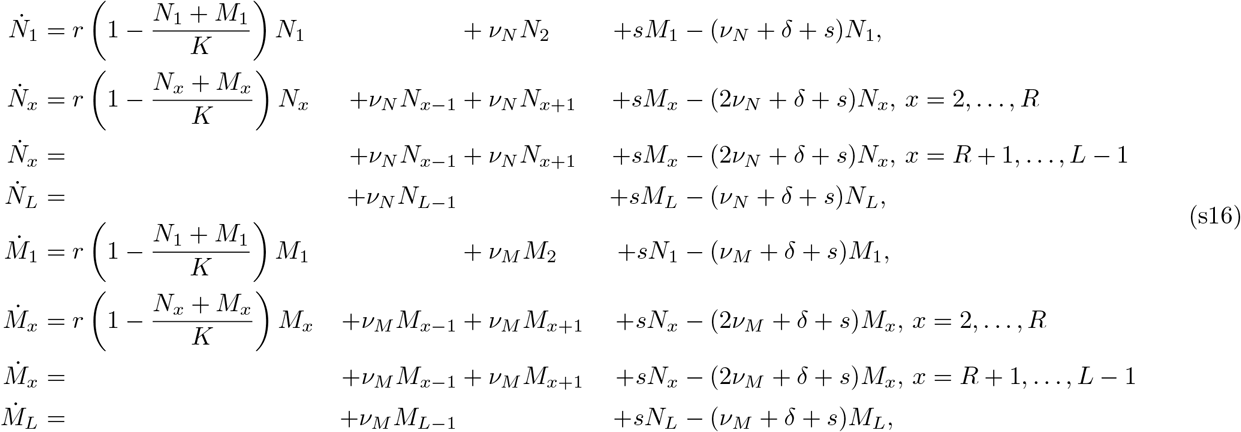

As before, the wild-type profile approaches the unique stable fixed point in the non-negative quadrant, which is either the trivial or the non-trivial fixed point. We next discuss the dynamics in the source-sink model (*L* = 2, *R* =1) and state the results for the staircase model (any *L, R*).

In the source-sink model, there is a closed form expression for the wild-type profile.

#### Theorem 6.

*The dynamical system* (s16) *for the source-sink model* (*L* = 2, *R* = 1) *has two fixed points*:

- *the trivial fixed point: N*_1,2_ = *M*_1,2_ = 0,
- *the non-trivial fixed point*:

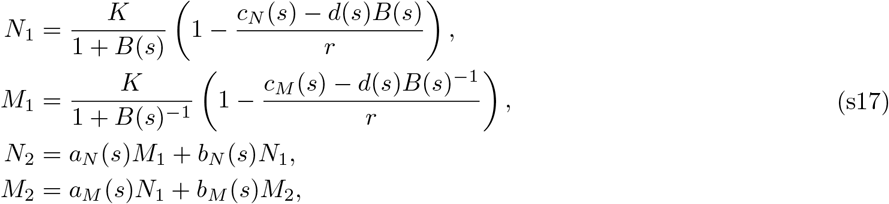

*with coefficients*

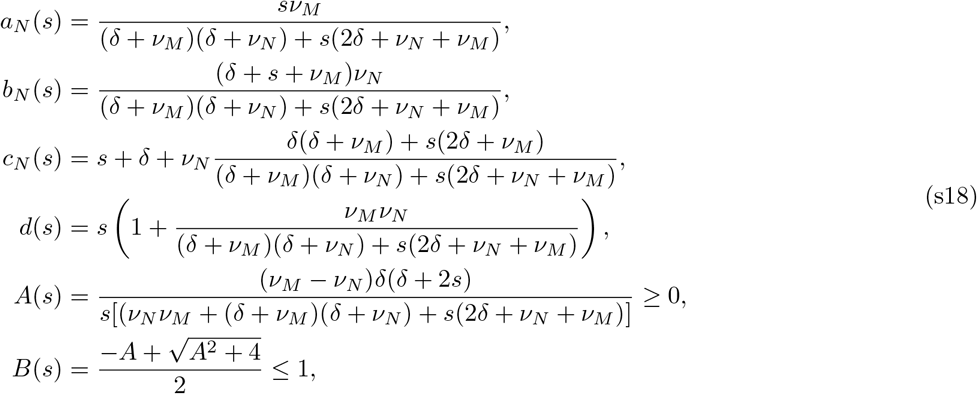

*and other follow from N* ↔ *M symmetry. Importantly, M*_1_ = *B*(*s*)*N*_1_.

In the main text, we claimed that the diagonal of Fig. 3b corresponds to the population of uniform motility *ν_N_* = *ν_M_* ≡ *ν*. A dynamical system for (*N* + *M*)_1,2_ behaves as the basic source-sink model of Theorem 1, leading to a non-trivial fixed point

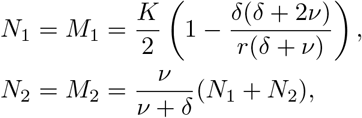

which is unaffected by switching rate *s*. In the main text, we also showed that phenotypic switching can modulate the adaptation regime. This effect is controlled by the dimensionless number *A*(*s*). As *A*(*s*) contains only the *δ, s* rates (besides motilities), the low switching (*A* ≫ 1, *s* ≪ *δ*) and high switching (*A* ≪ 1, *s* ≫ *δ*) regimes can be distinguished and described.

#### Corollary 2.

*At low switching* (*A* ≫ 1, *s* ≪ *δ*), *the non-trivial fixed point of the source-sink model* (*L* = 2, *R* = 1) *is*:

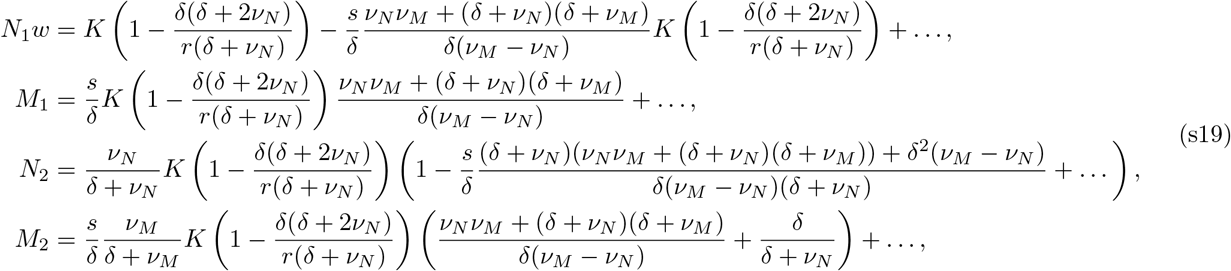

*with the dimensionless coefficients*

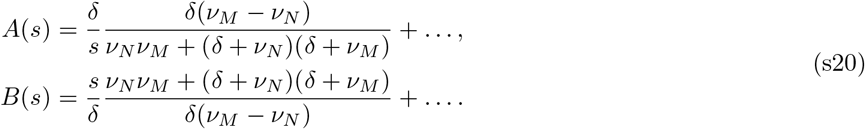

*In this case, the slower phenotype dominates the faster phenotype* (*N*, 2 ≫ *M*, 2) *and the system behaves as if only the slower phenotype was present* ((*N* + *M*)_1,2_ *matches Theorem 1 with v* = *ν_N_*).

*At high switching* (*A* ≪ 1, *s* ≫ *δ*), *the non-trivial fixed point of the source-sink model* (*L* = 2, *R* = 1) *is*:

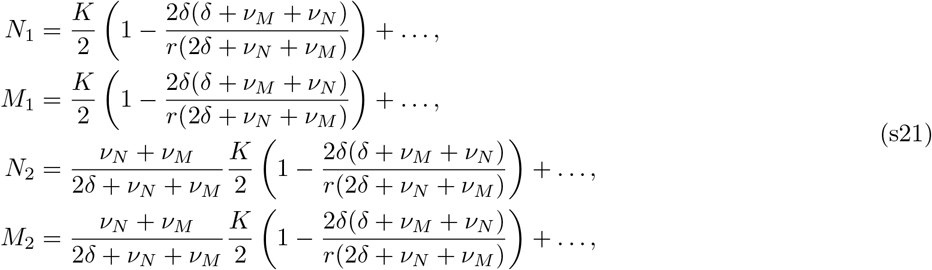

*with dimensionless coefficients*

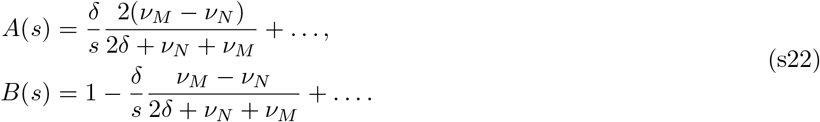

*In this case, both phenotypes are similarly important* (*N*_1,2_ ≈ *M*_1,2_) *and the system behaves as if only an average phenotype was present* ((*N* + *M*)_1,2_ *matches Theorem 1 with v* = (*ν_M_* + *ν_N_*)/2).

This result proves the following claim we made in the main text that at low switching (*A* ≪ 1, *s* ≫ *δ*), the population adapts as if only the slower phenotype was present, and that at high switching (*A* ≪ 1, *s* ≫ *δ*), the population adapts as if it had a uniform motility that matches the average motility.

In the main text, we also discussed the deadly motility regime. In contrast to systems with uniform motility, the deadly motility regime now occupies the (*ν_N_*, *ν_M_, s*) space; and instead of critical motility, there is a critical surface in the (*ν_N_, v_M_, s*) space.

#### Corollary 3.

*The deadly motility regime for the source-sink model* (*L* = 2, *R* = 1) is *described as a subspace of* (*ν_N_*, *ν_M_, s*) *space such that*:

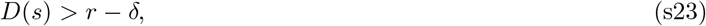

*with*

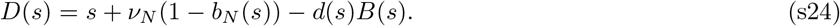

*D*(*s*) *has the following properties*:

- *D*(*s*) *is a strictly increasing smooth function of s* ∈ ℝ^+^
- 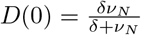
- 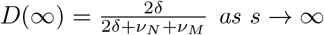 *as s* → ∞

*The deadly motility regime exist whenever the environment is sink-like* 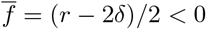, *and*

- *the slower phenotype is in the deadly motility regime* min(*ν_N_, v_M_*) > *ν_c_, and switching s is arbitrary, or*
- *the average phenotype is in the deadly motility regime* (*ν_N_* + *ν_M_*)/2 > *ν_c_, and switching s is above some critical value s* > *s_c_*(*ν_N_, v_M_*).

*In particular, the critical surface asymptotes with* min(*ν_N_, v_M_*) = *ν_c_ at low switching* (*s* ≪ *δ*) *and with* (*ν_N_* + *ν_M_*)/2 = *ν_c_ at high switching* (*s* ≫ *δ*).

For completeness, we state the results for the staircase model, which can be used to support the same claims as those made above for the simpler source-sink model. The asymptotic form of the wild-type profile can be derived.

#### Theorem 7.

*In the low motility combination v_N_, v_M_ ≪ δ, the non-trivial fixed point of the staircase model is given by*:

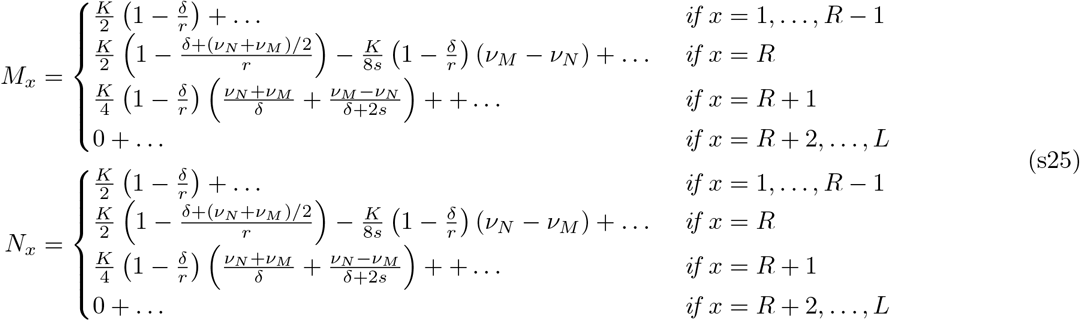

*In the high motility combination v_N_, v_M_ ≫ δ, the non-trivial fixed point of the staircase model is given by*:

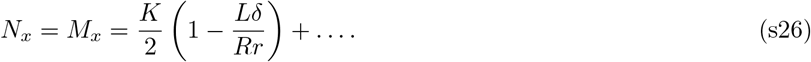

*In the mixed motility combination v_N_ ≪ δ ≪ v_M_, the non-trivial fixed point of the staircase model is given by*

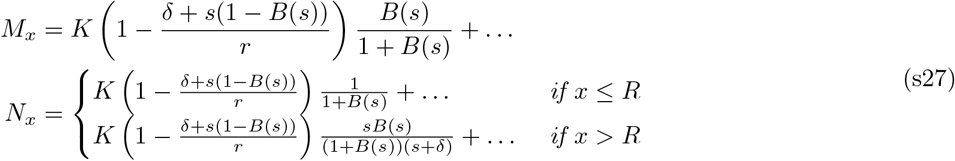

*with*

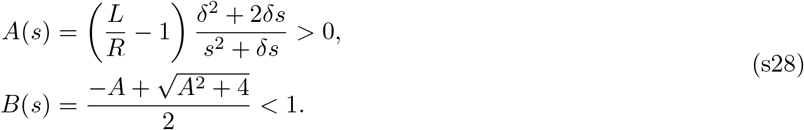

*Importantly, to lowest order N_x_/M_x_* = *B*(*s*) *in the region above the staircase x* ≥ *R*.

A non-perturbative result characterising the behaviour of the staircase model at low and high switching can also be derived analytically.

#### Theorem 8.

*At low switching s* ≪ *δ, the overall wild-type profile behaves as if it has a single motility of v* = min(*ν_N_, v_M_*). *At high switching s* ≫ *δ, the overall wild-type profile behaves as if it has a single motility of ν* = (*ν_N_ + v_M_*)/2.

*In particular, this result affects the deadly motility regime in the* (*ν_N_, v_M_, s*) *space. Let* 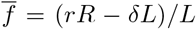 *be the average wild-type fitness. The wild-type profile is non-trivial iff*:

- *the environment is source-like* 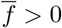, *motility rates v_N_, v_M_ are arbitrary, and switching rate s is arbitrary,*
- *the environment is sink-like* 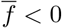 *and*

– *switching is low s* ≪ *δ and the slower phenotype is in the deadly motility regime* min(*ν_N_, v_M_*) > *ν_c_*,
– *switching is high s* ≫ *δ and the average-motility phenotype is in the deadly motility regime* (*ν_N_ v_N_*)/2 > *ν_c_*.

### S6. Density-dependent motility

In the main text, we studied the wild-type profiles of populations with density-dependent motility. We briefly outline important features of these wild-type profiles. The wild-type profile can be described with the mean-field equations:

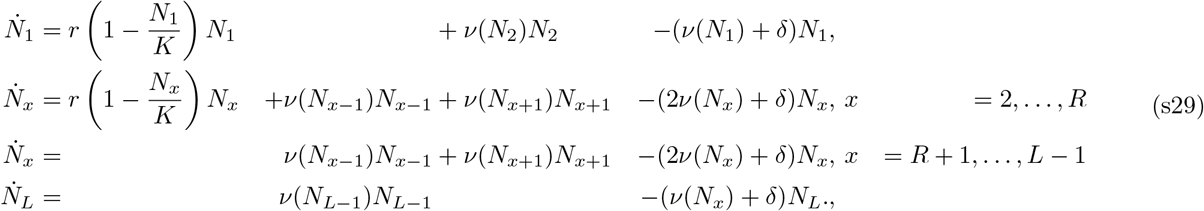

where

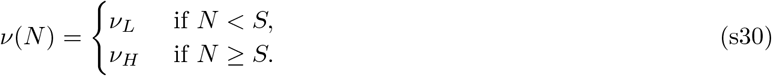

Since the forcing of this dynamical system is discontinuous whenever *N_x_* = *S*, the fixed-point analysis is more complicated and we only mention two important features of this system.

First, multiple stable fixed points can exist. For example, consider a situation when a low-density motility phenotype has motility above the critical motility *ν_L_* > *ν_c_* and the high-density motility phenotype is slower *ν_H_* < *ν_L_*. If the system is initially started from a low density, the low-density motility phenotype cannot survive and a deadly motility regime occurs as in Fig. 4c (attraction to the trivial fixed-point). However, if the system is initially started from a high density, the high-density motility phenotype can survive and no deadly motility regime would have occured in Fig. 4c (attraction to a non-trivial fixed-point).

Second, the surfaces of discontinuous forcing (*N_x_* = *S* for some *x*) might attract the trajectories. This result often happens in the slow-to-fast switching case, see Fig. 4b at low threshold *S*. In this case, the bulk of the high-density population is driven very close to density *N_x_* = *S*.

### S7. Proofs

#### Theorem 1.

*System* (1) *admits exactly two fixed points*

- *trivial fixed point: N*_2_ = *N*_1_ = 0,
- *non-trivial fixed point:*

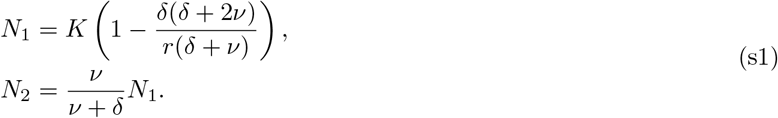

*Exactly one of these fixed points is stable and corresponds to a stable wild-type profile, with non-negative cell numbers N*_1,2_ ≥ 0. *Let* 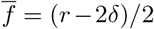 *be the average wild-type fitness. The stable wild-type profile corresponds to the non-trivial fixed point iff*

- *the environment is source-like* 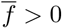 *and cell motility ν is arbitrary, or*
- *the environment is sink-like* 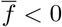 *and cell motility is constrained to v* ∈ [0, *ν_c_*], *where the critical motility is*

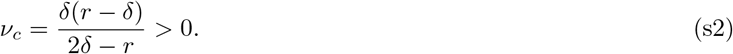

*Proof.* The two fixed points can be found by setting 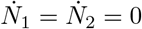. The stability of these fixed points can be checked by considering the signs of trace and determinant of the Jacobian matrix:

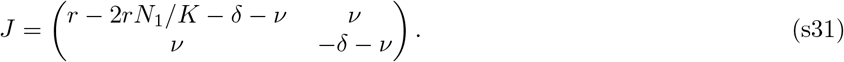

For example, the trivial fixed point has tr *J* = *r* – 2*δ* – 2*ν* and det *J* = –*δ*(*r* – *δ*) + *ν*(2*δ* – *r*). If 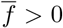, or if 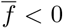 and *ν* < *ν_c_*, then det *J* < 0 and the trivial fixed point is a saddle node. Otherwise, det *J* > 0, tr *J* < 0 and the trivial fixed point is stable. Similar analysis applies to the non-trivial fixed point.

#### Theorem 2.

*Let N_x_ be any fixed point of* (s6) *that lies in the strictly positive quadrant N_x_* > 0. *Note that N_x_ coincides with the non-trivial fixed point N_x_ if it is stable. This wild-type profile N_x_ of resistance R satisfies*:

1. *N_x_ is a decreasing function of x*.
2. *In any compartment above the staircase, the effective wild-type birth rate r*(1 – *N_x_*/*K*) *is bigger than the death rate δ*.
3. *N_x_ is concave above the staircase, i.e., on* [1,*R* + 1].
4. *N_x_ is convex below the staircase, i.e., on* [*R, L*].
5. *The curvature of the wild-type population profile at position x is proportional to wild-type fitness* 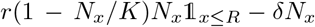 *at this position and inversely proportional to motility*.
6. *The mutant fitness* 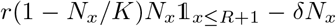 *is a positive, convex and increasing function on* [1,*R* + 1], *and a constant* –*δ function on* [*R* + 2, *L*]. *In particular, mutant fitness is maximised in the overlap region and matches the wild-type fitness characterised above in compartments x* ≤ *R*.
7. *In any compartment N_x_* < *K*(1 – *δ/r*).
8. *r* > *δ is a necessary condition for the existence of this non-trivial fixed point in the strictly positive quadrant.*

*Proof. Part 1 and 2:* Rewrite the fixed point equation for (s6) as

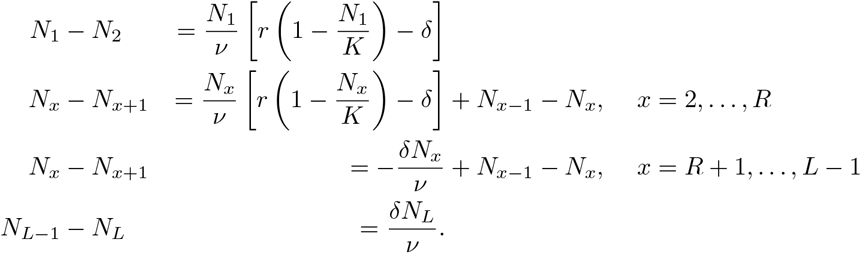

Firstly, assume that *N*_1_ ≤ *N*_2_. The first equation implies that *r*(1 –*N*_1_/*K*)–*δ* ≤ 0. As *N*_1_ ≤ *N*_2_, also *r*(1–*N*_2_/*K*)–*δ* ≤ *r*(1 – *N*_1_/*K*) – *δ* ≤ 0. Then, the second equation implies that *N*_2_ ≤ *N*_3_. Inductively, the first *L* – 1 equations imply that

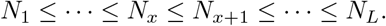

However, the last equations then implies that *N_L_* ≤ 0, contradicting our restriction to *N_x_* > 0. Therefore, *N*_1_ > *N*_2_. The first equation implies that *r*(1 – *N*_1_/*K*) – *δ* > 0. As *N*_1_ > *N*_2_, *r*(1 – *N*_2_/*K*) – *δ* > *r*(1 – *N*_1_/*K*) – *δ* > 0. The second equation implies that *N*_2_ – *N*_3_ > 0. Inductively, the first *L* – 1 equations imply the conclusion of part 1 that

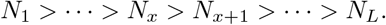

Notice that along the way, part 2 has also been proven.

*Part 3:* Notice that above the staircase,

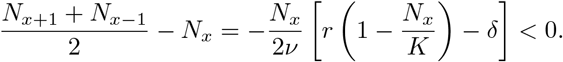

*Part 4:* Notice that below the staircase,

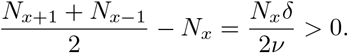

*Part 5:* Write the fixed point equation in the form:

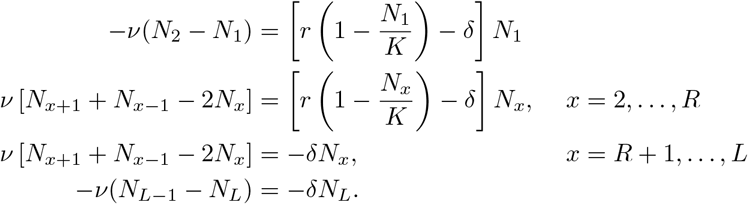

*Part 6:* The mutant fitness is given by the difference between the effective birth-rate (resulting from competition with wild-type) and death rate. Therefore, on [1, *R* + 1], this mutant fitness is

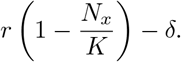

Therefore, it is increasing (by part 1), positive (by part 2) and convex (by part 3). On [*R* + 2, *L*], mutants cannot divide and their fitness is –*δ*.

*Part 7:* Follows directly from part 2.

*Part 8:* If *r* ≤ *δ*, part 6 implies that *N_x_* ≤ 0.

#### Theorem 3.

*In the low motility regime v* ≪ *δ, the non-trivial fixed point is given by*:

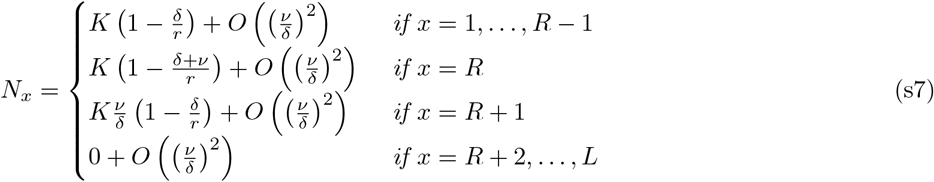

*In the high motility regime ν ≫ δ, the non-trivial fixed point is given by*:

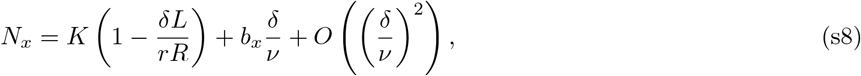

*where the coefficients b_x_ satisfy*

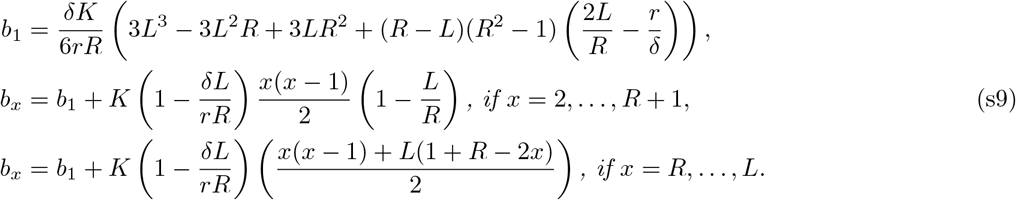

*Proof.* For the low motility regime, assume that 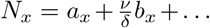 Then, at *O*(1) of (s6), we recover the equilibrium of local birth/death dynamics:

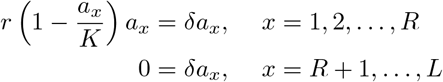

implying that *a_x_* = *K*(1 – *δ/r*) when *x* ≤ *R*, and *a_x_* = 0 otherwise. At *O*(*ν/δ*), we recover the first order corrections to balance this local dynamics with curvature of the population curve:

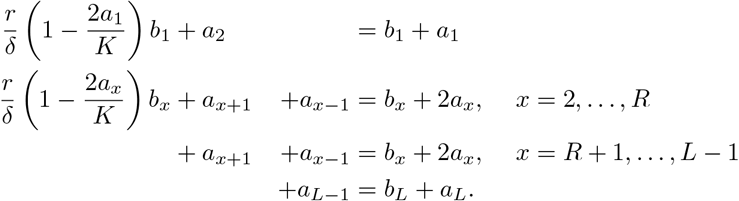

Using the previous formulae for *a_x_* this can be solved for *b_x_*. The result follows.

For the high motility regime, assume that 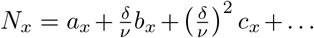 Then, at 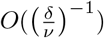, we only get the spatial diffusion with neglected local birth-death processes:

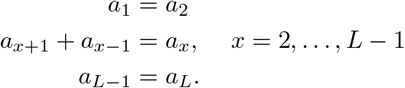

This result implies that *a_x_* = *α* for some constant *α*. Also notice that this system is under-determined and constant *α* must be fixed from the equations at the next order. At *O*(1), the first order correction to curvature due to local birth/death processes is:

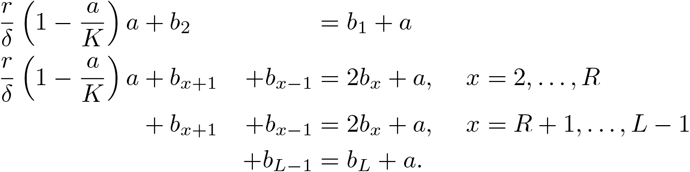

Summing the equations yields

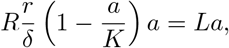

and thus

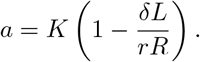

With this choice of *a*, the system of equations for *b_x_* takes a form of under-determined system of difference equations. By solving this system, *b_x_*’s can can be expressed in terms of *b*_1_ as in (s9). In order to fix *b*_1_, the equations for *c_x_* at order 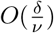 need to be found:

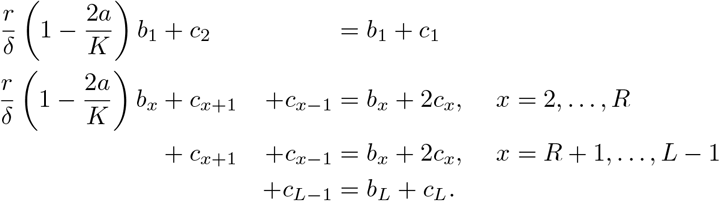

Summation cancels the cx coefficients and gives a single equation for *b_x_*:

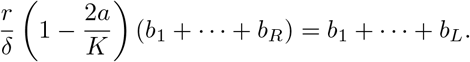

Using the relationships for *b_x_*:

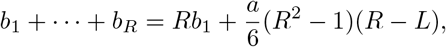

and

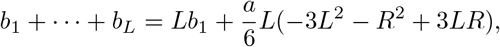

the coefficient *b*_1_ is obtained as in (s9).

#### Corollary 1.

*Let* 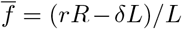 *be the average wild-type fitness. Then, the stable wild-type profile is non-vanishing iff*

- *the environment is source-like* 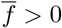 *and motility ν is arbitrary, or*
- *the environment is sink-like* 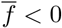 *and motility is constrained to v* ∈ [0, *ν_c_*], *where to first order*

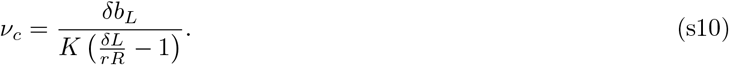

*Proof.* Notice that a stable wild-type profile *N_x_* is expected to be decreasing in *x* by Theorem 2. Hence, a non-trivial fixed point stays within the positive quadrant and remains stable iff *N_L_* > 0. To the lowest order, this condition is equivalent to 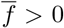. If 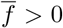, the wild-type profile exists for any *ν*. If 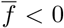, the first order approximation dictates that *N_L_* > 0 iff *ν* < *ν_c_* with *ν_c_* given by (s10). At *ν* = *ν_c_*, the non-trivial wild-type fixed point collides with the trivial wild-type fixed point and exchanges stability through a transcritical bifurcation.

#### Theorem 4.

*Fix a resistance state R and define* 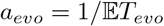. *Then*,

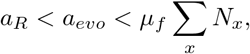

*In particular, when critical motility v_c_ exists, the adaptation rate a_R_ decreases to* 0 *faster than μ_f_* ∑_*x*_ *N_x_ as* 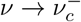. *Moreover, no adaptation can occur if the wild-type population is extinct.*

*Proof.* To find the upper bound on *a_evo_*, we need to find the lower bound on the waiting time *T_evo_* till the stable wild-type population produces a mutant in the overlap region (mutants migrating from other compartments count as well). This waiting time *T_evo_* is bounded below by the waiting time till a mutant is produced anywhere *T_a_*, i.e., *T_evo_* > *T_a_* In each spatial compartment *x*, only the most resistant wild-type cells can mutate. Call their number 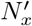. Then, the waiting time *T_x_* till the compartment *x* produces a mutant is 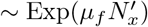. The waiting times *T_x_* are independent and *T_a_* = min_*x*_ *T_x_*. Therefore, 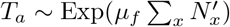. Finally,

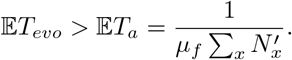

Thus,

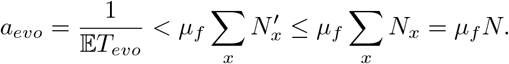

Result follows.

#### Theorem 6.

*The dynamical system* (s16) *for the source-sink model* (*L* = 2, *R* = 1) *has two fixed points*:

- *the trivial fixed point: N*_1,2_ = *M*_1,2_ = 0,
- *the non-trivial fixed point:*

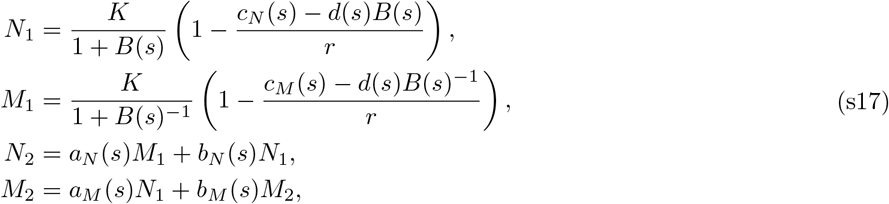

*with coefficients*

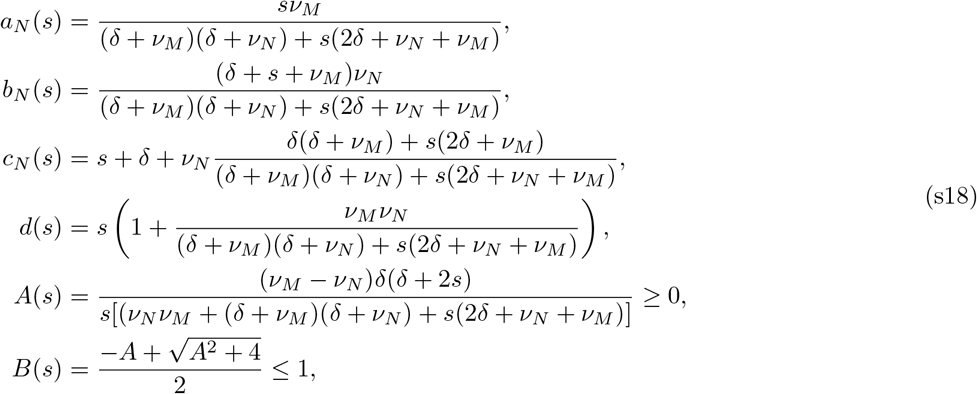

*and other follow from N* ↔ *M symmetry. Importantly, M*_1_ = *B*(*s*)*N*_1_.

*Proof.* Notice that the system (s16) is symmetric under *M* ↔ *N*. Now, express *M*_2_, *N*_2_ in terms of *N*_1_, *M*_1_ using the equations 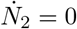 and 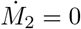:

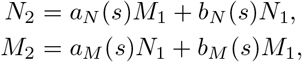

with the coefficients

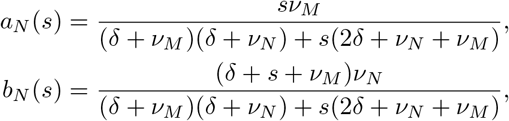

and *a_M_*(*s*), *b_M_*(*s*) follow by *N* ↔ *M*. This changes the the equations 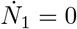 and 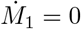 into

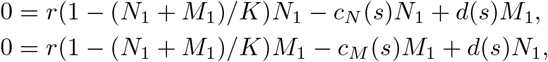

with coefficients

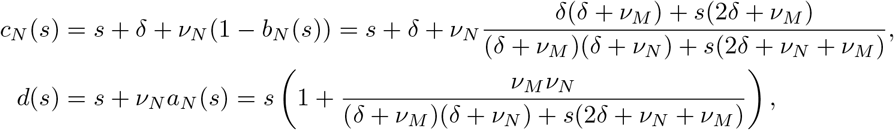

and *c_M_*(*s*) follows by *N* ↔ *M*. Multiplying the first equation by *M*_1_, the second equation by *N*_2_ and subtracting, we obtain

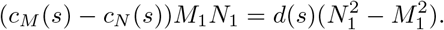

Therefore, either *N*_1_ = 0 or *N*_1_ ≠ 0. In the first case, we obtain the trivial fixed point by following back through the equations. In the latter case,

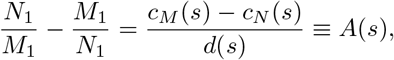

with

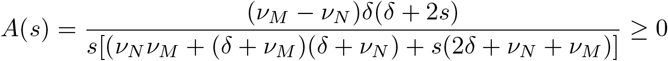

Therefore,

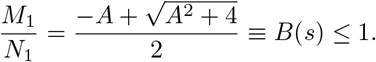

Finally, this result can be used to derive:

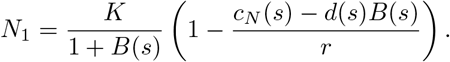

We also note that the symmetry *N* ↔ *M* is preserved as

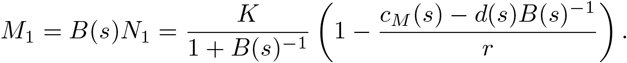

#### Corollary 3.

*The deadly motility regime for the source-sink model* (*L* = 2, *R* = 1) *is described as a subspace of* (*ν_N_*, *ν_M_, s*) *space such that*:

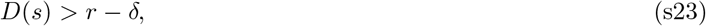

*with*

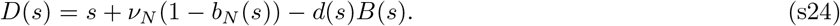

*D*(*s*) *has the following properties*:

- *D*(*s*) *is a strictly increasing smooth function of s* ∈ *R*^+^
- 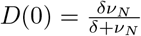.
- 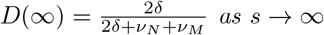 as *s* → ∞

*The deadly motility regime exist whenever the environment is sink-like* 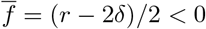, *and*

- *the slower phenotype is in the deadly motility regime* min(*ν_N_, v_M_*) > *ν_c_, and switching s is arbitrary, or*
- *the average phenotype is in the deadly motility regime* (*ν_N_* + *ν_M_*)/2 > *ν_c_, and switching s is above some critical value s* > *s_c_*(*ν_N_, v_M_*).

*In particular, the critical surface asymptotes with* min(*ν_N_, v_M_*) = *ν_c_ at low switching* (*s* ≪ *δ*) *and with* (*ν_N_ + v_M_*)/2 = *ν_c_ at high switching* (*s* ≫ *δ*).

*Proof.* From Theorem 6, the deadly motility regime is implicitly described 0 > *r* – *c_N_*(*s*) + *d*(*s*)*B*(*s*). Using *c_N_*(*s*) = *s* + *δ* + *ν_N_*(1 – *b_N_*(*s*), we get the condition in terms of *D*(*s*). Notice that *b_N_*(*s*), *d*(*s*) and *B*(*s*) are all smooth strictly decreasing functions of *s* ∈ ℝ^+^, hence *D*(*s*) is smooth strictly increasing. The limits follow using the formulae in the Theorem 6. Now, fix *ν_N_, v_M_* and vary *s*.

First, we are in the deadly motility regime irrespective of switching *s* if *D*(0) > *r* – *δ*. Notice that this condition is equivalent to 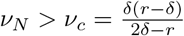, where we recognise the critical motility *ν_c_* in Theorem 1.

Second, we can reach the deadly motility regime for sufficiently high switching rates *s* > *s_c_*, where *s_c_* is the unique critical switching *s_c_* > 0 given by *D*(*s_c_*) = *r* – *δ*. Such a critical switching rate exists precisely when *D*(∞) > *r* – *δ*. This happens precisely when 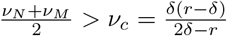.

#### Theorem 7.

*In the low motility combination v_N_, v_M_* ≪ *δ, the non-trivial fixed point of the staircase model is given by*:

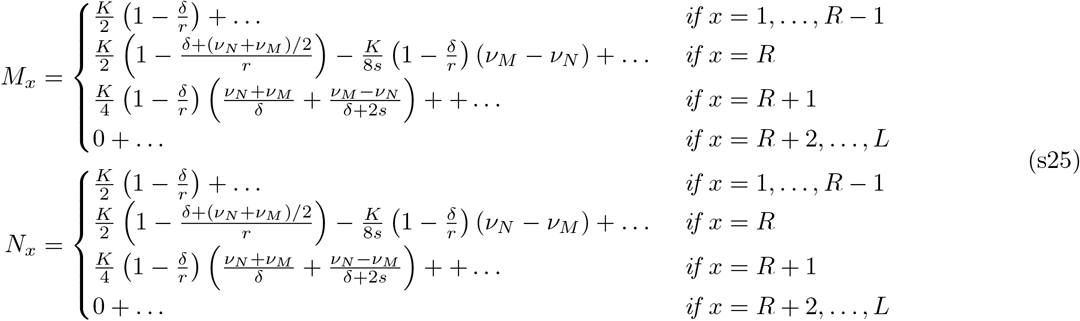

*In the high motility combination v_N_, v_M_* ≫ *δ, the non-trivial fixed point of the staircase model is given by*:

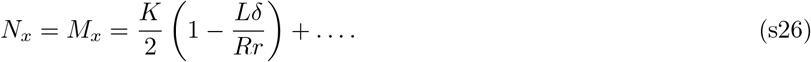

*In the mixed motility combination v_N_ ≪ *δ* ≪ *ν_M_*, the non-trivial fixed point of the staircase model is given by*

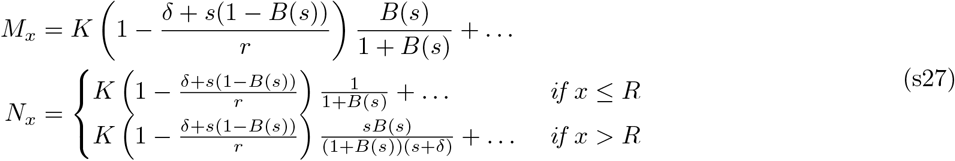

*with*

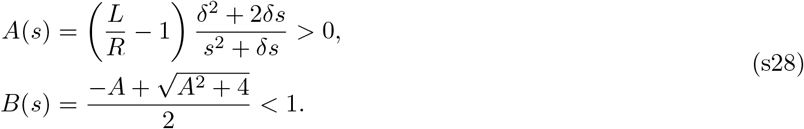

*Importantly, to lowest order N_x_/M_x_* = *B*(*s*) *in the region above the staircase x* ≥ *R*.

*Proof.* **Low motility combination.** Assume that *ν_N_, v_M_* are two independent small parameters, 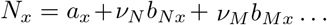, and 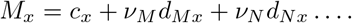 As before, expand in dimensional powers of 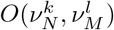 to simplify the algebra.

At *O*(1), the equations describe the local compartmental equilibrium:

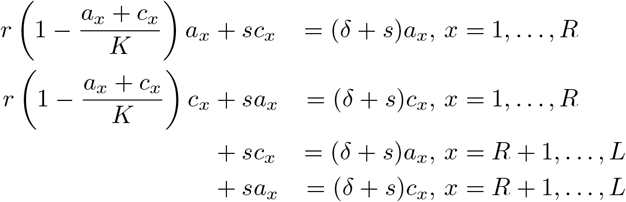

Subtracting the first two and the latter two equations, we can see that

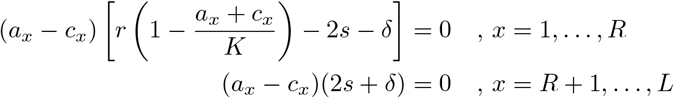

For *x* > *R*, this implies that *a_x_* = *c_x_*, and in turn that *a_x_* = 0 = *c_x_*. For *x* ≤ *R*, one of the brackets must vanish. If the latter bracket was to vanishes, then the original equations imply that *a_x_* + *c_x_* = 0. This is impossible as *a_x_,c_x_* > 0. Therefore, the first bracket vanishes, *a_x_* = *c_x_* and the original equations imply that

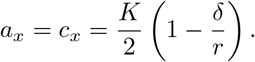

For further use, call this constant 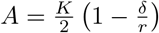. To summarise,

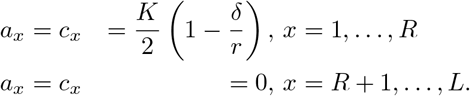

At *O*(*ν_N_*), the equations give the first order corrections to curvature as

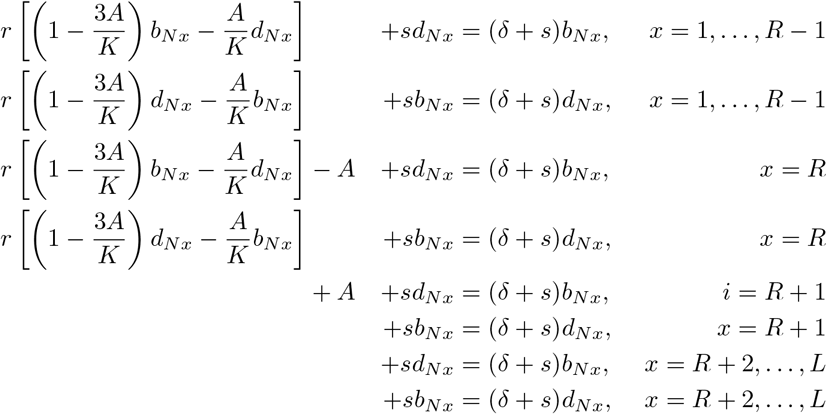

Using the subtraction as before, it can be seen that *b_Nx_* = *d_Nx_* = 0 unless *x* = *R* or *R* +1. By solving the appropriate system of linear equations for *x* = *R, R* + 1, we also see that

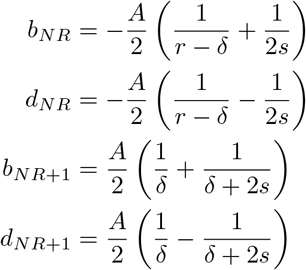

At *O*(*ν_M_*), the equations are the same up to swapping *M* with *N*, and *b* with *d*.

**High motility combination.** To simplify the algebra, assume a large single parameter *ν* such that *ν_N_, v_M_* ~ *O*(*ν*), denote the orders *ν^k^* as *O*(*k*) and expand *N_x_* = *a_x_* + *b_x_*/*ν* + …, and *M_x_* = *c_x_* + *d_x_*/*ν*…

At *O*(–1), the equations describe the motility equilibrium by

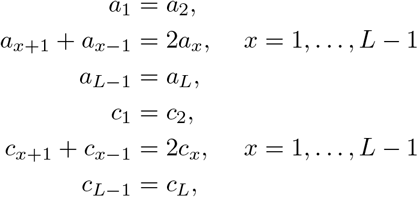

which imply that *a_x_* = *a* and *c_x_* = *c* for some *a, c* constants.

At *O*(0), the equations describe the first order correction to curvature due to local compartmental effects by

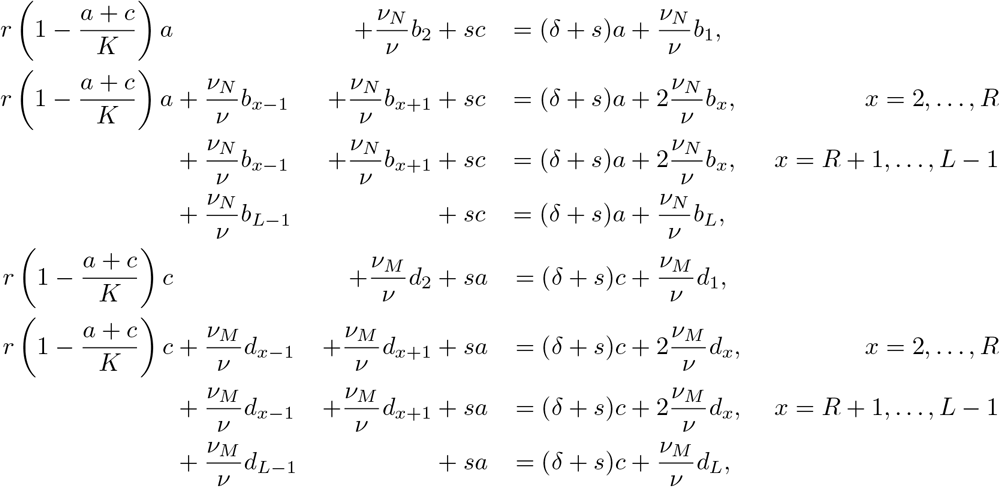

Summing the first and later four equations, we get

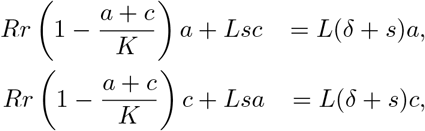

Multiplying the first equation by *c* and the latter by a, and subtracting, we get *Ls*(*c*^2^ – *a*^2^) = 0. Therefore, *c* = *a*. Then, we get

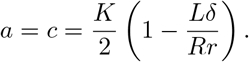

**Mixed motility combination.** Assume that there is a small *ν* such that *ν_N_* ~ *O*(*ν*) and *ν_M_* ~ *O*(1/*ν*). Then, *N_x_* = *a_x_* + *b_x_ν* + …, and *M_x_* = *c_x_* + *d_x_ν* ….

At *O*(–1), the equations for high motility equilibrium are:

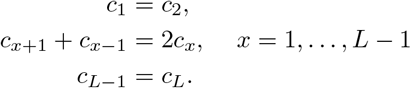

This result implies that there is a constant *c* such that *c_x_* = *c*.

At *O*(0), the local dynamics of the slow population involves the faster population:

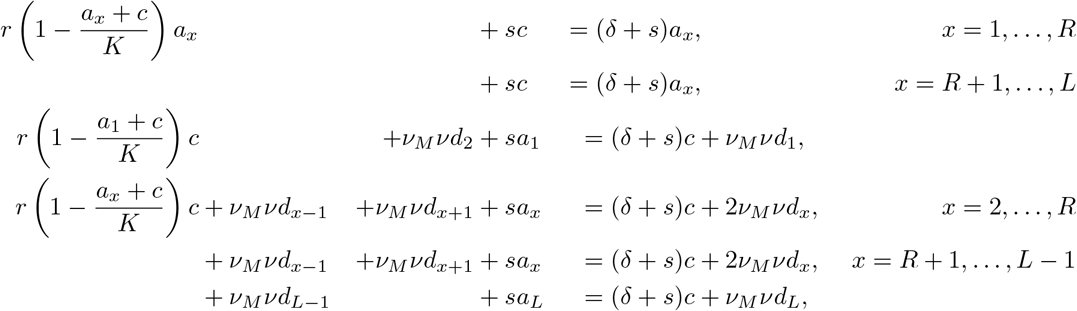

Notice that the first equation can be thought of as a quadratic equation for *a_x_* when *c* is thought of as a parameter:

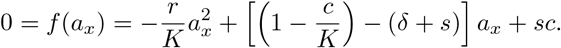

Since –*r/K* < 0 and *f* (0) = *sc* > 0, the parabola has exactly one root with *a_x_* > 0. Therefore, *a_x_* = *α* when *x* ≤ *R*. The second equation similarly gives that *a_x_* = *b* when *x* > *R*.

By summing the other equations, we get the system for variables *α, b, c*:

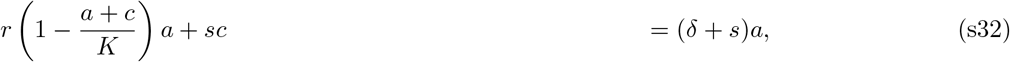

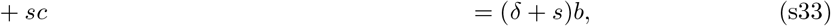

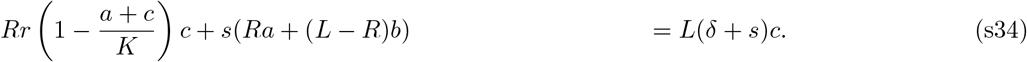

Firstly, eliminate *b* from the equation (s34) to get

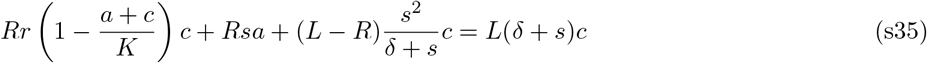

Multiplying equation (s35) by *a* and subtracting it from the equation (s32) multiplied by *Rc*, we can obtain

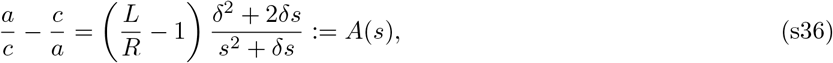

which defines a dimensionless number *A*(*s*) > 0. The equation for *c*/*α* can be solved by requiring that *c*/*α* > 0:

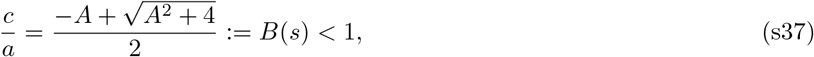

where *B*(*s*) is the proportion of high motility phenotype to low motility phenotype above the staircase. Combining (s32) and (s37), it is straightforward to find the solution

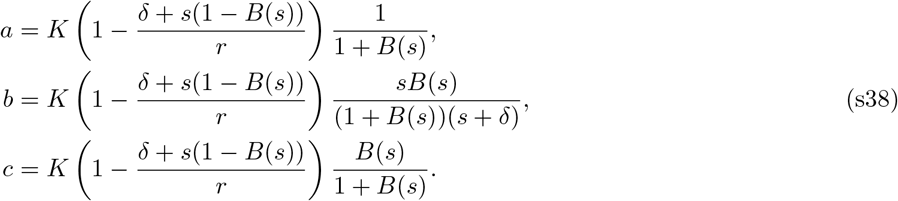

#### Theorem 8.

*At low switching s* ≪ *δ*, *the overall wild-type profile behaves as if it has a single motility of v* = min(*ν_N_, v_M_*). *At high switching s* ≫ *δ, the overall wild-type profile behaves as if it has a single motility of v* = (*ν_N_ + v_M_*)/2.

*In particular, this result affects the deadly motility regime in the* (*ν_N_, v_M_, s*) *space. Let* 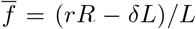 *be the average wild-type fitness. The wild-type profile is non-trivial iff*:

- *the environment is source-like* 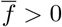, *motility rates v_N_, v_M_ are arbitrary, and switching rate s is arbitrary*,
- *the environment is sink-like* 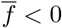 *and*

– *switching is low s* ≪ *δ and the slower phenotype is in the deadly motility regime* min(*ν_N_, v_M_*) > *ν_c_*,
– *switching is high s* ≫ *δ and the average-motility phenotype is in the deadly motility regime* (*ν_N_* + *ν_M_*)/2 > *ν_c_*.

*Proof.* As for the source-sink model, the deadly motility regime corresponds to region in the (*ν_N_, v_M_, s*) space. Now, we will fix the motility rates *ν_N_, v_M_* and vary *s*.

At high switching (*s* ≫ *δ*), the dominant dynamics 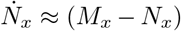 (and *N_x_* ↔ *M_x_*) pushes the dynamics close to the surface *N_x_* = *M_x_*. On this surface, we can define coordinates 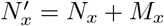 and derive the dynamics

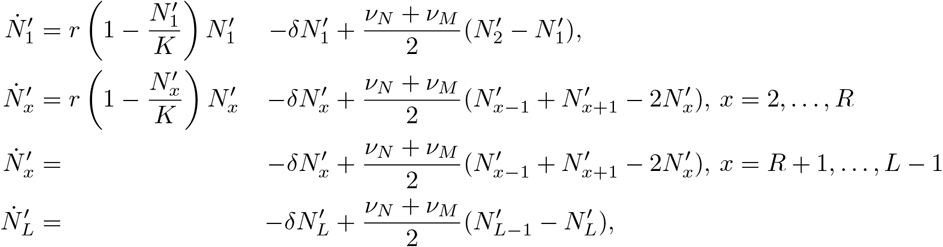

Therefore, the overall population 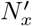 behaves as a population with average motility. In particular, the overall population is in the deadly motility regime at high switching (*s* ≫ *δ*) iff the average-motility population is.

For low switching, we prove the stability of the non-trivial fixed point at *s* = 0. This fixed point is given by *M_x_* = 0 and *N_x_*, which coincides with the non-trivial fixed point of the staircase model for a single phenotype of motility *ν* = *ν_N_* (s6). The corresponding Jacobian can be derived from (s16) and has the form

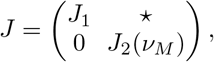

where

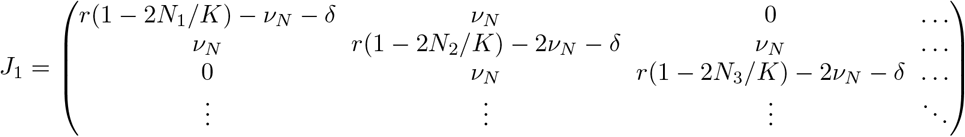

and

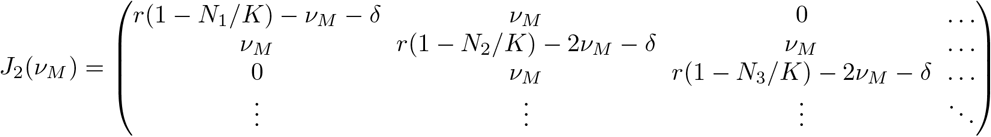

We prove the stability of the non-trivial fixed point if we show that the eigenvalues of *J* have negative real part. Notice that the set of eigenvalues of *J* is the union of the sets of eigenvalues of *J*_1_ and *J*_2_(*ν_M_*). Therefore, it suffices to show that the eigenvalues of *J*_1_ and *J*_2_(*ν_M_*) are negative. The eigenvalues of Ji must be negative since *J*_1_ is the Jacobian corresponding to the non-trivial fixed point of the staircase model for a single phenotype with motility *ν_N_*, which is stable from previous analysis. The proof that the eigenvalues of *J*_2_(*ν_M_*) are negative for *ν_M_* > *ν_N_* is identical to the proof [S18], which consists of two steps. First, one shows that the largest eigenvalue of *J*_2_(*ν_N_*) is 0 because (*N*_1_,…, *N_L_*) is the eigenvector and only the largest eigenvalue can have an eigenvector with positive entries. Second, one shows that this eigenvalue increases as *ν_M_* increases from *ν_N_*. To sum up, the overall population is dominated by the slower population *N_x_* at low switching (*s* ≪ *δ*).

The result for the existence of the deadly motility regime follows, as the system behaves as a single phenotype of single motility which interpolates between the slower motility min(*ν_N_*, *ν_M_*) (at low switching *s* ≪ *δ*) and the average motility (*ν_N_* + *ν_M_*)/2 (at high switching *s* ≫ *δ*).

### S8. Figure Notes

This section aims to provide the parameter values used for producing all our figures as well as other specific guidance for reproducing our results fully. Unless stated otherwise, we follow previous work [S7] and use as lattice size *L* = 8, carrying capacity *K* = 10^5^, mutation rates *μ_f_* = 10^-7^ and *μ_b_* = 10^-4^, division rate *r* =1 and death rate *δ* = 0.1 (resp. *δ* = 0.3) for source-like (resp. sink-like) environments.

Specifically, in Fig. 1, the death rate is *δ* = 0.1. Panel *b* was created from a simulation of the staircase model with *ν* = 1*e* – 2 for low motility and *ν* = 1 for high motility. Panel *d* was created from the same simulations. Moreover, the black curve describing the total population numbers was plotted by finding the non-trivial fixed point of the mean-field theory (s6) via the Newton-Raphson method.

In Fig. 2, motility rate *ν* is varied in logarithmic steps between 10^-5^ and 10, and the appropriate death rate *δ* = 0.1, 0.3 is chosen for the source-like and sink-like environments, respectively. The dots of panel *b* correspond to simulations while the lines correspond to the analytical techniques. Panel *c* was produced by the analytical techniques. Panel *d* was plotted by finding the non-trivial fixed point of the mean-field theory (s6) via the Newton-Raphson method.

In Fig. 3, the death rate is *δ* = 0.3 and the switching rates s are *s* = 0 (low switching *s* ≪ *δ*) and *s* = 5 (high switching *s* ≫ *δ*). In panel *b*, motility rates *ν*_1,2_ are varied in logarithmic steps between 10^-5^ and 10 and the resulting heatmap plots of the adaptation rate are constructed for resistance state *R* = 5 (Fig. 1b) from simulations, where *R* is the genotype of highest resistance before the adaptation jump *R* → *R* + 1. In panel *c*, motility combinations are represented by (*ν*_1_, *ν*_2_) = (10^-2^,10^-4^) (low motility), (*ν*_1_, *ν*_2_) = (3.16,10^-4^) (mixed motility) and (*ν*_1_, *ν*_2_) = (3.16,1) (high motility). Wild-type profiles are plotted for resistance state *R* = 4 from simulations. Black curves describe the total number of cells in each phenotype and were plotted by finding the non-trivial fixed point of the mean-field theory s39 via the Newton-Raphson method,

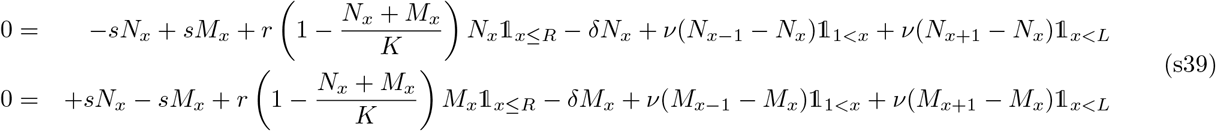

In panel b of Fig. 4, the death rate is *δ* = 0.3 and switching thresholds *S* are *S* = 9.5 × 10^4^ (high threshold *S* ≫ *K*(1 – *δ/r*)) and *S* = 100 (low threshold *S* ≪ *K*(1 – *δ/r*)). Motility rates *ν_L,H_* are varied in logarithmic steps between 10^-5^ and 10 and the resulting heatmap plots of the adaptation rate are constructed for resistance state *R* =2 from simulations. In panel *c*, the death rate is *δ* = 0.1 and switching thresholds *S* are *S* = 9.5 × 10^4^ (high threshold *S* ≫ *K*(1 – *δ/r*)) and *S* = 10^4.5^ (low threshold *S* ≪ *K*(1 – *δ/r*)). This choice is different from the one used in panel *b* but allows for better visualisation of the wild-type profiles when the scale of cell numbers is kept linear as in previous figures. The mixed motility combination is represented by (*ν_L_, v_H_*) = (3.16,10^-4^) (fast-to-slow) and (*ν_L_,v_H_*) = (10^-4^, 3.16) (slow-to-fast). The wild-type profiles (including the black curves) are plotted for resistance state *R* = 1 from simulations. The Newton-Raphson method is not reliable in this case because the step-function used for density-dependent motility is not smooth.

In Fig. 5, we consider *L* = 2 and *R* = 1. *K* = 10^5^ and *r* = 1 are chosen as usual and mutation rates do not need not be chosen as they do not affect the dynamics on ecological time-scales. In panel b, the phase portraits are plotted with matplotlib [S19] for the choice of the death rate *δ* = 0.25 and the motility rates *ν* = 0, 0.1, 100 (source-like environment), resp. for the death rate *δ* = 0.75 and the motility rates *ν* = 0, 0.1, 5 (sink-like environment). In panel *c*, the bifurcation diagrams are plotted using the solution for the non-trivial fixed point (s1). The source-like environment is represented by *δ* = 0.25 and the sink-like environment is represented by *δ* = 0.75.

In Fig. S1, the death rate *δ* = 0.1, 0.3 is chosen for source-like and sink-like environment, respectively. Motility rate *ν* is varied in logarithmic steps between 10^-5^ and 10 and resistance cost *c* is varied in logarithmic steps between 10^-4.5^ and 10^-0.5^. The resulting heatmap plots of adaptation rate are constructed for resistance state *R* = 3 from simulations.

In Fig. S2, the death rate is *δ* = 0.1. Motility rate *ν* is varied in logarithmic steps between 10^-5^ and 10 and the multiplicative increase in death rate *σ* is varied linearly between 1 and 9. The resulting heatmap plots of the adaptation rate are constructed for resistance state *R* = 5 from simulations.

In Fig. S3, the death rate is *δ* = 0.1. Motility rate *ν* is varied in logarithmic steps between 10^-5^ and 10 and the chemotactic probability to move up the antibiotic gradient *p* is varied linearly between 0.1 and 0.9. The resulting heatmap plots of the adaptation rate are constructed for resistance state *R* = 5 from simulations.

In panel a of Fig. S4 the switching rate is varied in logarithmic steps between 10^-5^ and 10. The mixed motility combination is chosen as (*ν*_1_, *ν*_2_) = (10^-3^,10) and the appropriate death rate δ = 0.1, 0.3 is chosen for the source-like and sink-like environment, respectively. The dots of panel b correspond to simulations while the lines are obtained by analytical techniques for a population of homogeneous motility min(*ν*_1_, *ν*_2_) at low switching and (*ν*_1_ + *ν*_2_)/2 at high switching. In panel *b*, the death rate is *δ* = 0.1 and the switching rate is varied *s* = 0, 0.001, 0.01, 0.1, 1. Motility rates *ν*_1,2_ are varied in logarithmic steps between 10^-5^ and 10 and the resulting heatmap plots of the adaptation rate are constructed for resistance state *R* =5 from simulations. In panel c, the death rate is *δ* = 0.3 and the switching rate is varied *s* = 0.1, 0.4, 0.67, 0.9, 1, 10. Motility rates *ν*_1,2_ are varied in logarithmic steps between *ν* = 10^-5^ and *ν* = 10. The survival probability is computed as follows. We run the Newton-Raphson method to search for a fixed point of (s39) in the non-negative quadrant *N_x_, M_x_* ≥ 0. If the search returns the trivial (resp. non-trivial) fixed point, we set the survival probability to 0 (resp. 1).

In Fig. S5, the death rate is *δ* = 0.1 and switching thresholds S are *S* = 9.5 × 10^4^ (high threshold *S* ≫ *K*(1 – *δ*/*r*)) and *S* = 100 (low threshold *S* ≪ *K*(1 – *δ/r*)). Motility rates *ν_L,H_* are varied in logarithmic steps between 10^-5^ and 10 and the resulting heatmap plots of the adaptation rate are constructed for resistance state *R* = 2 from simulations.

In Fig. S6, the death rate *δ* = 0.1, 0.3 is chosen for the source-like and sink-like environment, respectively. Motility rate *ν* is varied in logarithmic steps between 10^-5^ and 10 and the horizontal gene transfer rate *h* is in logarithmic steps between 10^-7^ and 10^-3^. The resulting heatmap plots of the adaptation rate are constructed for resistance state *R* = 4 from simulations.

## SUPPLEMENTARY FIGURES

**FIG. S1.**
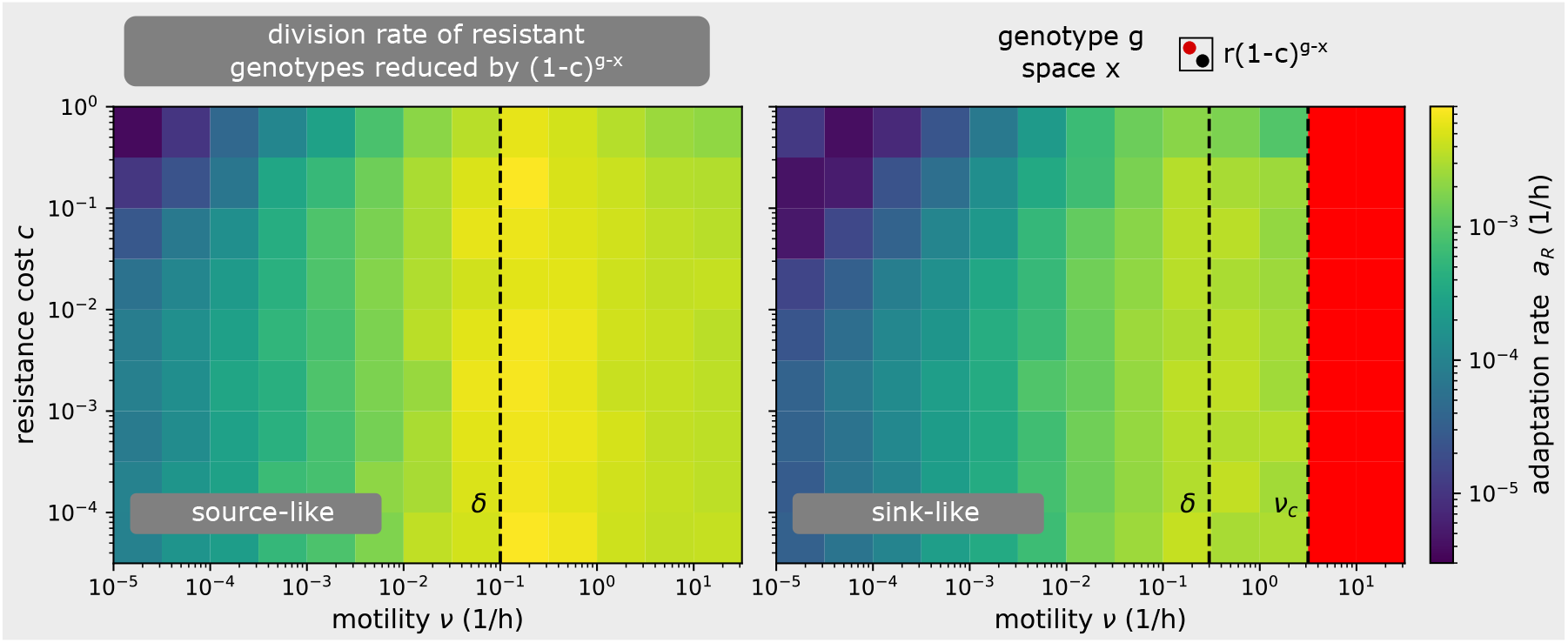
Effect of resistance costs. Resistance cost *c* is modelled as a reduction in the division rate of genotypes *g* at position *x* by a factor of (1 – *c*)^*g–x*^ as in [S7]. The heatmap of adaptation rate on the (*ν, c*) plane explains how resistance cost *c* changes the relationship between motility rate *ν* and adaptation rate *a_R_*. Generically, resistance cost does not affect the relationship between adaptation rate and motility: low motility accelerates adaptation while high motility decelerates adaptation. Only when the motility rate is very low *ν* ≪ *δ* and resistance cost is very high 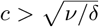, the adaptation rate is decreased by *c*, as proved in [S7]. However, such resistance costs are considered high [S8]. Even if the motility is low as *ν* = 10^-2^*δ* (i.e., 99% of bacteria do not migrate during their lifetime), the resistance cost is important only if *c* > 0.1 [S8]. Moreover, resistance cost cannot influence the deadly motility regime (red) as in this regime the susceptible wild-type goes extinct. A detailed description of this model can be found in SI Text S4.

**FIG. S2.**
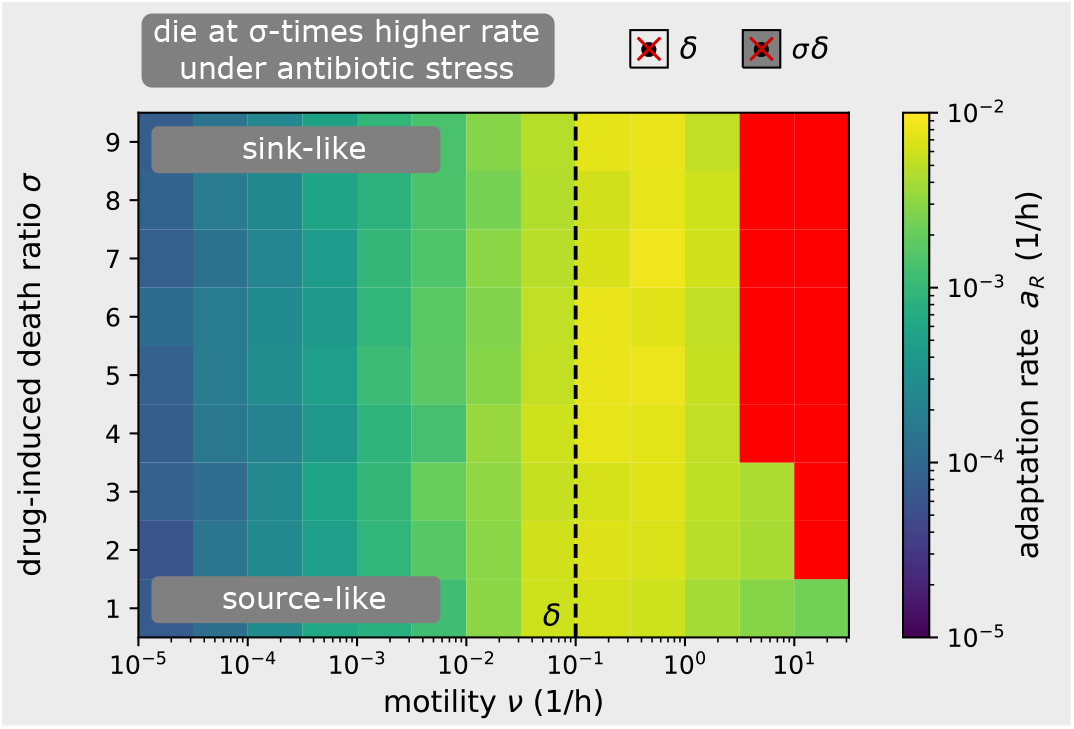
Effect of bactericidal antibiotics. Bactericidal antibiotics are considered by increasing the death rate of susceptible genotypes *g* < *x* (under the staircase) *σ*-times. The heatmap of adaptation rate on the (*ν, σ*) plane explains how *σ* changes the relationship between motility *ν* and adaptation rate *a_R_*. In the low motility regime *ν* < *δ*, the adaptation rate increases with motility *ν* and is unaffected by *σ*. In the high motility regime *ν* > *δ*, the adaptation rate decreases with motility *ν*. Moreover, even if the environment is source-like for *σ* = 1, the high motility regime can transition into the deadly motility regime (red) provided the bactericidal effect is strong enough *σ* ≫ 1. A detailed description of this model can be found in SI Text S4.

**FIG. S3.**
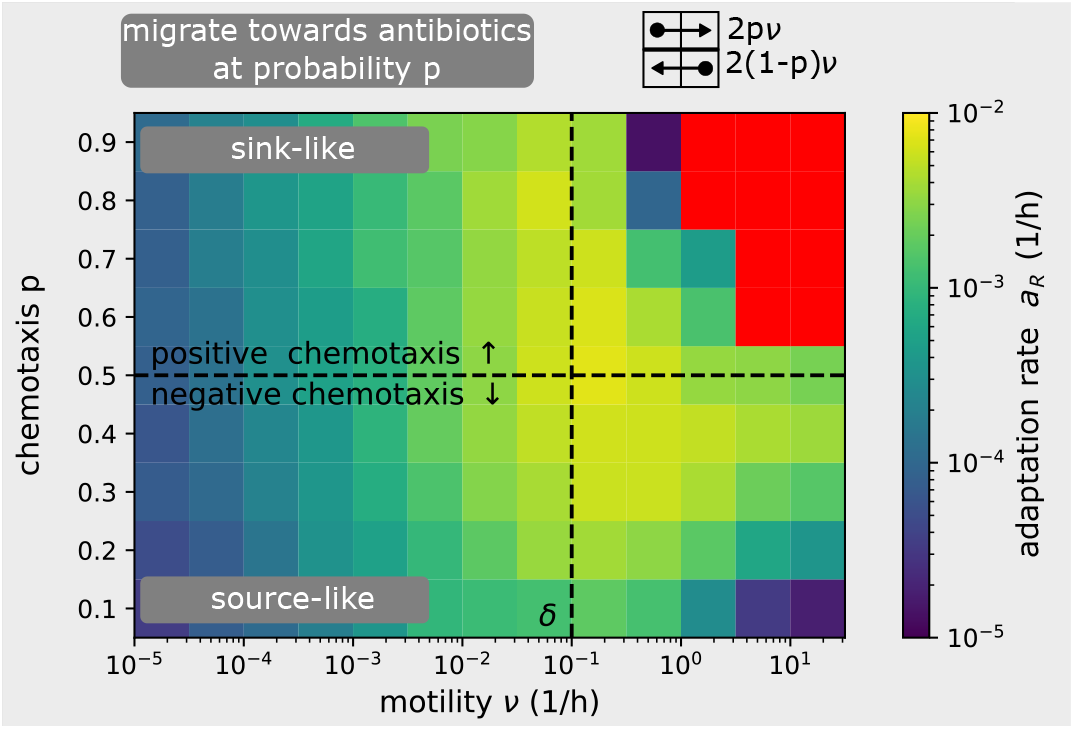
Effect of chemotaxis. Motility bias in the chemical gradient (chemotaxis) is considered by varying the probability *p* that a cell moves up or down the antibiotic gradient. In our original model *p* = 0.5. Cell movement is biased up the antibiotic gradient if *p* > 0.5 (positive chemotaxis), and down the gradient *p* < 0.5 (negative chemotaxis). The heatmap of adaptation rate on the (*ν, σ*) plane explains how *p* changes the relationship between motility *ν* and adaptation rate *a_R_*. In the low motility regime *ν* < *δ*, the adaptation rate increases with motility *ν* and is unaffected by chemotaxis *p*. In high motility regime *ν* > *δ*, positive chemotaxis *p* > 0.5 converts the high motility regime into the deadly motility regime even when the environment is source-like at *p* = 0.5. Negative chemotaxis *p* < 0.5 prevents the deadly motility regime and slows down the adaptation in the high motility regime, compared to unbiased motion *p* = 0.5. A detailed description of this model can be found in SI Text S4.

**FIG. S4.**
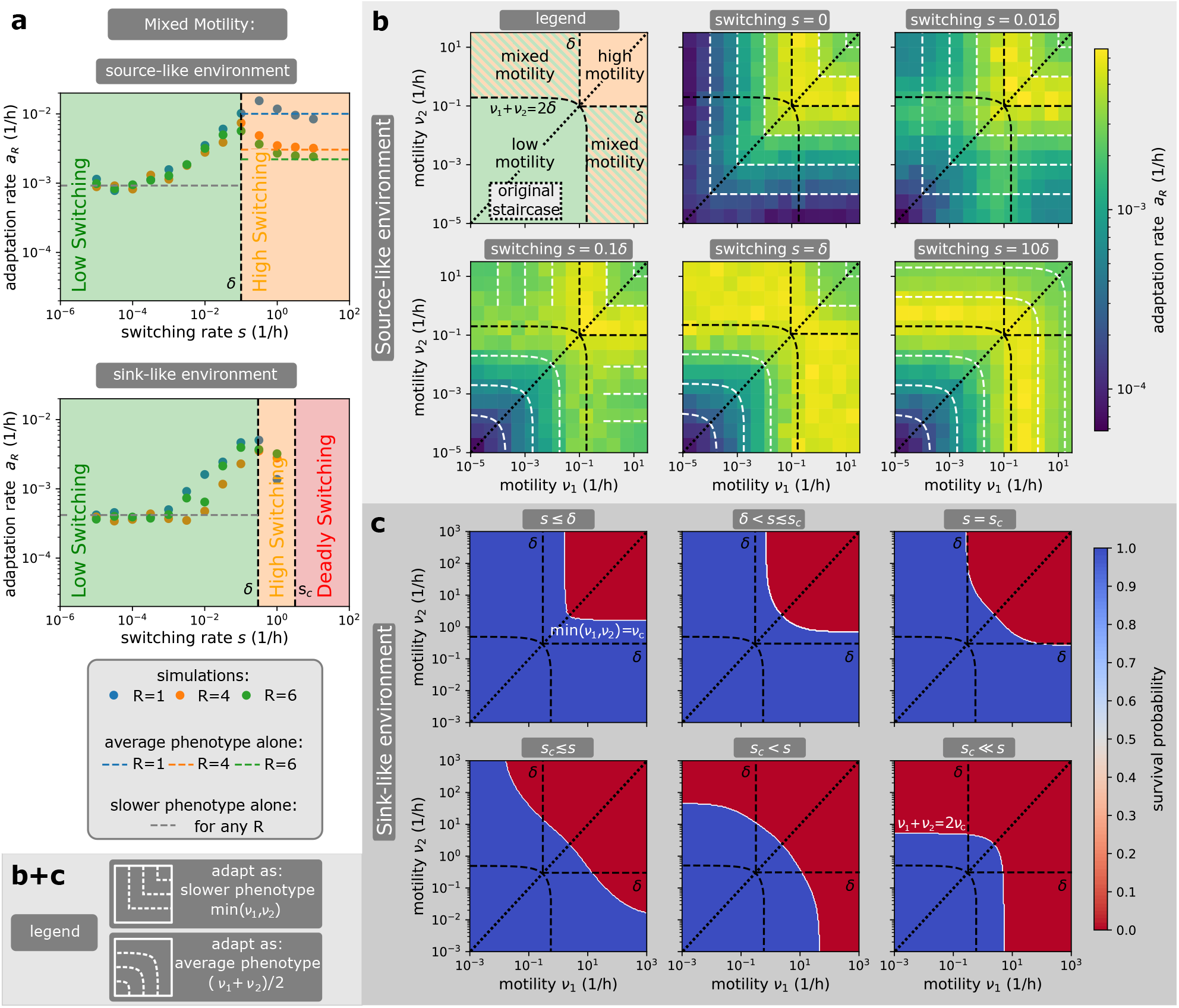
Further effects of switching between motility phenotypes stochastically. **a)** Switching rate *s* modifies the adaptation rate *a_R_* and the adaptation regime in the mixed motility combination *ν*_1_ ≪ *δ* ≪ *ν*_2_. Dots correspond to simulations, while the asymptotic lines are derived with analytical techniques. At low switching (*s* ≪ *δ*), the population adapts as if a single slower phenotype is present, whose adaptation rate is independent of the resistance state *R* (grey dashed line). At high switching (*s* ≪ *δ*), the population behaves as if a single phenotype of average motility is present, whose adaptation rate depends on the number of compartments *R* where wild-type can divide (coloured dashed lines). Moreover, if the environment is sink-like and the average motility is above the critical motility, there is a critical switching rate *s_c_* which limits bacterial survival. **b)** Adaptation rate heatmap on the (*ν*_1_,*ν*_2_) plane for different switching rates *s*. This is the same plot as in Fig. 3b but for a source-like environment instead of a sink-like environment. The (*ν*_1_, *ν*_2_) plane can be partitioned into different combinations of adaptation regimes and its diagonal corresponds to a population of a single motility. Bacteria adapt as if all cells had the same effective motility, which corresponds to the intersection of the level sets (white dashed lines) with the diagonal. At low switching rate *s*, the effective motility matches the slower motility phenotype present min(*ν*_1_, *ν*_2_). At high switching rate *s*, the effective motility matches the average motility (*ν*_1_ + *ν*_2_)/2. Unlike for the sink-like environment, there is no deadly motility regime for any motility *ν*_1,2_ and switching *s* in the source-like environment. **c)** Deadly motility regime and critical surface. In sink-like environments, there is an abrupt change in survival probability from 1 to 0 when the population is sufficiently well-mixed. The deadly motility regime corresponds to the space with zero survival probability (red) and is bounded by a critical surface (white). The critical surface asymptotes with min(*ν*_1_, *ν*_2_) = *ν_c_* at low switching (*s* ≪ *δ*) and with (*ν*_1_ + *ν*_2_)/2 = *ν_c_* at high switching (*s* ≫ *δ*).

**FIG. S5.**
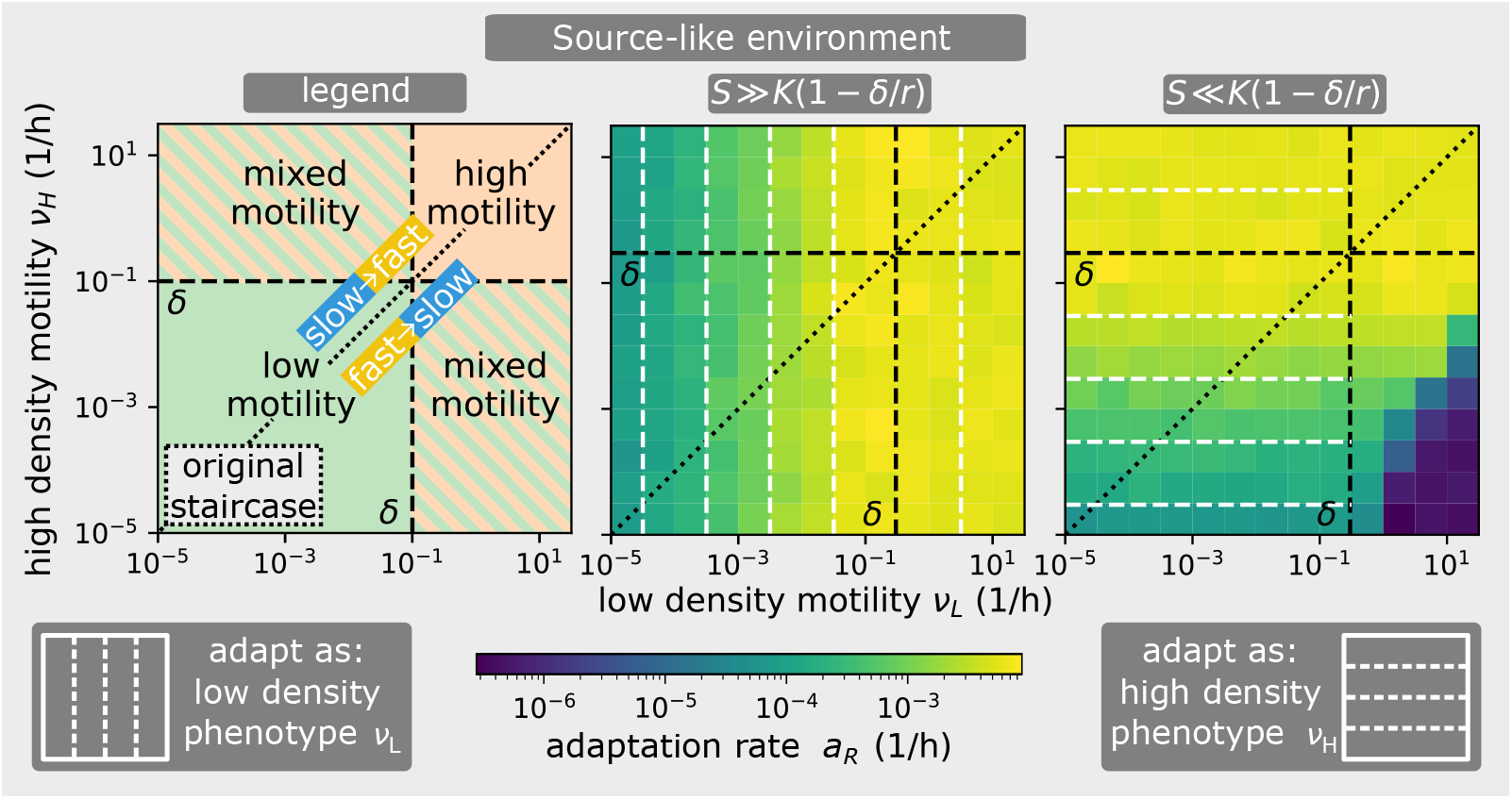
Further effects of density-dependent motility. Adaptation rate heatmap on the (*ν_L_, v_H_*) plane for different switching thresholds *S* in the source-like environment, where *ν_L,H_* are the low-density/high-density motility rates (Fig. 4a). This is the same plot as in Fig. 4b but for a source-like environment instead of a sink-like environment. The (*ν_L_,v_H_*) plane can be partitioned into different combinations of adaptation regimes and its diagonal corresponds to a population of a single motility. The diagonal separates slow-to-fast and fast-to-slow switching combinations, which differ in relative motility at low-to-high density. Bacteria generically adapt as if all cells had the same effective motility, which corresponds to the intersection of the level sets (white dashed lines) with the diagonal. This effective motility generically matches the low-density (resp. high-density) motility at high (resp. low) threshold S. At low threshold S, this generic rule has one exception. The adaptation rate is reduced in the mixed motility combination of the fast-to-slow switching case. The reduction in adaptation rate in slow-to-fast switching in Fig. 4b is caused by the critical motility, which does not appear in the source-like environment.

**FIG. S6.**
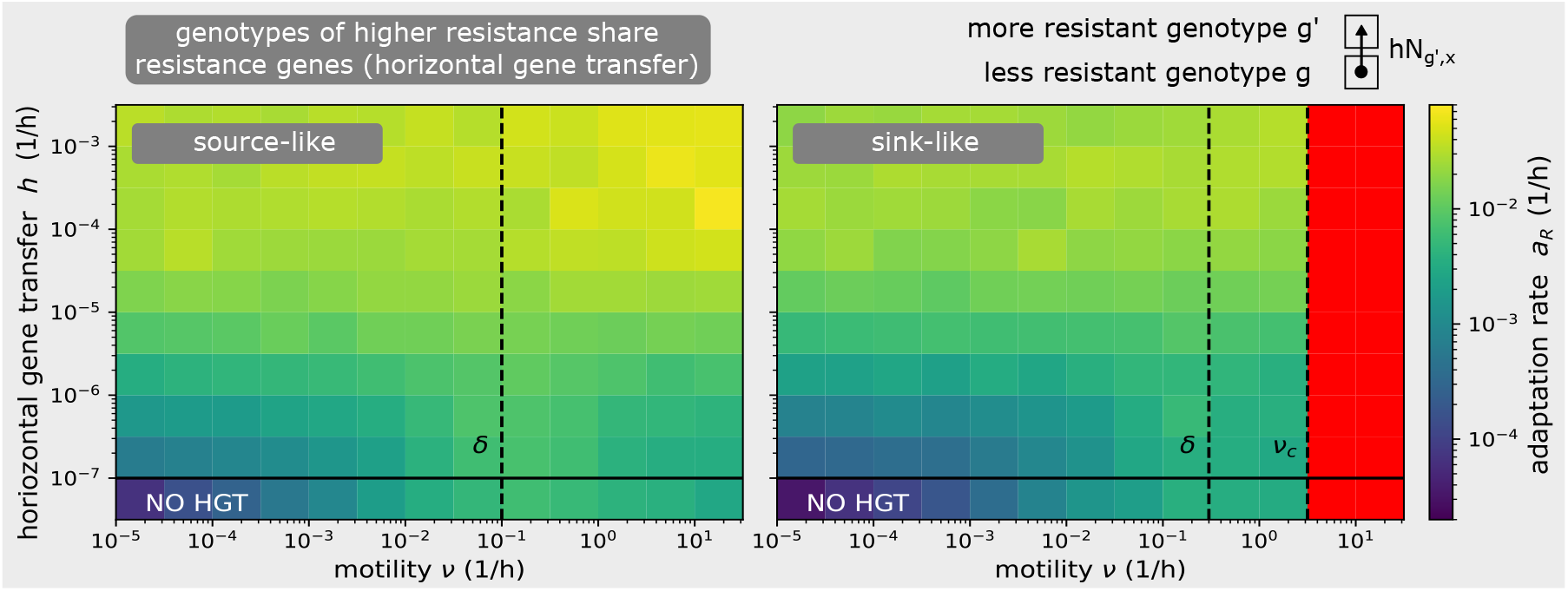
Effect of horizontal gene transfer (HGT). The heatmap of adaptation rate on the (*ν, h*) plane explains how the HGT rate *h* changes the relationship between motility *ν* and adaptation rate *a_R_*. HGT is modelled by allowing cells of less resistant genotypes g to acquire resistance from *N_x,g’_* more resistant cells of genotype *g*’ > *g* located at the same position *x* at a rate *h* × *N_x,g’_*. When HGT rate is low, the relationship between adaptation rate and motility is not significantly changed: low motility accelerates adaptation while high motility decelerates adaptation. When HGT rate is high, the adaptation rate is significantly increased and changes in motility do not play a significant role. However, the required HGT threshold *h* ≈ 10^-5^ corresponds to *hK/δ* ≈ 1 horizontal gene transfers per generation. Such HGT rates are considered very high, compared to the upper bound of 10^-4^ transfers per generation known from the literature [S12]. Moreover, HGT cannot influence the deadly motility regime (red) as the donors of resistance genes do not have enough time to form before the wild-type goes extinct. A detailed description of this model can be found in SI Text S4.

## SUPPLEMENTARY MOVIES

**MOVIE S1.**
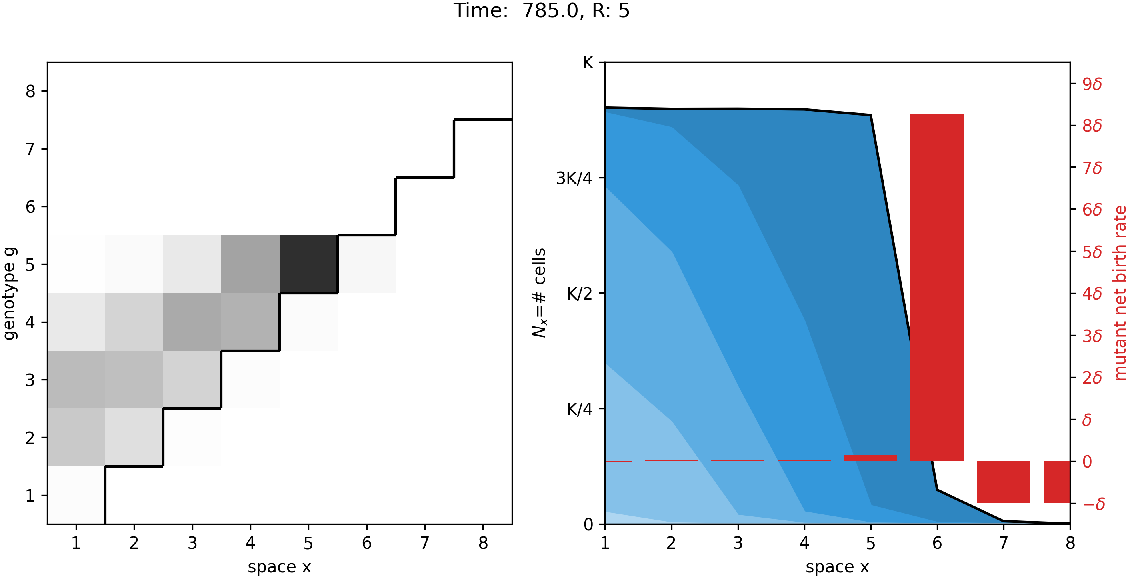
Bacterial adaptation in the low motility regime (*ν* < *δ*). Simulation of a bacterial population evolving antibiotic resistance in the staircase model in the low motility regime. It adapts and expands its range in a stepwise fashion, leaving behind an inclined comet tail (left) where shades of gray represent cell density in each compartment, which corresponds to a diversity of strains with different antibiotic susceptibility (right). Find a full description in the caption of Fig. 1. Parameters used: *L* = 8, *K* = 10^5^, *r* =1, *δ* = 0.1, *μ_f_* = 10^-7^, *μ_b_* = 10^-4^, *ν* = 0.01.

**MOVIE S2.**
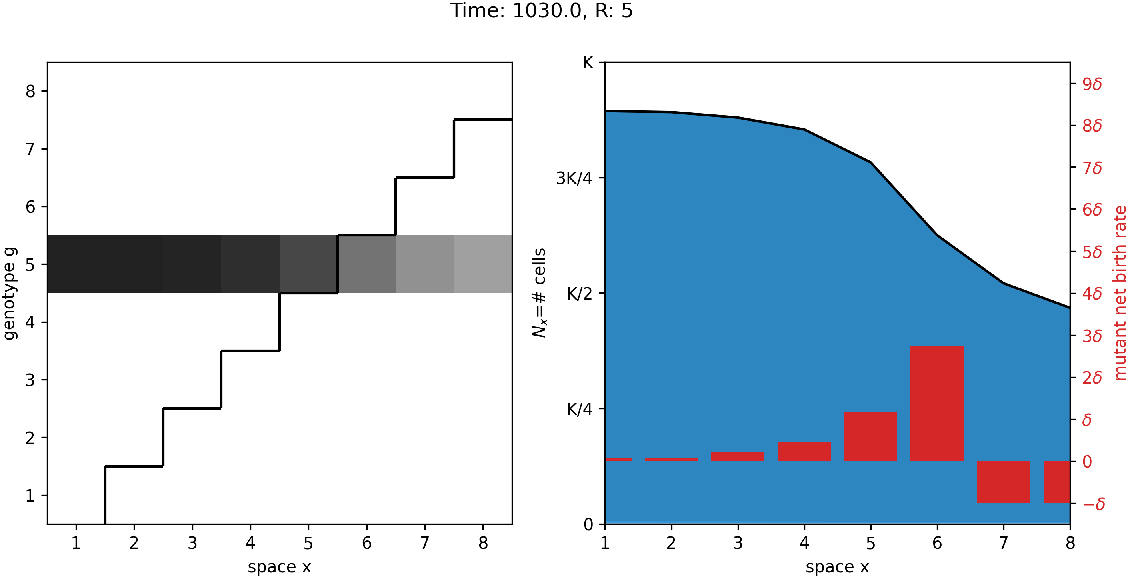
Bacterial adaptation in the high motility regime (*ν* > *δ*). Similar to Movie S1, but for the high motility regime. Note that the comet tail that was inclined in the low motility regime is now horizontal (left), and the population is made of a single strain that can grow in *x* ≤ *R* (right). Find a full description in the caption of Fig. 1. Parameters used: *L* = 8, *K* = 10^5^, *r* = 1, *δ* = 0.1, *μ_f_* = 10^-7^, *μ_b_* = 10^-4^, *ν* =1.

**MOVIE S3.**
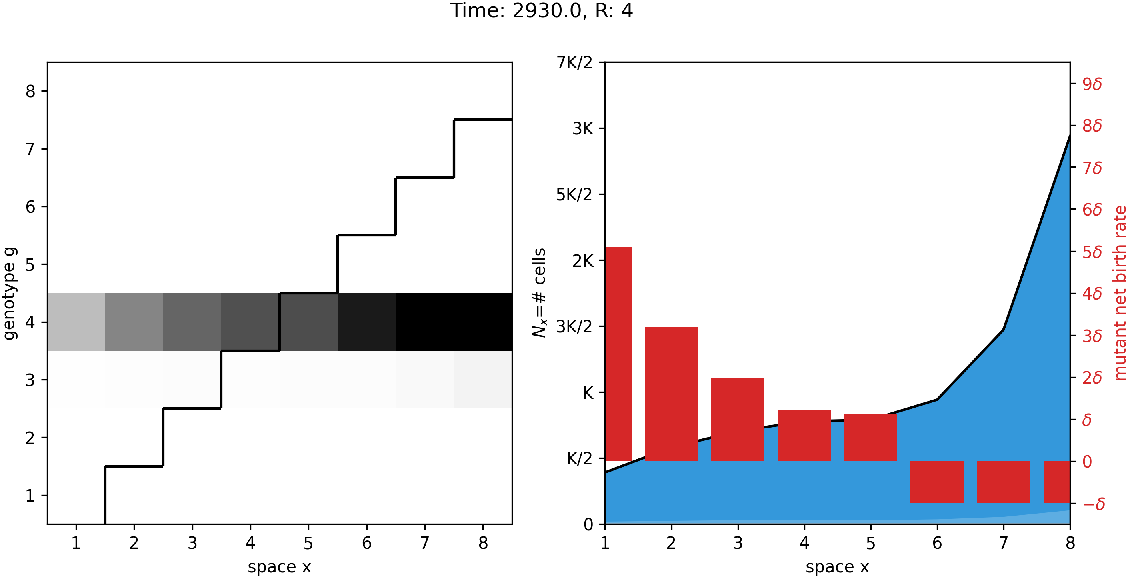
Effect of positive chemotaxis. Similar to Movie S2 but now *p* > 0.5 and the population survives. Parameters used: *L* = 8, *K* = 10^5^, *r* =1, *δ* = 0.1, *μ_f_* = 10^-7^, *μ_b_* = 10^-4^, *ν* =1, *p* = 0.7.

**MOVIE S4.**
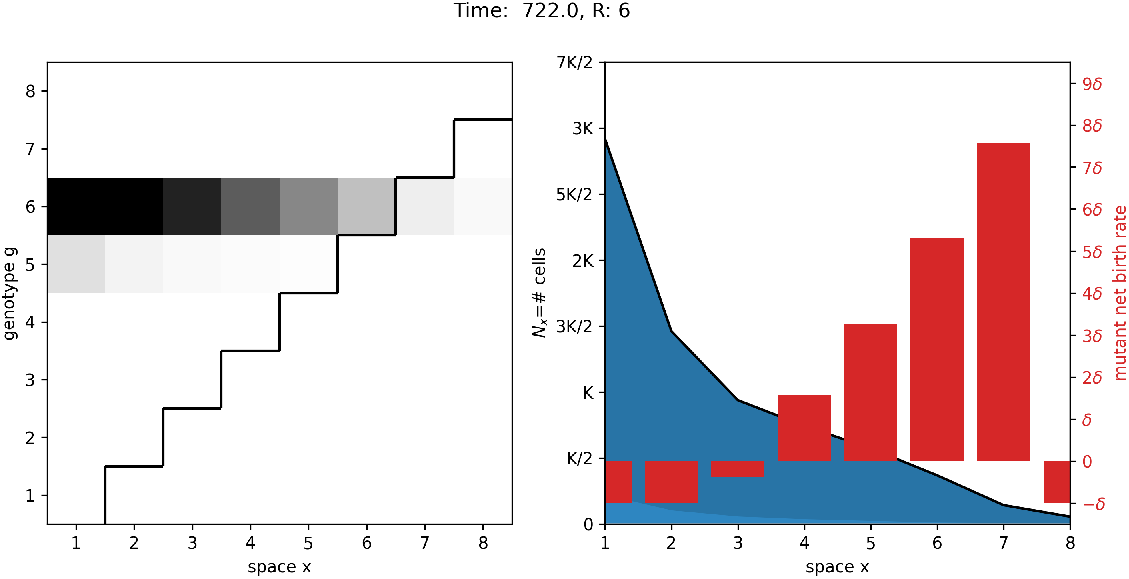
Effect of negative chemotaxis. Similar to Movie S2 but now *p* < 0.5. Parameters used: *L* = 8, *K* = 10^5^, *r* = 1, *δ* = 0.1, *μ_f_* = 10^-7^, *μ_b_* = 10^-4^, *ν* =1, *p* = 0.3.

**MOVIE S5.**
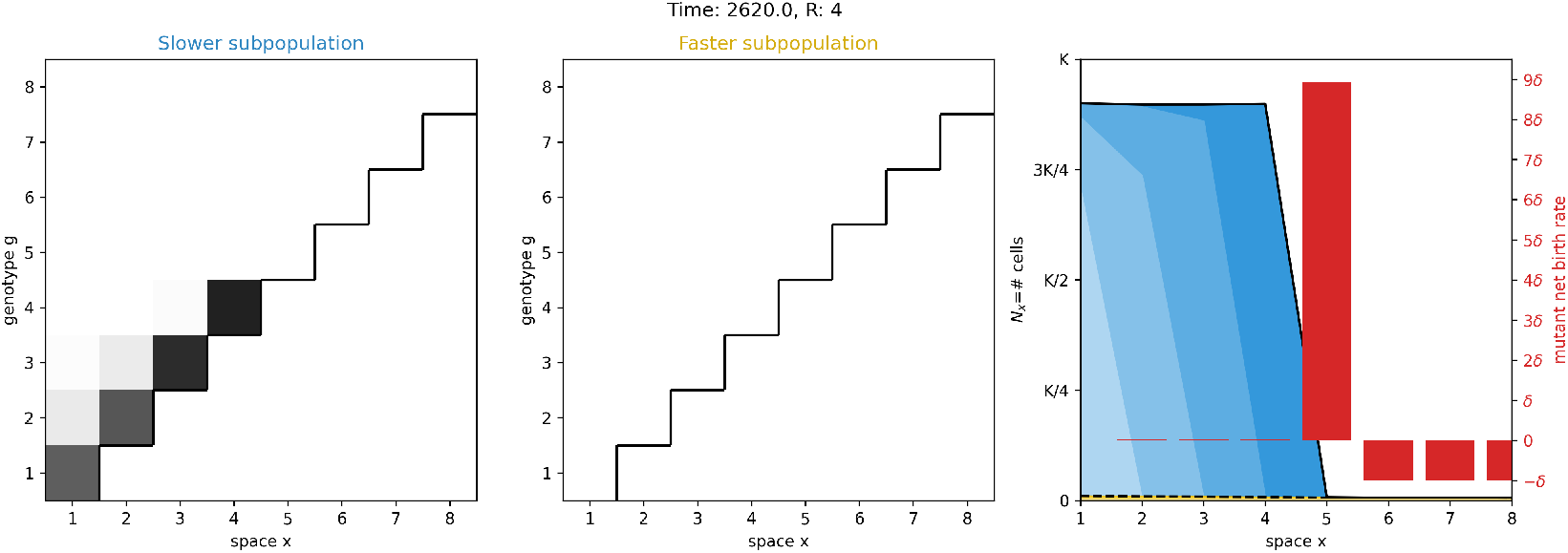
Effect of stochastic switching when the switching rate is low. Simulation of a bacterial population evolving antibiotic resistance in the staircase model with stochastic switching between motility phenotypes for the mixed motility combination min(*ν*_1_, *ν*_2_) < *δ* < (*ν*_1_ + *ν*_2_)/2. Find a full description in the caption of Fig. 3. Parameters used: *L* = 8, *K* = 10^5^, *r* = 1, *δ* = 0.1, *μ_f_* = 10^-7^, *μ_b_* = 10^-4^, *ν*_1_ = 10^-4^, *ν*_2_ = 10^0.5^, *s* = 10^-3^.

**MOVIE S6.**
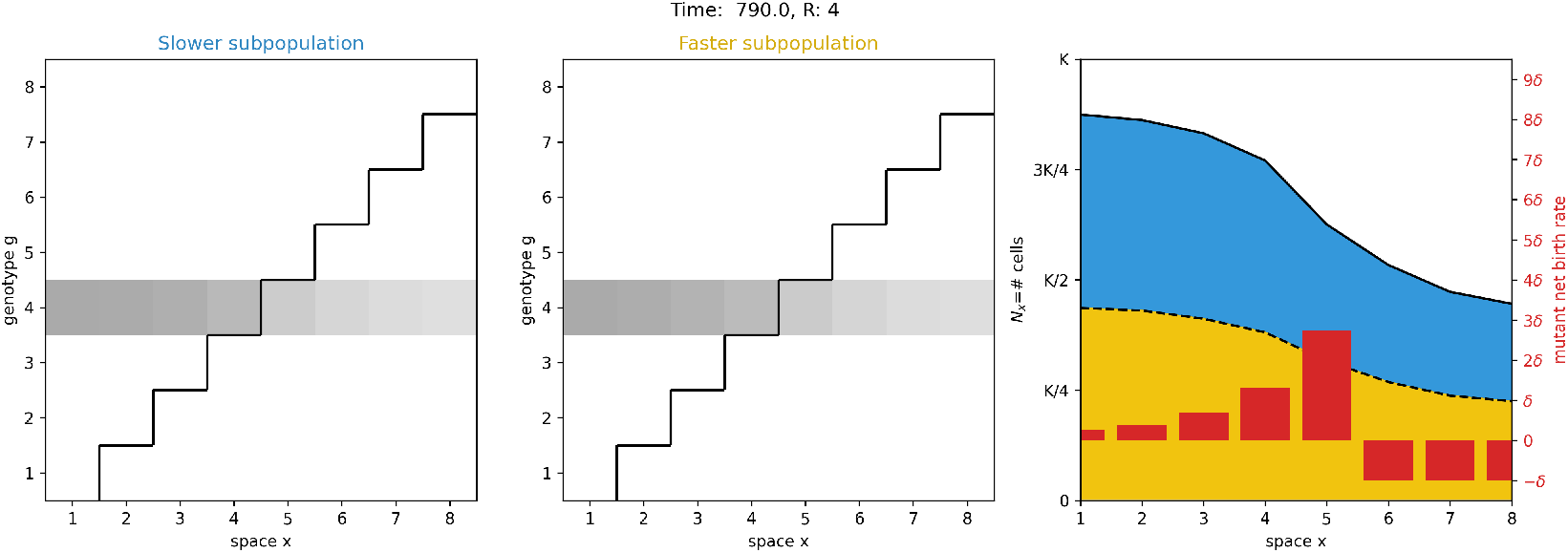
Effect of stochastic switching when the switching rate is high. Similar to Movie S5 but with high rates of switching between motility phenotypes. Find a full description in the caption of Fig. 3. Parameters used: *L* = 8, *K* = 10^5^, *r* =1, *δ* = 0.1, *μ_f_* = 10^-7^, *μ_b_* = 10^-4^, *ν*_1_ = 10^-4^, *ν*_2_ = 10^0.5^, *s* = 5.

**MOVIE S7.**
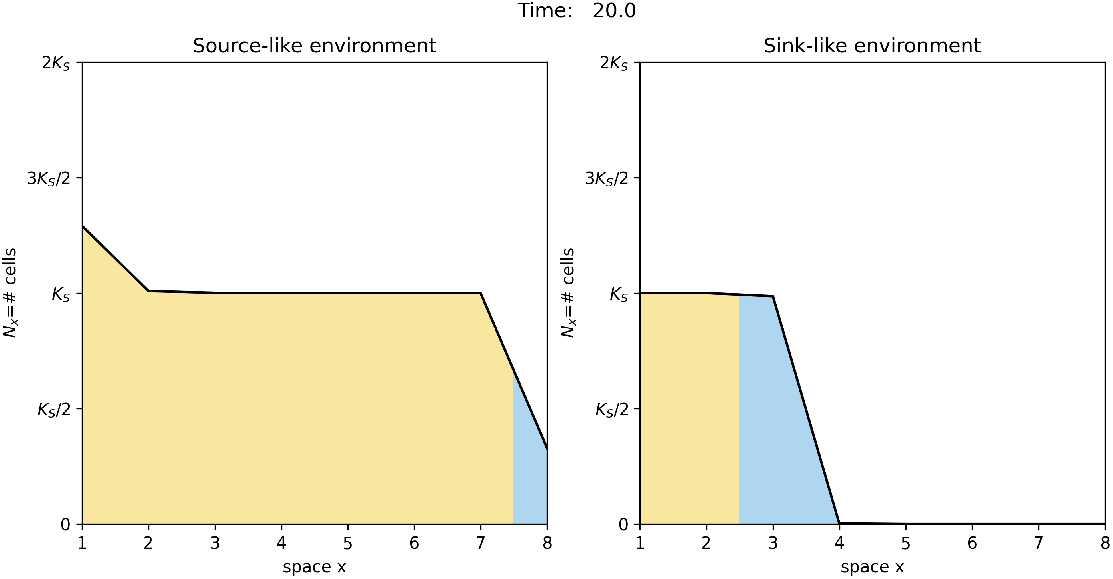
Effect of density-dependent motility. Simulation of a bacterial population evolving antibiotic in the staircase model with slow-to-fast density-dependent motility. If the environment is source-like (left), the faster high-density population bulk (yellow) expands its range by pulling the slower low-density population front (blue). If the environment is sink-like (right), and the faster high-density motility phenotype moves above the critical motility, this expansion is halted. As a result of this reduced antibiotic exposure, the adaptation rate is decreased in the top row of Fig. 4b (sink-like environment) but not in the top row of Fig. S5 (source-like environment). Parameters used: *L* = 8, *K* = 10^5^, *r* = 1, *δ* = 0.1 (left) and *δ* = 0.3 (right), *μ_f_* = 10^-7^, *μ_b_* = 10^-4^, *ν_L_*. = 10^-3^, *ν_H_* = 10^0.5^, *S* = 10^4^.

## Footnotes

1 Other fixed points are at play in the general staircase model but are irrelevant for the dynamics. The system of equations for the fixed points 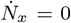 can be manipulated into a polynomial equation of degree 2^*R*^ by repeated substitution and elimination of variables. Therefore, there are at most 2^*R*^ fixed points by the Fundamental Theorem of Algebra. Indeed, when *ν* = 0, the equations decouple and there are two choices for 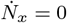 in *x* ≤ *R* (*N_x_* = 0 or *N_x_* = *K*(1 – *δ/r*)) and a single choice of 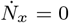 in *x* > *R* (*N_x_* = 0), leading to 2^*R*^ fixed points. Importantly, this includes the unstable trivial fixed point (*N_x_* = 0), the stable non-trivial fixed point (*N_x_* = *K*(1 – *δ/r*) in *x* ≤ *R, N_x_* = 0 in *x* > *R*), and other 2^*R*^ – 2 unstable irrelevant fixed points. As *ν* increases, the non-negative (*N_x_* ≥ 0) and non-positive (*N_x_* ≤ 0) quadrants become forward invariant with the trivial fixed being the only allowed fixed point on the boundary of these quadrants. The other quadrants are called irrelevant. Indeed, the irrelevant fixed points are unstable in directions perpendicular to the boundary of the positive quadrant and are therefore pushed into irrelevant quadrants as *ν* increases from 0. Moreover, the irrelevant fixed points move on smooth trajectories as *ν* increases and cannot smoothly enter into either the positive or negative quadrants, as this would require a bifurcation with the trivial fixed point and discontinuous d*N_x_*/d*ν*. Numerical simulations suggest that the irrelevant fixed points bifurcate to annihilate each other, and never bifurcate to appear again in some of the quadrants. One can define new coordinates: *N* = (*N*_1_ + … + *N_L_*)/*L* along the diagonal *N*_1_ = … = *N*_L_ and other coordinates for directions perpendicular to this diagonal. As *ν* → ∞, the dynamics in directions perpendicular to the diagonal quickly pushes the system close to the diagonal. On the diagonal a slower dynamics occurs, governed by 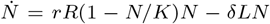. Therefore, only the trivial and non-trivial fixed points are preserved as *ν* → ∞, while the other fixed points must annihilate through bifurcations with each other. This explanation justifies why we have restricted our attention to the trivial and non-trivial fixed points only, which are the only fixed points to affect the dynamics in the physically relevant non-negative quadrant.

2 This definition coincides with the definition of convexity for functions ℝ → ℝ when the points on the graph, (1, *N*_1_),…, (*L*, *N_L_*), are joined by line segments.

## REFERENCES

[1] Adler, J. Chemotaxis in bacteria. Science 153, 708–716 (1966).

[2] Berg, H. C. E. coli in Motion (Springer-Verlag New York, 2004).

[3] Gude, S. et al. Bacterial coexistence driven by motility and spatial competition. Nature 578, 588–592 (2020).

[4] Josenhans, C. & Suerbaum, S. The role of motility as a virulence factor in bacteria. International Journal of Medical Microbiology 291, 605–614 (2002).

[5] Ottemann, K. M. & Miller, J. F. Roles for motility in bacterial–host interactions. Molecular Microbiology 24, 1109–1117 (1997).

[6] Stocker, R., Seymour, J., Samadani, A., Hunt, D. & Polz, M. Rapid chemotactic response enables marine bacteria to exploit ephemeral microscale nutrient patches. Proceedings of the National Academy of Sciences of the United States of America 105, 4209–14 (2008).

[7] Yawata, Y. et al. Competition–dispersal tradeoff ecologically differentiates recently speciated marine bacterioplankton populations. Proceedings of the National Academy of Sciences 111, 5622–5627 (2014).

[8] Davies, S. C., Fowler, T., Watson, J., Livermore, D. M. & Walker, D. Annual report of the chief medical officer: infection and the rise of antimicrobial resistance. The Lancet 381, 1606–1609 (2013).

[9] Zhang, Q. et al. Acceleration of emergence of bacterial antibiotic resistance in connected microenvironments. Science 333, 1764–1767 (2011).

[10] Baym, M., Stone, L. K. & Kishony, R. Multidrug evolutionary strategies to reverse antibiotic resistance. Science 351, aad3292 (2016).

[11] Hol, F. J., Hubert, B., Dekker, C. & Keymer, J. E. Density-dependent adaptive resistance allows swimming bacteria to colonize an antibiotic gradient. The ISME journal 10, 30–38 (2016).

[12] Kearns, D. B. A field guide to bacterial swarming motility. Nature Reviews Microbiology 8, 634–644 (2010).

[13] Lai, S., Tremblay, J. & Déziel, E. Lai s, tremblay j, deziel e.. swarming motility: a multicellular behaviour conferring antimicrobial resistance. environ microbiol 11: 126-136. Environmental microbiology 11, 126–36 (2008).

[14] Butler, M. T., Wang, Q. & Harshey, R. M. Cell density and mobility protect swarming bacteria against antibiotics. Proceedings of the National Academy of Sciences 107, 3776–3781 (2010).

[15] Bhattacharyya, S., Walker, D. M. & Harshey, R. M. Dead cells release a ‘necrosignal’that activates antibiotic survival pathways in bacterial swarms. Nature communications 11, 1–12 (2020).

[16] Coleman, S. R., Blimkie, T., Falsafi, R. & Hancock, R. E. Multidrug adaptive resistance of pseudomonas aeruginosa swarming cells. Antimicrobial agents and chemotherapy 64, e01999–19 (2020).

[17] Kim, W., Killam, T., Sood, V. & Surette, M. G. Swarm-cell differentiation in salmonella enterica serovar typhimurium results in elevated resistance to multiple antibiotics. Journal of bacteriology 185, 3111–3117 (2003).

[18] Partridge, J. D., Ariel, G., Schvartz, O., Harshey, R. M. & Be’er, A. The 3d architecture of a bacterial swarm has implications for antibiotic tolerance. Scientific reports 8, 1–11 (2018).

[19] Overhage, J., Bains, M., Brazas, M. D. & Hancock, R. E. Swarming of pseudomonas aeruginosa is a complex adaptation leading to increased production of virulence factors and antibiotic resistance. Journal of bacteriology 190, 2671–9 (2008).

[20] Roth, D. et al. Identification and characterization of a highly motile and antibiotic refractory subpopulation involved in the expansion of swarming colonies of p aenibacillus vortex. Environmental microbiology 15, 2532–2544 (2013).

[21] Turnbull, A. L. & Surette, M. G. L-cysteine is required for induced antibiotic resistance in actively swarming salmonella enterica serovar typhimurium. Microbiology 154, 3410–3419 (2008).

[22] Oliveira, N. M. et al. Suicidal chemotaxis in bacteria. Nature Communications Accepted (2022).

[23] Liu, Y., Kyle, S. & Straight, P. D. Antibiotic stimulation of a bacillus subtilis migratory response. Msphere 3, e00586–17 (2018).

[24] Bru, J.-L. et al. Pqs produced by the pseudomonas aeruginosa stress response repels swarms away from bacteriophage and antibiotics. Journal of bacteriology 201, e00383–19 (2019).

[25] Hermsen, R. & Hwa, T. Sources and sinks: a stochastic model of evolution in heterogeneous environments. Physical review letters 105, 248104 (2010).

[26] Hermsen, R., Deris, J. & Hwa, T. On the rapidity of antibiotic resistance evolution facilitated by a concentration gradient. Proceedings of the National Academy of Sciences 109, 10775–10780 (2012).

[27] Greulich, P., Waclaw, B. & Allen, R. J. Mutational pathway determines whether drug gradients accelerate evolution of drug-resistant cells. Physical review letters 109, 088101 (2012).

[28] Hermsen, R. The adaptation rate of a quantitative trait in an environmental gradient. Physical Biology 13, 065003 (2016).

[29] Gralka, M., Fusco, D., Martis, S. & Hallatschek, O. Convection shapes the trade-off between antibiotic efficacy and the selection for resistance in spatial gradients. Physical Biology 14, 045011 (2017).

[30] De Jong, M. G. & Wood, K. B. Tuning spatial profiles of selection pressure to modulate the evolution of drug resistance. Phys. Rev. Lett. 120, 238102 (2018).

[31] Steel, H. & Papachristodoulou, A. The effect of spatiotemporal antibiotic inhomogeneities on the evolution of resistance. Journal of theoretical biology 486, 110077 (2019).

[32] Pulliam, H. R. Sources, sinks, and population regulation. The American Naturalist 132, 652–661 (1988).

[33] Oliveira, N. M., Foster, K. R. & Durham, W. M. Singlecell twitching chemotaxis in developing biofilms. Proceedings of the National Academy of Sciences 113, 6532–6537 (2016).

[34] Sousa, A. M., Machado, I. & Pereira, M. O. Phenotypic switching: an opportunity to bacteria thrive. Science Against Microbial Pathogens: Communicating Current Research and Technological Advances 1(2012).

[35] Miller, M. B. & Bassler, B. L. Quorum sensing in bacteria. Annual Reviews in Microbiology 55, 165–199 (2001).

[36] Oliveira, N. M. et al. Biofilm formation as a response to ecological competition. PLoS biology 13, e1002191 (2015).

[37] Kearns, D. B. & Losick, R. Swarming motility in undomesticated bacillus subtilis. Molecular microbiology 49, 581–590 (2003).

[38] Kepler, T. & Perelson, A. Drug concentration heterogeneity facilitates the evolution of drug resistance. Proceedings of the National Academy of Sciences of the United States of America 95 20, 11514–9 (1998).

[39] Fu, F., Nowak, M. A. & Bonhoeffer, S. Spatial heterogeneity in drug concentrations can facilitate the emergence of resistance to cancer therapy. PLOS Computational Biology 11, 1–22 (2015).

[40] Lambert, G. et al. An analogy between the evolution of drug resistance in bacterial communities and malignant tissues. Nature Reviews Cancer 11, 375–382 (2011).

[41] Wu, A. et al. Cell motility and drug gradients in the emergence of resistance to chemotherapy. Proceedings of the National Academy of Sciences 110, 16103–16108 (2013).

[42] Kirkpatrick, M. & Barton, N. Evolution of a species’ range. The American Naturalist 150, 1–23 (1997).

[43] Lenormand, T. Gene flow and the limits to natural selection. Trends in Ecology & Evolution 17, 183–189 (2002).

[44] Angert, A., Bontrager, M. & Ågren, J. What do we really know about adaptation at range edges? Annual Review of Ecology, Evolution, and Systematics 51, 341–361 (2020).

[45] Polechová, J. & Barton, N. H. Limits to adaptation along environmental gradients. Proceedings of the National Academy of Sciences 112, 6401–6406 (2015).

[46] Bulmer, M. Multiple niche polymorphism. The American Naturalist 106, 254–257 (1972).

[47] Holt, R. D. & Gomulkiewicz, R. How does immigration influence local adaptation? a reexamination of a familiar paradigm. The American Naturalist 149, 563–572 (1997).

[48] Nagylaki, T. Conditions for the existence of clines. Genetics 80, 595–615 (1975).

[49] Niehus, R., Mitri, S., Fletcher, A. G. & Foster, K. R. Migration and horizontal gene transfer divide microbial genomes into multiple niches. Nature communications 6, 1–9 (2015).

[50] Cohan, F. M. The effects of rare but promiscuous genetic exchange on evolutionary divergence in prokaryotes. The American Naturalist 143, 965–986 (1994).

[51] Cohan, F. M. Does recombination constrain neutral divergence among bacterial taxa? Evolution 49, 164–175 (1995).

[52] Shapiro, B. J., David, L. A., Friedman, J. & Alm, E. J. Looking for darwin’s footprints in the microbial world. Trends in microbiology 17, 196–204 (2009).

[53] Gevers, D. et al. Re-evaluating prokaryotic species. Nature Reviews Microbiology 3, 733–739 (2005).

[54] Levin, B. R. Periodic selection, infectious gene exchange and the genetic structure of e. coli populations. Genetics 99, 1–23 (1981).

[55] Haas, P. A., Oliveira, N. M. & Goldstein, R. E. Subpopulations and stability in microbial communities. Physical Review Research 2, 022036 (2020).

[56] Gibson, M. A. & Bruck, J. Efficient exact stochastic simulation of chemical systems with many species and many channels. The journal of physical chemistry A 104, 1876–1889 (2000).

[57] Gillespie, D. T. Exact stochastic simulation of coupled chemical reactions. The journal of physical chemistry 81, 2340–2361 (1977).

## REFERENCES

S1. Bulmer, M. Multiple niche polymorphism. The American Naturalist 106, 254–257 (1972).

S2. Holt, R. D. & Gomulkiewicz, R. How does immigration influence local adaptation? a reexamination of a familiar paradigm. The American Naturalist 149, 563–572 (1997).

S3. Lenormand, T. Gene flow and the limits to natural selection. Trends in Ecology & Evolution 17, 183–189 (2002).

S4. Hastings, A. Dynamics of a single species in a spatially varying environment: the stabilizing role of high dispersal rates. Journal of mathematical biology 16, 49–55 (1982).

S5. Lipsitch, M. The rise and fall of antimicrobial resistance. Trends in microbiology 9, 438–444 (2001).

S6. Andersson, D. I. The biological cost of mutational antibiotic resistance: any practical conclusions? Current opinion in microbiology 9, 461–465 (2006).

S7. Hermsen, R., Deris, J. & Hwa, T. On the rapidity of antibiotic resistance evolution facilitated by a concentration gradient. Proceedings of the National Academy of Sciences 109, 10775–10780 (2012).

S8. Hermsen, R. & Hwa, T. Sources and sinks: a stochastic model of evolution in heterogeneous environments. Physical review letters 105, 248104 (2010).

S9. Oliveira, N. M. et al. Suicidal chemotaxis in bacteria. Nature Communications. Accepted (2022).

S10. Bru, J.-L. et al. Pqs produced by the pseudomonas aeruginosa stress response repels swarms away from bacteriophage and antibiotics. Journal of bacteriology 201, e00383–19 (2019).

S11. Von Wintersdorff, C. J. et al. Dissemination of antimicrobial resistance in microbial ecosystems through horizontal gene transfer. Frontiers in microbiology 173 (2016).

S12. Niehus, R., Mitri, S., Fletcher, A. G. & Foster, K. R. Migration and horizontal gene transfer divide microbial genomes into multiple niches. Nature communications 6, 1–9 (2015).

S13. Cohan, F. M. The effects of rare but promiscuous genetic exchange on evolutionary divergence in prokaryotes. The American Naturalist 143, 965–986 (1994).

S14. Cohan, F. M. Does recombination constrain neutral divergence among bacterial taxa? Evolution 49, 164–175 (1995).

S15. Shapiro, B. J., David, L. A., Friedman, J. & Alm, E. J. Looking for darwin’s footprints in the microbial world. Trends in microbiology 17, 196–204 (2009).

S16. Gevers, D. et al. Re-evaluating prokaryotic species. Nature Reviews Microbiology 3, 733–739 (2005).

S17. Levin, B. R. Periodic selection, infectious gene exchange and the genetic structure of e. coli populations. Genetics 99, 1–23 (1981).

S18. Hastings, A. Can spatial variation alone lead to selection for dispersal? Theoretical Population Biology 24, 244–251 (1983).

S19. Barrett, P., Hunter, J., Miller, J. T., Hsu, J.-C. & Greenfield, P. matplotlib–a portable python plotting package. Astronomical data analysis software and systems XIV 347, 91 (2005).

